# Kin selection as a modulator of human handedness: sex-specific, parental and parent-of-origin effects

**DOI:** 10.1101/2023.08.08.552537

**Authors:** Bing Dong, Silvia Paracchini, Andy Gardner

**Affiliations:** School of Biology, University of St Andrews, Dyers Brae, St Andrews KY16 9TH, UK; School of Medicine, University of St Andrews, North Haugh, St Andrews KY16 9TF, UK

**Keywords:** evolution, game theory, lateralization, inclusive fitness, genomic imprinting, neurodevelopmental disorders

## Abstract

The frequency of left-handedness in humans is ∼10% worldwide and slightly higher in males than females. Twin and family studies estimate the heritability of human handedness at around 25%. The low but substantial frequency of left-handedness has been suggested to imply negative frequency-dependent selection, e.g. owing to a “surprise” advantage of left-handers in combat against opponents more used to fighting right-handers. Because such game-theoretic hypotheses involve social interaction, here, we perform an analysis of the evolution of handedness based on kin-selection, which is understood to play a major role in the evolution of social behaviour generally. We show that: (1) relatedness modulates the balance of right-handedness versus left-handedness, according to whether left-handedness is marginally selfish versus marginally altruistic; (2) sex differences in relatedness to social partners may drive sex differences in handedness; (3) differential relatedness of parents and offspring may generate parent-offspring conflict and sexual conflict leading to the evolution of maternal and paternal genetic effects in relation to handedness; and (4) differential relatedness of maternal-origin versus paternal-origin genes may generate intragenomic conflict leading to the evolution of parent-of-origin-specific gene effects—such as “genomic imprinting”—and associated maladaptation.

## 1 Introduction

Most humans show a preference for—or a difference in proficiency of—one hand over the other for a range of tasks (McManus 2019, Papadatou-Pastou et al. 2020). The frequency of left-handedness in humans is estimated at 10.6%, fairly stably across regions and populations, and is somewhat higher in males (11.6%) than in females (9.5%) (Papadatou-Pastou et al. 2020). Twin studies (Medland et al. 2009) and family studies (Lien et al. 2015) reveal that handedness is heritable, with additive genetic effects appearing to explain around 25% of the variance (Medland et al. 2009, Somers et al. 2015). Genome-wide association studies (GWAS) have identified 41 loci influencing handedness and explaining around 6% of the heritability (Cuellar-Partida et al. 2021). A recent whole exome sequencing (WES) study in the UK Biobank has suggested an association between mutations in the *TUBB4B* gene and left-handedness and estimated that the heritability of left-handedness due to rare coding variants to be 0.91% (Schijven et al. 2023). Left-handedness is linked to some psychiatric disorders, such as autism spectrum disorders (ASD) (Markou et al. 2017), schizophrenia (Hirnstein & Hugdahl 2014) and dyslexia (Abbondanza et al. 2023), and there is an overlap among genes underlying these conditions, brain asymmetries and handedness (Papadatou-Pastou et al. 2020).

Although many taxa exhibit some form of lateralization (Rogers 1980, Vallortigara & Bisazza 2002, Ocklenburg & Güntürkün 2012, Ströckens et al. 2013, Versace & Vallortigara 2015), of which handedness is just one form, these typically involve roughly equal numbers of left-sided and right-sided individuals, and so the strong population bias towards right-handers is peculiarly human (Frayer et al. 2012, Ströckens et al. 2013, Caspar et al. 2022).

The left hemisphere dominance for language processing may have an important role in explaining the rightward bias of handedness, (Levy & Nagylaki 1972). Indeed, atypical language hemispheric lateralization is associated with the degree of left-handedness (Knecht et al. 2000, Mazoyer et al. 2014).

The stability of the ∼10% incidence of left-handedness in human populations through time (Coren & Porac 1977, McManus 1991, Frayer et al. 2012) and across regions (Papadatou-Pastou et al. 2020) has given rise to the suggestion that left-handedness is maintained by negative frequency-dependent selection, and this has motivated the development of a number of evolutionary game-theoretic hypotheses to explain the phenomenon (Raymond et al. 1996, Ghirlanda et al. 2009, Abrams & Panaggio 2012, Schaafsma et al. 2012, Faurie & Raymond 2013). As an illustrative example, the “combat hypothesis” suggests that left-handers suffer a basic disadvantage (Schaafsma et al. 2012, Zickert et al. 2018, Papadatou-Pastou et al. 2020)—e.g. perhaps owing to disruption of typical brain lateralisation—such that natural selection has resulted in them being in the minority, yet also enjoy a compensating advantage when they are sufficiently rare, owing to the element of surprise in combat and similar competitive interactions (Gibbons 1993, Raymond et al. 1996, Faurie & Raymond 2013). The combat hypothesis is in line with the higher incidences of left-handers among elite athletes in interactive sports, such as tennis, fencing and baseball (Wood & Aggleton 1989, Raymond et al. 1996, Loffing 2017).

These game-theoretic hypotheses centre upon social interaction, and kin selection—the part of natural selection that arises when individuals have an impact on the fitness of their genetically related social partners—plays a major role in the evolution of social behaviour across the tree of life (Hamilton 1964, Frank 1998, West et al. 2007a). In addition to influencing the overall incidence of traits within and across populations (Turner & Chao 1999, Queller et al. 2003, Sachs et al. 2004, West et al. 2007a), patterns of genetic relatedness can explain differences in trait levels between different individuals—such as sex differences (West et al. 2007a, Leedale et al. 2018)—and also modulate evolutionary conflicts of interest within families and even within individual genomes—resulting in the evolution of parental genetic effects (Wolf et al. 1998, Richardson et al. 2004, Kuijper & Johnstone 2016, Kuijper & Johnstone 2019) and parent-of-origin effects e.g. genomic imprinting (Haig 2000, 2002, Wilkins & Úbeda 2011, Crespi 2020). However, the scope for a modulating role of kin selection in the evolution of human handedness remains to be investigated.

Here we undertake a theoretical investigation of how relatedness and kin selection shape the biology of human handedness. First, we show that, at evolutionary equilibrium, left-handedness may be classified either as a “selfish” or an “altruistic” trait, depending on its fitness consequences for the individual and for her social partners, and that the direction of the modulating effect of genetic relatedness depends on which of these two situations applies. Second, we explore how demographic processes such as dispersal modulate the population level of left-handedness at evolutionary equilibrium, via their impact on the degree of genetic relatedness between social partners. Third, we investigate the consequences of sex-biased dispersal, and associated sex differences in an individual’s relatedness to social partners, for the evolution of sex differences in left-handedness. Fourth, we determine the consequences of extending genetic control of handedness to the individual’s parents, resulting in parent-offspring conflict and sexual conflict and the evolution of parental genetic effects in relation to human handedness. Fifth, we descend to the level of individual genes and investigate the scope for intragenomic conflict between maternal-origin versus paternal-origin genes and the resulting evolution of parent-of-origin effects—including genomic imprinting—in relation to human handedness. For the purpose of illustration and concreteness, in each case we derive quantitative predictions for explicit “within-group combat” and “between-group combat” game-theoretic scenarios, but more generally our analysis applies to any scenario in which an individual’s handedness has an impact upon their own reproductive success and that of genetically related social partners. Our model allows for handedness to be a highly polygenic trait and, although for ease of conceptualization we will often refer to handedness in a binary way, our analysis also readily accommodates a spectrum of handedness.

## 2 Results

### (a)#Kin selection and human handedness

Natural selection adapts individuals as if for the purpose of passing on their alleles to future generations (Hamilton 1964, Grafen 2006, West & Gardner 2013). There are two basic routes through which individuals can accomplish this: first, by promoting their own reproductive success (direct fitness); and, second, by promoting the reproductive success of their genetic relatives, who tend to share alleles in common (indirect fitness) (Hamilton 1964). According to Hamilton’s (1963, 1964, 1970) rule, a behaviour that incurs a fitness cost (*c*) for the actor can nevertheless be favoured by natural selection if it provides a sufficiently large fitness benefit (*b*) to a sufficiently closely related (*r*) social partner (specifically, such that -*c* + *r b* > 0). More generally, we can define four types of social behaviour, according to the sign of the fitness effects: traits incurring a cost for the actor and yielding a benefit for the recipient (*c* > 0 and *b* > 0) are “altruistic”; traits yielding a benefit for the actor and incurring a cost for the recipient (*c* < 0 and *b* < 0) are “selfish”; traits yielding a benefit for both parties (*c* < 0 and *b* > 0) are “mutually beneficial”; and traits incurring a cost for both parties (*c* > 0 and *b* < 0) are “spiteful” (Hamilton 1964, 1970, West et al. 2007b).

At evolutionary equilibrium, where natural selection favours neither an increase nor a decrease in the trait (*b r* - *c* = 0), then so long as relatedness is positive (*r* > 0) the trait must either be marginally altruistic (*c* > 0 and *b* > 0) or marginally selfish (*c* < 0 and *b* < 0) (Hitchcock et al. 2019). Accordingly, if natural selection acts in a negative frequency- dependent way in relation to human handedness—as suggested by the game-theoretic models (Ghirlanda & Vallortigara 2004, Ghirlanda et al. 2009, Abrams & Panaggio 2012)—such that it favours an increase in the incidence of left-handedness when this has dropped below a threshold level and favours a decrease in left-handedness when it has exceeded the threshold, then evolutionary equilibrium is attained when the incidence of left-handedness is at the threshold, and at this point left-handedness is either marginally altruistic or marginally selfish. If left-handedness is marginally altruistic then a higher degree of genetic relatedness between actor and recipient is expected to be associated with a higher incidence of left- handedness at the evolutionary equilibrium, whereas if left-handedness is marginally selfish then a higher degree of genetic relatedness is expected to be associated with a lower incidence of left-handedness.

Taking the combat hypothesis as a purely illustrative example, if we imagine that combat occurs mainly within human groups—between somewhat-related individuals and over reproductive resources—then the indirect-fitness consequences of enjoying a surprise advantage in combat owing to left-handedness are expected to be negative (because the opponent, who loses out, is a genetic relative), and hence at equilibrium this is expected to be exactly balanced by a direct-fitness benefit (owing to improved success in combat outweighing the basic disadvantage of left-handedness), such that left-handedness is a marginally selfish trait. In this scenario, a higher level of relatedness is expected to be associated with a lower incidence of left-handedness (Figure 1*a*). Alternatively, if combat mainly occurs between non-relatives in a group-warfare context in which success in combat is associated with a positive indirect-fitness effect owing to the benefits that accrue to the individual’s genetically related group mates, then at equilibrium this is expected to be exactly balanced by a direct-fitness cost (owing to the basic disadvantage of left-handedness failing to outweigh the improved success in combat), such that left-handedness is a marginally altruistic trait. In this scenario, a higher level of relatedness is expected to be associated with a higher incidence of left-handedness (Figure 1*a*).

**Figure 1.**
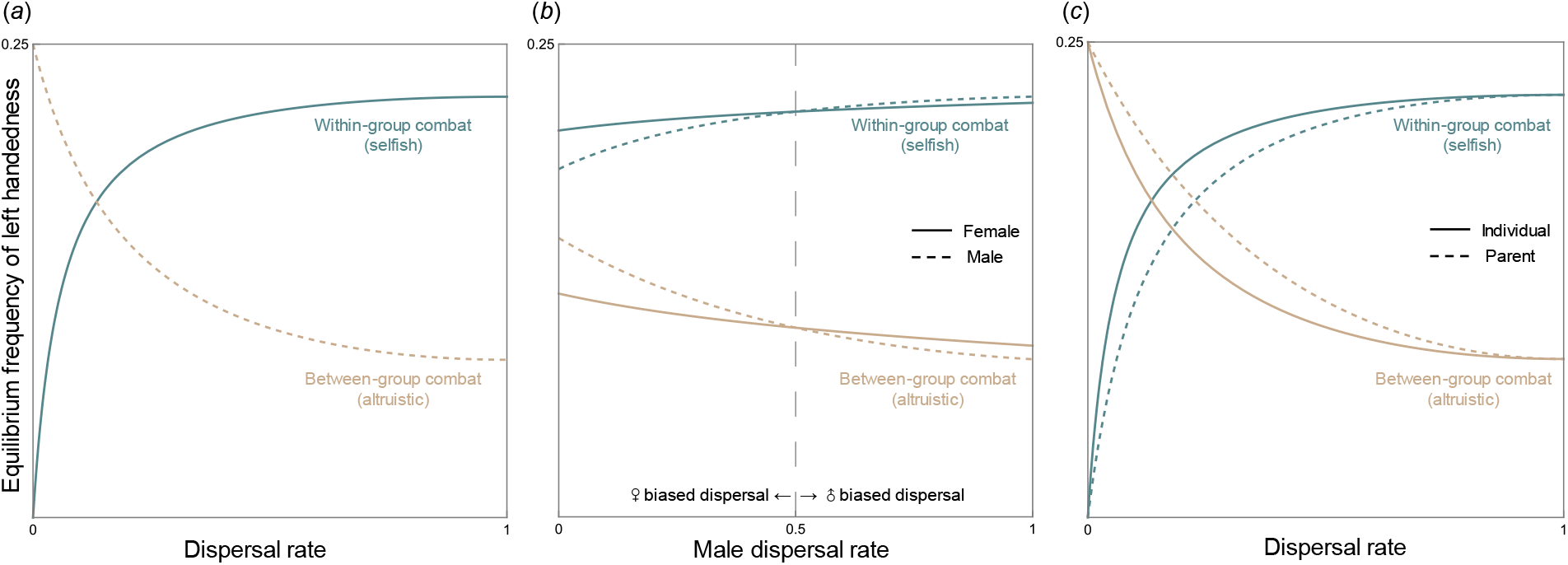
Level of left-handedness can be mediated by demographic features such as dispersal, as higher dispersal reduces relatedness between social partners. (*a*) Level of left- handedness is mediated by dispersal in the context of within-group combat (left-handedness is selfish) versus between-group combat (left-handedness is altruistic). (*b*) Sex effects in left- handedness: level of left-handedness can be mediated by sex and dispersal pattern (female/male biased dispersal). (*c*) Parental genetic effects in left-handedness: level of left- handedness can be mediated by dispersal, and further result in parent-offspring disagreement on handedness. Here, we set female dispersal rate *m*_f_ to be 0.5, the relative importance of combat in relation to other types of competitions for females *b*_f_ and males *b*_m_ both to be 1, and the number of individuals each sex born in the same patch *n* to be 5 (parameter details see Supplementary Material §§S1.3).

Relatedness will usually depend on the ecology and demography of the population, and so the above insights also yield predictions as to how population processes relate to the evolutionarily favoured incidence of left-handedness. As a concrete example, we consider the rate of dispersal. If individuals have a higher tendency to disperse away from their place of origin and pursue reproductive opportunities within other groups, then this is expected to result in lower relatedness between group mates. Accordingly, if left-handedness is marginally selfish—as, for example, in the within-group combat scenario—then as the rate of dispersal increases, the evolutionarily favoured level of left-handedness is expected to increase (Figure 1*a*). And, in contrast, if left-handedness is marginally altruistic—as, for example, in the between-group combat scenario—then as the rate of dispersal increases, the level of left-handedness is expected to decrease (Figure 1*a*). These predictions relate to contemporary and/or historical between-population comparisons and also, potentially, to the dynamics of handedness within a single population across evolutionary timescales in responses to demographic change (see Discussion).

### **(b)** Sex differences in human handedness

Above, we have shown that the average genetic relatedness between social partners—and the population processes that modulate this—is expected to influence the evolutionarily favoured incidence of left-handedness at a population level. Similarly, inter-individual differences in relatedness to one’s social partners—and the population processes responsible for such variation—are expected to drive differences in levels of left-handedness among different subdivisions of the population. In particular, sex-specific demographic processes—such as sex-biased dispersal—may result in a sex difference in the relatedness of social partners, which may favour a sex difference in the incidence of left-handedness. For example, all else being equal, female-biased dispersal is expected to result in relatedness between social partners being lower for woman than for men; hence, all else being equal, a higher level of left-handedness would be favoured among women than among men if left-handedness is marginally selfish (such as in the within-group combat scenario) and a higher level of left- handedness would be favoured among men than among women if left-handedness is marginally altruistic (such as in the between-group combat scenario) (Figure 1*b*). The opposite pattern is expected under male-biased dispersal (Figure 1*b*).

In addition to differences in relatedness, the sexes might also differ with respect to the fitness consequences—that is, the benefits and costs—associated with left-handedness. Such fitness differences would also be expected to modulate sex differences in incidence of left- handedness. For example, if the frequency-dependent advantage of left-handedness when rare applies more strongly to men than to women—as would be expected in the combat scenarios if men engage in combat more frequently than do women (Divale & Harris 1976, Micheletti et al. 2018) and/or if men have more to gain from winning in combat in terms of enhanced reproductive success (Gibbons 1993)—then, all else being equal, the incidence of left-handedness is expected to be higher among men than among women. More generally, these sex-difference results concern adaptive evolution, and are based upon considerations of female versus male fitness optima. Accordingly, they neglect non-adaptive sex differences arising, for example, from a greater vulnerability of males to developmental perturbation away from a default phenotype, which has been reported in disorders including ASD (Antaki et al. 2022), and this could offer alternative explanations for the higher incidence of left-handedness among males (see Discussion).

### **(c)** Parental genetic effects in human handedness

Above, we have shown how the evolutionarily favoured level of left-handedness may be modulated by the valuation that individuals place upon the reproductive success of social partners relative to their own reproductive success. This assumes that an individual’s own genotype controls the handedness phenotype and if, instead, the handedness phenotypes were controlled by the parental genotype—i.e. a “parental genetic effect”; (Trivers 1972, Trivers 1974, Wilson 1980, Wolf et al. 1998, Badyaev & Uller 2009, Hwang et al. 2020)—then we might expect the evolutionarily favoured incidence of left-handedness to reflect the relatedness valuations made by the individual’s parents. More generally, if an individual’s predisposition to left-handedness is modulated in part by the individual’s own genotype and also in part by the genotypes of the individual’s parents then we might expect an evolutionary conflict of interests—and associated evolutionary arms race—between parent and offspring (Trivers 1974), and between the parents themselves (Trivers 1972), as each party is favoured to move the handedness phenotype closer to their own fitness optimum.

If an individual’s handedness phenotype represents a trade-off between the individual’s own reproductive success and the reproductive success of the individual’s group mates, then in general terms we expect the individual’s parents to favour a balance that is relatively in support of the group mates’ reproductive interests and the individual to favour a balance that is relatively in support of their own reproductive interests, so long as there is relatedness among group mates (see Supplementary Material §§S1.7&S2.5 for details). This owes to individuals being genetically identical to themselves and only somewhat genetically related to their offspring. Accordingly, if left-handedness is a marginally selfish trait (as in the illustrative within-group combat scenario) then we expect parents to favour a lower predisposition for left-handedness in their offspring than their offspring would themselves favour, and if left-handedness is a marginally altruistic trait (as in the illustrative between-group combat scenario) then we expect parents to favour a higher predisposition for left-handedness in their offspring than their offspring would themselves favour (Figure 1*c*).

Moreover, although both parents are equally related to their offspring they may be differentially related to their offspring’s social partners, so that mothers and fathers may favour different dispositions for left-handedness among their offspring. For example, under female-biased dispersal, mothers are expected to be less related to their offspring’s social partners than are fathers, and hence more inclined to their offspring having a disposition for left-handedness if this is a marginally selfish trait (as in the illustrative within-group combat scenario) and less inclined to their offspring having a disposition for left-handedness if this is a marginally altruistic trait (as in the illustrative between-group combat scenario) and the opposite set of outcomes is expected under male-biased dispersal (Figure S1). Accordingly, considerations of patterns of relatedness and concomitant kin selection yields predictions as to parental genetic effects—including maternal genetic effects and paternal genetic effects—working at cross purposes with the individual’s own genome, as well as with each other, in relation to the individual’s handedness phenotype.

### **(d)** Parent-of-origin effects in human handedness

Above, we have shown that sex-specific demography—such as sex-biased dispersal—may generate differences in the relatedness valuations made by mothers and fathers regarding the reproductive success of their offspring versus their offspring’s social partners, resulting in the evolution of parental genetic effects in relation to handedness. Similarly, this relatedness asymmetry can also extend into the offspring’s own genome and ignite an evolutionary conflict of interests between the individuals own maternal-origin versus paternal-origin genes. Such intragenomic conflict in relation to other social traits has been suggested to drive in the evolution of parent-of-origin specific genetic effects, including genomic imprinting (Haig 2002; Gardner & Úbeda 2017)—and induce vulnerability to a number of associated developmental disorders, e.g. Silver-Russell syndrome (SRS) and Beckwith-Wiedemann syndrome (BWS) (Crespi 2011, Wilkins & Úbeda 2011, Crespi 2020).

For example, if left-handedness is marginally selfish (such as in the within-group combat scenario) then under female-biased dispersal the relatedness between social partners through maternal-origin genes—all else being equal—will be lower than the relatedness through paternal-origin genes, and hence maternal-origin genes are expected to favour a higher level of left-handedness than are paternal-origin genes (Figure 2); whereas under male-biased dispersal relatedness will be higher through maternal-origin genes than through paternal-origin genes, and hence maternal-origin genes are expected to favour a lower level of left-handedness than are paternal-origin genes (Figure 2). Conversely, when left-handedness is marginally altruistic (such as in the between-group combat scenario) then under female-biased dispersal maternal-origin genes are expected to favour a lower level of left-handedness than are paternal-origin genes, whereas under male-biased dispersal maternal-origin genes are expected to favour a higher level of left-handedness than are paternal-origin genes (Figure 2).

**Figure 2.**
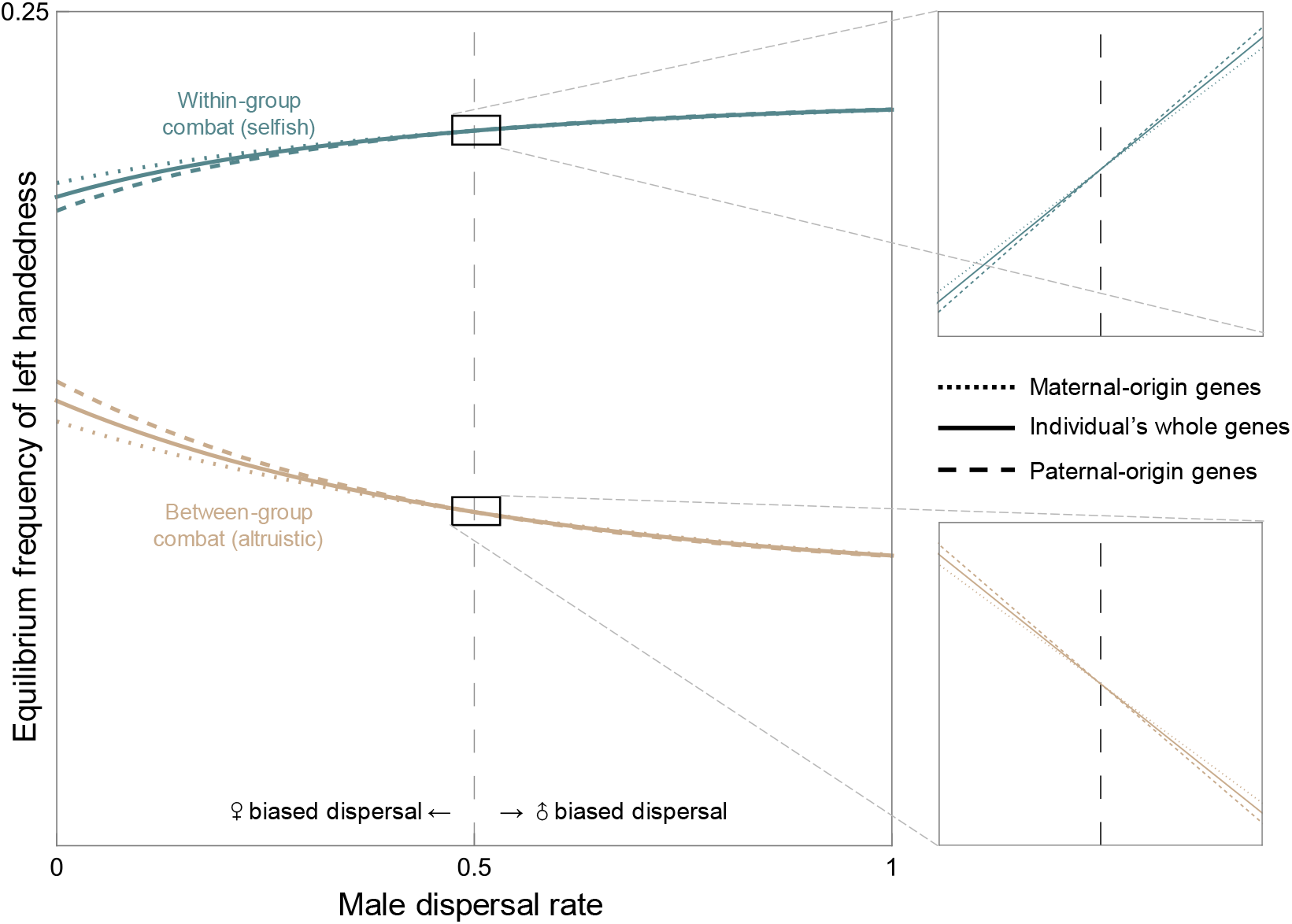
Parent-of-origin effects in left-handedness: level of left-handedness can be mediated by where the genes are inherited (from mother versus from father) effects and dispersal pattern (female/male biased dispersal) in the context of within-group combat (left- handedness is selfish) versus between-group combat (left-handedness is altruistic). Here, we set female dispersal rate *m*_f_ to be 0.5, the relative importance of combat in relation to other types of competitions for females *b*_f_ and males *b*_m_ both to be 1, and the number of individuals each sex born in the same patch *n* to be 5.

According to the kinship theory of genomic imprinting (Haig 2002), this form of intragenomic conflict will typically lead to one of the copies of the gene being silenced. Specifically, according to the “loudest voice prevails” principle (Haig 1996), the two copies of the gene at the affected locus are favoured to adjust their level of expression in opposite directions, such that the one favouring a higher level of left-handedness will act to increase the level of left-handedness while the one favouring a lower level of left-handedness will act to decrease the level of left-handedness, with perhaps no net change in the actual level of left-handedness, until the gene being favoured to decrease its expression falls silent, after which point the other gene will increase its expression to a level corresponding with its evolutionarily favoured level of left-handedness. At a locus for which an increase in gene expression results in an increase in the level of left-handedness—a “left-handedness promoter” locus—it is the gene that favours a higher level of left-handedness that is expected to remain expressed while the gene that favours a lower level of left-handedness is silenced, and at a locus for which an increase in gene expression results in a decrease in the level of left-handedness—a “left-handedness inhibitor” locus—it is the gene that favours a lower level of left-handedness that is expected to remain expressed while the gene that favours a higher level of left-handedness is silenced. Accordingly, the function of the gene product determines the direction of imprint.

For example, if left-handedness is marginally selfish (e.g. within-group combat), then under female-biased dispersal we expect left-handedness promoters to be maternally expressed and paternally silenced and left-handedness inhibitors to be maternally silenced and paternally expressed, and under male-biased dispersal left-handedness promoters are expected to be maternally silenced and paternally expressed and left-handedness inhibitors to be maternally expressed and paternally silenced; however if left-handedness is marginally altruistic (e.g. between-group combat), then under female-biased dispersal we expect left-handedness promoters to be maternally silenced and paternally expressed and left-handedness inhibitors to be maternally expressed and paternally silenced, and under male-biased dispersal left-handedness promoters are expected to be maternally expressed and paternally silenced and left-handedness inhibitors to be maternally silenced and paternally expressed (Figure 3).

**Figure 3.**
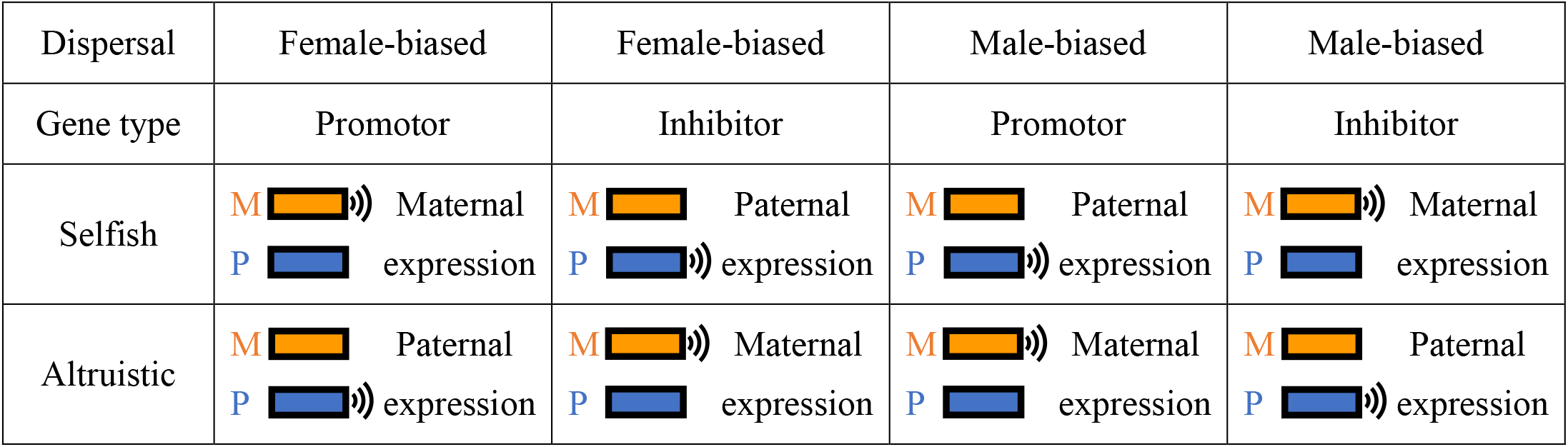
How dispersal type and gene type modulate the expression/imprinting pattern of maternal- versus paternal-origin genes in relation to the level of left-handedness, according to the kinship theory (see Supplementary Material Figure S3 for the phenotypic consequences of gene deletions, gene duplications, epimutations and uniparental disomies).

## 3 Discussion

Although game theoretic attempts to explain the evolutionary maintenance of a substantial minority of left-handed individuals in human population fundamentally hinge upon social interaction, and although kin selection is a fundamental driver of social evolution, the possible role for kin selection in modulating the evolution of human handedness has previously been neglected. We have shown how patterns of relatedness—and the demographic processes underpinning these—are expected to shape patterns of human handedness. Specifically, our kin-selection analyses reveal that: (1) genetic relatedness between social partners—modulated by population processes such as dispersal—is expected to influence the population level of left-handedness in a direction that depends upon whether left-handedness is marginally selfish (as in our illustrative within-group combat scenario) versus marginally altruistic (as in our illustrative between-group combat scenario); (2) sex-specific demography—such as sex-biased dispersal—can result in differences in sex differences in relatedness to one’s social partners, which may go some way to explaining sex differences in incidence of left-handedness; (3) differences in relatedness valuations made by different family members can ignite conflicts of interest between parents and offspring and between an individual’s mother and father over their handedness phenotype, driving the evolution of parental genetic effects; and (4) such relatedness differences may even ignite evolutionary conflicts of interest within the individual’s own genome, with maternal-origin and paternal-origin genes favouring different handedness phenotype, which is expected to drive the evolution of parent-of-origin effects—such as “genomic imprinting”—in relation to handedness.

Our analyses have shown that the degree of genetic relatedness between social partners whose reproductive success is modulated by each other’s handedness phenotypes is expected to modulate the evolutionary equilibrium frequency of left-handedness in the population, with higher relatedness being associated with a lower level of left-handedness when left-handedness tends to benefit the individual at the expense of social partners (selfishness) and a higher level of left-handedness when left-handedness tends to benefit social partners at the expense of the individual (altruism). The degree of relatedness is itself expected to depend on ecological and demographic parameters such as rate of dispersal, which higher dispersal of individuals tending to reduce the extent of genetic relatedness between social partners. At a comparative level, variation in ecological and demographic parameters between different human populations could potentially explain between-population differences in incidence of left-handedness, and variation in ecological and demographic parameters within a single human population over time might explain temporal differences in the incidence of left-handers, but only insofar as the variation in ecology and demography occurs over a relatively long timescale and the evolutionary fine-tuning of handedness occurs over a relatively short timescale. Our analysis offers little quantitative guidance as to the relevant timescales, but the population bias towards right-handedness does appear to have already been in place when hominin lineages diverged from the great apes around seven million years ago (Uomini & Ruck 2018, Papadatou-Pastou et al. 2020).

Our analysis also reveals that sex-specific selection can give rise to sex differences in handedness. We have shown how sex-specific demographies—such as sex-biased dispersal— may lead to sex differences in relatedness between social partners and hence sex-differences in the level of left-handedness favoured by females versus males. Whether humans have been characterised by sex-biased dispersal in our evolutionary past, and in which direction, remains a controversial topic: the traditional view is that human dispersal has been female- biased (Ember 1975), but evidence has also been marshalled in support of dispersal having been unbiased or mixed (Marlowe 2004). Our use of sex-biased dispersal is merely as an illustration, and the results extend more generally to any ecological and demographic factors that result in sex-differences in relatedness to one’s social partners—such as patterns of inbreeding (Wilkins & Haig 2003). In addition to relatedness, our analysis has emphasised that sex difference in left-handedness might also reflect sex differences in the costs and/or benefits of left-handedness. For example, men are generally understood to engage in—and to benefit from winning—combat more than do women (Divale & Harris 1976, Micheletti et al. 2018), which could explain a higher incidence in otherwise-costly left-handedness on account of a surprise advantage in combat settings. The higher incidence of left-handedness in males could also arise for non-adaptive reasons, such as sexually differential liability thresholds (Khramtsova et al. 2019, Merikangas & Almasy 2020), whereby the number of risk alleles required for an individual to exhibit a minority phenotype is greater for females than males, i.e. the female buffering effect.

Our analysis reveals the potential for parental genetic effects to occur in relation to left-handedness, such that alleles carried by a parent exert an influence on their offspring’s handedness phenotype, irrespective of whether the offspring carries the same alleles. These parental genetic effects are expected to arise evolutionarily as a consequence of parents having different interests regarding their offspring’s handedness phenotype, and our analysis yields predictions as to patterns of such gene effects depending on the sex of parent and offspring (see Supplementary Material §§S1.7 and S2.5). Schmitz et al. (2022)’s analyses suggested parental effects on hand preference, and stronger maternal effects than paternal effects in another multidimensional laterality trait—footedness. Parental genetic effects have been suggested to arise in neurodevelopmental disorders associated with handedness, such as maternal genetic effects in relation to loci associated with ASD—potential loci include *SHANK3* on chromosome 22 and *WBSCR17* on chromosome 7q11—but these findings are not replicated in other individual data set (Connolly et al. 2017). The predictions of our analysis therefore offer a new perspective for understanding the role of parental genetic effects in neurodevelopmental disorders.

Finally, our analysis also reveals maternal-origin versus paternal-origin genes within an individual’s own genome may come into conflict in relation to their carrier’s handedness phenotype, and how this conflict may lead to the evolution of parent-of-origin-specific gene expression. Genomic imprinting is associated with a variety of debilitating disorders, with parent-of-origin-specific clinical effects and nonstandard patterns of inheritance that are often predictable in light of the kinship theory (Wilkins & Ubeda 2011). Our results concerning patterns of imprinting allow us to make predictions as to the effects of a range of different mutational and epimutational perturbations of imprinted loci affecting handedness (Figure S3). For example, a gene deletion at an imprinted locus is expected to have no impact on the phenotype if the gene was to be silenced anyway, but it is expected to have a—potentially major—impact upon the phenotype if it was to be expressed such that no functional gene product at all will derive from the affected locus (Figure S3). Such effects might often be lethal insofar as they involve disruption to early stages of brain development when left-right asymmetry is usually established. These predictions could potentially enhance our understanding of various neurodevelopmental disorders associated with handedness. A range of neurodevelopmental conditions are associated with elevated level of left (or non-right) handedness, e.g. dyslexia or developmental language disorders (Abbondanza et al. 2023, Packheiser et al. 2023), schizophrenia (Hirnstein & Hugdahl 2014), and ASD (Markou et al. 2017). Several loci that are associated with ASD have been suggested to have a parent-of-origin effects—with maternally over-expressed components including a region between *LOC391642* and *LOC645641* on chromosome 4 and the *LRRC16A* gene on chromosome 6, and paternally over-transmitted genes including the *STPG2* gene on chromosome 4 and the *TBC1D4* gene on chromosome 13— but these findings are not replicated (Connolly et al. 2017). Considering novel parent-of-origin effects on complex traits have recently been reported with larger samples and new method such as probabilistic approach (Hofmeister et al. 2022), we suggest parent-of-origin effects might be more widespread than anticipated.

In relation to parent-of-origin effects, we have focused on the “loudest voice prevails” model of the evolution of genomic imprinting (Haig 1996), which applies here to loci whereby a greater level of gene expression either increases (“left-handedness promoter”) or decreases (“left-handedness inhibitor”) the likelihood of the individual exhibiting left-handedness. For loci at which an intermediate level of gene expression yields a right-handed phenotype and deviations in gene expression (in either direction) are liable to yield a left-handed phenotype—in line with the developmental instability hypothesis of handedness (Yeo & Gangestad 1993)—we might instead expect the gene that favours a greater incidence of left-handedness to exhibit more stochastic expression, i.e. the “chaotic voice prevails” logic of Úbeda et al. (2014). More generally, the kinship theory of genomic imprinting, as it currently stands, predicts genomic imprinting of all loci that experience parent-of-origin conflict, yet empirical studies suggest that genomic imprinting is quite rare—around 1% of genes in the human genome (Luedi et al. 2007). Clearly, there are additional requirements for a locus to evolve imprinting, and our hope is that through confronting theoretical predictions with empirical data, the theory can be further refined.

The strong population bias in favour of one sidedness type while the other remains a substantial minority appears to be an exclusively human phenomenon. However, lateralization itself has a taxonomically widespread occurrence. The historical view that lateralization is unique in humans was disputed in 1970s during a renaissance of lateralization studies (Güntürkün et al. 2020), and since then lateralization has been reported across the animal kingdom (Rogers 1980, Vallortigara & Bisazza 2002, Ocklenburg & Güntürkün 2012, Ströckens et al. 2013, Versace & Vallortigara 2015). Some species only show lateralization at individual level, such as paw preference in rodents (Manns et al. 2021), and in cats and dogs (Ocklenburg et al. 2019), turning preferences in insects (Hassall et al. 2007) and in fishes (Vallortigara & Rogers 2005), and eye preference in octopuses (Byrne et al. 2004). While lateralization at population level seems to be relatively rarer (Vallortigara & Rogers 2005, Meguerditchian et al. 2013), supporting evidence has steadily accumulated from studies of indoor/captive individuals and from the field (Forrester et al. 2013, Ströckens et al. 2013), including hand preference in nonhuman primates (Caspar et al. 2022), foot (Rogers 1980) and eye preferences (Brown & Magat 2011) in Australian parrots, left-leg preference for prey touching in spitting spiders (Ades & Ramires 2002), right-leg preference in kicking undesirable males by female mosquitoes (Benelli et al. 2015), turning bias in ants (Hunt et al. 2014) and a higher frequency of being attacked on the right in trilobites (Babcock 1993).

Ghirlanda et al. (2009) argued that population-level brain lateralization can occur in two steps: first, individuals should benefit from increased cognitive efficiency by being lateralized in either direction (Levy 1977, Güntürkün et al. 2000, Rogers et al. 2004, Vallortigara & Rogers 2005, Llaurens et al. 2009); second, a population-level bias in preference to one direction should bring additional benefits, e.g. the majority of individuals moving in the same direction creates a dilution effect which reduces the chances of being eaten by predators (Ghirlanda & Vallortigara 2004, Vallortigara & Rogers 2005), while the minority may also enjoy a surprise advantage if predators learn which direction the majority of their prey prefer (Ghirlanda & Vallortigara 2004). The surprise advantage of left-handedness has been found in elite athletes competing in sports such as fencing, boxing, baseball and table tennis (Abrams & Panaggio 2012, Loffing 2017, Papadatou-Pastou et al. 2020). Though the additional benefits were first discussed in relation to prey-predator interactions, similar benefits might also emerge from intraspecific interactions. Ghirlanda et al (2009) and Abrams & Panaggio (2012) have suggested that the population balance of right-handers versus left-handers reflects the relative prevalence of cooperative versus competitive interactions, with cooperative interactions promoting the fitness of the majority handedness type and competitive interactions promoting the fitness of the minority handedness type. All of these game theoretical models focus on social interactions, which are very likely to be mediated by genetic relatedness as shown in general cases of social evolution, yet our investigation is the first time to consider kin selection in human handedness. The predicted effect of relatedness on the evolution of handedness crucially depends on whether left-handedness is marginally altruistic or selfish. Although current data are not sufficient for answering that question, our analyses provide a framework within which future data can be motivated and conceptualised.

## Notes

### Competing Interest Statement

The authors have declared no competing interest.

