## Supplementary Material for "Kin selection as a modulator of human handedness: sex-specific, parental and parent-of-origin effects"

**Supplementary Material for Manuscript "*Kin selection as a modulator of human handedness: sex-specific, parental and parent-of-origin effects*"**

Bing Dong[1,*], Silvia Paracchini[2], Andy Gardner[1]

1. School of Biology, University of St Andrews, Dyers Brae, St Andrews KY16 9TH, UK
2. School of Medicine, University of St Andrews, North Haugh, St Andrews KY16 9TF, UK

This Supplementary Material includes:

Figure S1
Figure S2
Figure S3

**1 | Within-group Combat**
1.1 | Population model
1.2 | Fitness
1.3 | Kin selection
1.4 | Sex-biased dispersal
1.5 | Parent-of-origin effects
1.6 | Sex-specific effects
1.7 | Parental genetic effects

**2 | Between-group Combat**

2.1 | Kin selection

2.2 | Sex-biased dispersal

2.3 | Parent-of-origin effects

2.4 | Sex-specific effects

2.5 | Parental genetic effects

**References**

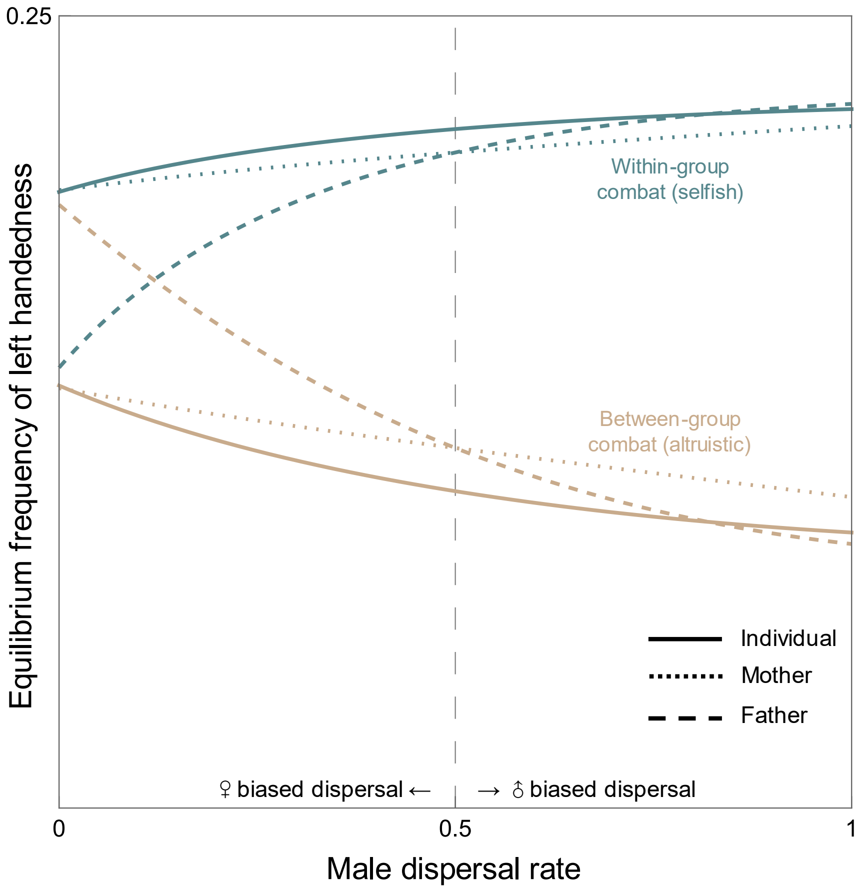


Figure S1. Maternal versus paternal genetic effects in left-handedness: level of left-handedness can be mediated by dispersal, and further result in mother-father-offspring disagreement on handedness in the context of within-group combat (left-handedness is selfish) versus between-group combat (left-handedness is altruistic). Details see §§S1.7&S2.5.

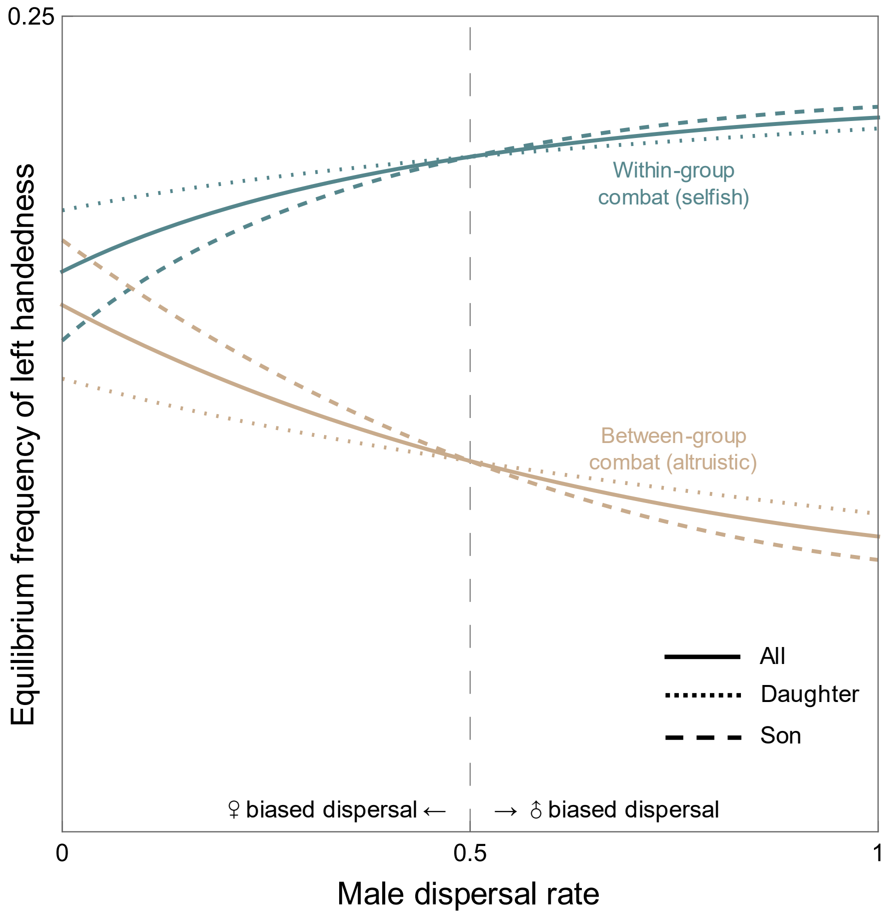


Figure S2. Parental genetic effects in left-handedness: level of left-handedness can be mediated by parental effects and dispersal pattern (female/male biased dispersal) in the context of within-group combat (left-handedness is selfish) versus between-group combat (left-handedness is altruistic). (Solid: all offspring, Dotted: daughters, Dashed: sons.)

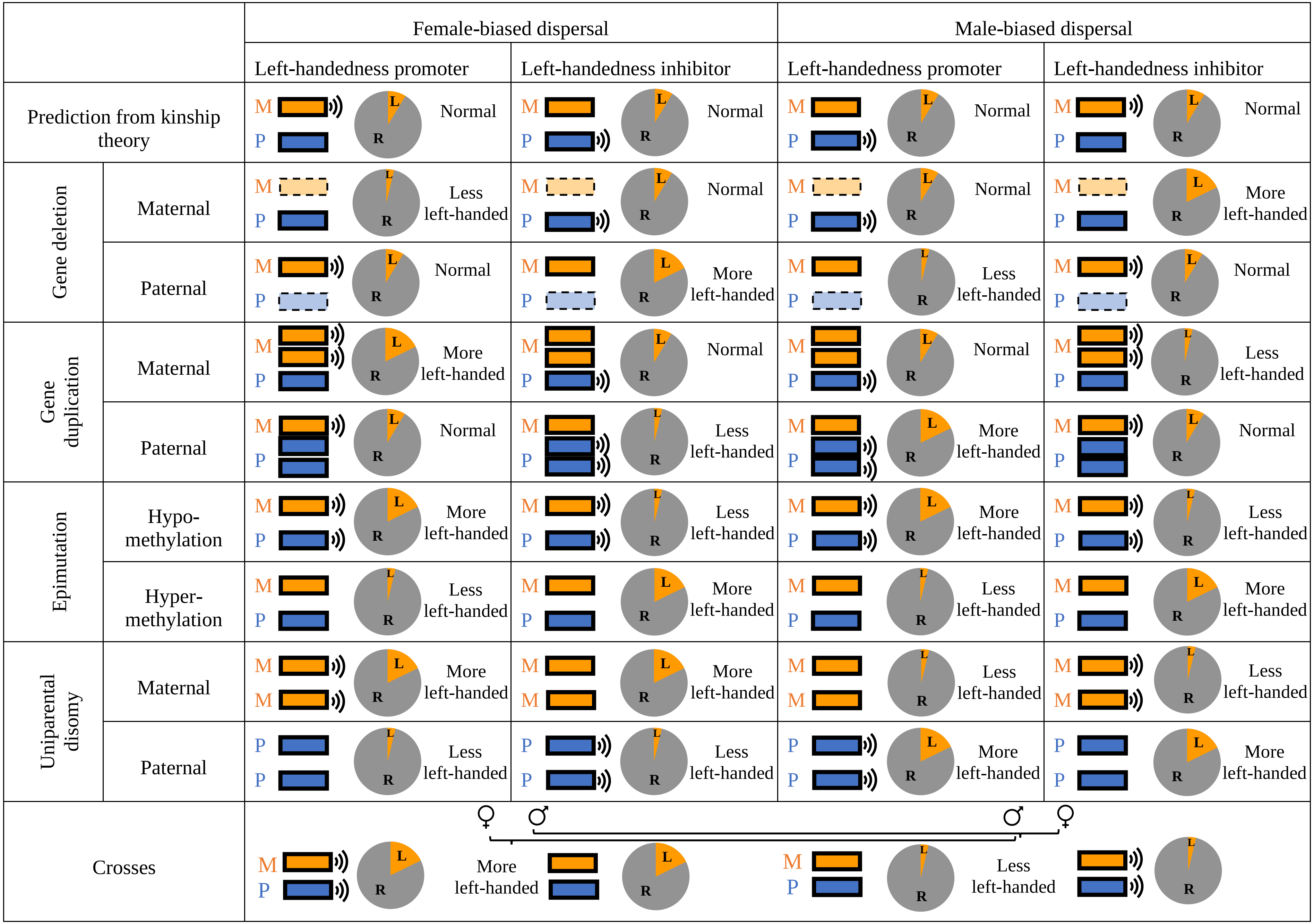


Figure S3. Phenotypic consequences on handedness of gene deletions, gene duplications, epimutations and uniparental disomies.

### 1 | Within-group combat

#### 1.1 | Population model

We assume a large population, separated into $N$ patches (where $N$ is large) each containing $n$ women and $n$ men (where $n$ may be small). Adults may engage in same-sex combat, and we model the fitness consequences of this combat by modulating the survival of their offspring to adulthood, which is mathematically equivalent to modulating the combatants' fecundity (Taylor & Frank 1996). Specifically: we assign each female a large number $K$ of offspring fathered by each male in the patch, with an even sex ratio; all parents then die; and offspring undergo random mortality, with each offspring's probability of survival depending on the handedness of their parents and of their parents' social partners, reflecting their parents' success in combat—including a surprise advantage to individuals with the rarer handedness type—and also any intrinsic disadvantage of left-handers over right-handers. Survivors then form subgroups of $n$ woman and $n$ men at random with their patch mates, and $N$ subgroups are chosen at random across the whole population with each being assigned a patch in which to live, and all other subgroups perishing—i.e. a "tribe splitting" (Haldane 1932) or "group budding" (Gardner & West 2006) model of population structure. Finally, with probability $m_f$ for women and probability $m_m$ for men, individuals may disperse away from their assigned patch to take up a random spot in another patch vacated by another same-sex disperser, such that these parameters modulate the relatedness structure of groups without affecting fitness (Gardner & West 2006).

#### 1.2 | Fitness

We assume that an individual's payoff from combat is proportional to their competitive ability relative to that of their same-sex social interactants. We assume that each individual's competitive ability is proportional to the average disposition for the opposite handedness within their social arena, such that the individual's competitive ability is greatest when their own handedness is the opposite of all of their opponents—representing the surprise advantage of the minority handedness type. For simplicity, we will often refer to handedness as if it were a binary trait, so that an individual's disposition for left-handedness is the probability that they will develop as left-handed, but more generally our analysis also applies to scenarios in which individuals exhibit quantitative degrees of left- versus right-handedness. That is: with probability $x$ the focal individual is left-handed and has competitive ability 1-$y$, where $y$ is the average disposition for left-handedness in the social arena; and with probability 1-$x$ the focal individual is right-handed and has competitive ability $y$. And the social arena is made up of a proportion $y$ of left-handed individuals with competitive ability 1-$y$ and a proportion 1-$y$ of right-handed individuals with competitive ability $y$. Accordingly, the focal individual's relative competitive ability is

$$x\frac{(1-y)}{y(1-y)+(1-y)y}+(1-x)\frac{y}{y(1-y)+(1-y)y} \tag{S1}$$

which simplifies to

$$\frac{x}{2y}+\frac{1-x}{2(1-y)} \tag{S2}$$

Hence, we may express the fitness of a focal juvenile by

$$\begin{aligned} w = &\left(1-b_{\mathrm{f}}+b_{\mathrm{f}}\left(\frac{x_{\mathrm{Mo}}}{2y_{\mathrm{Mo}}}+\frac{1-x_{\mathrm{Mo}}}{2(1-y_{\mathrm{Mo}})}\right)\right)(1-c_{\mathrm{f}}x_{\mathrm{Mo}})\left(1-b_{\mathrm{m}}\right. \\ &\left.+b_{\mathrm{m}}\left(\frac{x_{\mathrm{Fa}}}{2y_{\mathrm{Fa}}}+\frac{1-x_{\mathrm{Fa}}}{2(1-y_{\mathrm{Fa}})}\right)\right)(1-c_{\mathrm{m}}x_{\mathrm{Fa}}) \end{aligned} \tag{S3}$$

where $x_{\mathrm{Mo}}$ is the probability of the juvenile's mother developing as left-handed, $x_{\mathrm{Fa}}$ is the probability of the juvenile's father developing as left-handed, $y_{\mathrm{Mo}}$ is the probability of a random adult female from the juvenile's mother's group developing as left-handed, $y_{\mathrm{Fa}}$ is the probability of a random adult male from the focal juvenile's father's group developing as left-handed, $b_{\mathrm{f}}$ is the relative importance of combat compared with other types of competition for females, $b_m$ is the relative importance of combat for males, $c_f$ is the intrinsic cost of developing as left-handed for females and $c_m$ is the intrinsic cost of developing as left-handed for males. Average fitness $\bar{w}$ is found by substituting $x_{Mo} = y_{Mo} = z_f$, and $x_{Fa} = y_{Fa} = z_m$ in expression (S3) where $z_f$ is the population average value of left-handedness for females, and $z_m$ is the population average value of left-handedness for males. Accordingly, the relative fitness of the focal juvenile is given by $W = w/\bar{w}$ or

$$W = \left(1 - b_\mathrm{f} + b_\mathrm{f}\left(\frac{x_\mathrm{Mo}}{2y_\mathrm{Mo}} + \frac{1 - x_\mathrm{Mo}}{2(1 - y_\mathrm{Mo})}\right)\right)\left(\frac{1 - c_\mathrm{f}x_\mathrm{Mo}}{1 - c_\mathrm{f}z_\mathrm{f}}\right)\left(1 - b_\mathrm{m} + b_\mathrm{m}\left(\frac{x_\mathrm{Fa}}{2y_\mathrm{Fa}} + \frac{1 - x_\mathrm{Fa}}{2(1 - y_\mathrm{Fa})}\right)\right)\left(\frac{1 - c_\mathrm{m}x_\mathrm{Fa}}{1 - c_\mathrm{m}z_\mathrm{m}}\right) \quad \text{(S4)}$$

### 1.3 | Kin selection

#### *1.31 | Marginal fitness and evolutionary equilibrium*

We assume that genes at an autosomal locus G control their carrier's probability of developing as left-handed (see §S1.7 for the consequences of relaxing this assumption), that the two genes in this diploid locus have equal control over the individual's phenotype (see §S1.5 for the consequences of relaxing this assumption), and that genes are expressed in the same way by female and male carries (see §S1.6 for the consequences of relaxing this assumption). We denote the genic value for left-handedness of a gene drawn from locus G from a focal juvenile by $g$. We further denote the additive genetic breeding value—i.e. the average of the corresponding genic values—for left-handedness of the focal juvenile's parent by $\tilde{g}$, the average breeding value of all the adults in the focal juvenile's parents' group by $\tilde{g}'$, and the average breeding value of the population by $\bar{g}$. Employing Taylor-Frank kin-selection methodology (Taylor & Frank 1996), the condition for natural selection—the sum of direct selection and indirect (i.e. kin) selection—to favour an increase in left-handedness is given by d$W$/d$g$ > 0, where

$$\frac{\mathrm{d}W}{\mathrm{d}g} = \frac{\partial W}{\partial x_{\mathrm{Mo}}}\frac{\mathrm{d}x_{\mathrm{Mo}}}{\mathrm{d}\tilde{g}}\frac{\mathrm{d}\tilde{g}}{\mathrm{d}g} + \frac{\partial W}{\partial y_{\mathrm{Mo}}}\frac{\mathrm{d}y_{\mathrm{Mo}}}{\mathrm{d}\tilde{g}'}\frac{\mathrm{d}\tilde{g}'}{\mathrm{d}g} + \frac{\partial W}{\partial x_{\mathrm{Fa}}}\frac{\mathrm{d}x_{\mathrm{Fa}}}{\mathrm{d}\tilde{g}}\frac{\mathrm{d}\tilde{g}}{\mathrm{d}g} + \frac{\partial W}{\partial y_{\mathrm{Fa}}}\frac{\mathrm{d}y_{\mathrm{Fa}}}{\mathrm{d}\tilde{g}'}\frac{\mathrm{d}\tilde{g}'}{\mathrm{d}g}$$
$$= \left(\frac{\partial W}{\partial x_{\mathrm{Mo}}}p_{\mathrm{OM}} + \frac{\partial W}{\partial y_{\mathrm{Mo}}}p_{\mathrm{JA}} + \frac{\partial W}{\partial x_{\mathrm{Fa}}}p_{\mathrm{OF}} + \frac{\partial W}{\partial y_{\mathrm{Fa}}}p_{\mathrm{JU}}\right)\gamma \tag{S5}$$

where $p_{\mathrm{OM}}$ is the consanguinity (i.e. probability of identity by descent; Bulmer 1994) between the focal juvenile and its mother, $p_{\mathrm{JA}}$ is the consanguinity between the focal juvenile and a random adult female in its parent group, $p_{\mathrm{OF}}$ is the consanguinity between the focal juvenile and its father, $p_{\mathrm{JU}}$ is the consanguinity between the focal juvenile and a random adult male in its parent group, $\gamma = \mathrm{d}x_{\mathrm{Mo}}/\,\mathrm{d}\tilde{g} = \mathrm{d}y_{\mathrm{Mo}}/\mathrm{d}\tilde{g}' = \mathrm{d}x_{\mathrm{Fa}}/\mathrm{d}\tilde{g} = \mathrm{d}y_{\mathrm{Fa}}/\mathrm{d}\tilde{g}'$ is the mapping between genotype and phenotype, and all the derivatives are evaluated at the population average $g = \bar{g}$. Accordingly, the condition for an increase in left-handedness to be favoured is:

$$\frac{\partial W}{\partial x_{\mathrm{Mo}}}p_{\mathrm{OM}} + \frac{\partial W}{\partial y_{\mathrm{Mo}}}p_{\mathrm{JA}} + \frac{\partial W}{\partial x_{\mathrm{Fa}}}p_{\mathrm{OF}} + \frac{\partial W}{\partial y_{\mathrm{Fa}}}p_{\mathrm{JU}} > 0 \tag{S6}$$

Here for the investigation on how kin selection mediates handedness generally, we assume there is no sex-biased dispersal ($m_{\mathrm{f}} = m_{\mathrm{m}} = m$), thus $p_{\mathrm{O}} = p_{\mathrm{OM}} = p_{\mathrm{OF}}$, $p_{\mathrm{J}} = p_{\mathrm{JA}} = p_{\mathrm{JU}}$, while this assumption will be relaxed in later sections (§S1.4 Sex-biased dispersal, §S1.5 Parent-of-origin effect, §S1.6 Sex-specific effects and §S1.7 Parental genetic effects). Using expression (S4) to calculate the corresponding partial derivatives, the condition for natural selection to favour an increase in left-handedness is

$$-\frac{(b_{\mathrm{f}} + b_{\mathrm{m}})(1-2z)\left(r_{\mathrm{J}} - r_{\mathrm{O}}\right)}{2(1-z)z} - \frac{c_{\mathrm{f}}r_{\mathrm{O}}}{1-c_{\mathrm{f}}z} - \frac{c_{\mathrm{m}}r_{\mathrm{O}}}{1-c_{\mathrm{m}}z} > 0 \tag{S7}$$

where $r_{\mathrm{O}} = p_{\mathrm{O}}/p_{\mathrm{I}}$ is the relatedness between an individual and its offspring, $r_{\mathrm{J}} = p_{\mathrm{J}}/p_{\mathrm{I}}$ is the relatedness of an individual to a random adult in its parent's group, $r_{\mathrm{I}} = p_{\mathrm{I}}/p_{\mathrm{I}}$ is the relatedness of an individual to itself, and $p_{\mathrm{I}}$ is the consanguinity of a focal individual to itself. Letting $f(z)$ be the LHS of expression (S7), then at evolutionary equilibrium if there is an intermediate level of left-handedness $z^*$, this satisfies $f(z^*) = 0$. For example, setting $c_f = c_m = 1$, we have

$$z^* = \frac{1}{2}\frac{(b_f + b_m)(r_J - r_O)}{(b_f + b_m)r_J - (2 + b_f + b_m)r_O} \quad \text{(S8)}$$

*1.32 | Relatedness*

The consanguinity between a juvenile and its parent $p_O$ is given by

$$p_O = \frac{1}{2}p_I + \frac{1}{2}f \quad \text{(S9)}$$

That is: with probability 1/2 the gene picked from the juvenile comes from that parent, in which case the consanguinity is that between the parent and itself, i.e. $p_I$; and with probability 1/2 the gene comes from the other parent, in which case the consanguinity is that of mating partners, *f*. The consanguinity between the focal juvenile and a random adult in its parents' social group $p_J$ is:

$$p_J = \frac{1}{2}\left(\frac{1}{n}p_I + \frac{n-1}{n}(1-m)^2 p_x\right) + \frac{1}{2}f \quad \text{(S10)}$$

That is: with the probability 1/2 the juvenile's gene comes from the parent of the same sex as the adult, in which case with probability 1/*n* the adult is the parent and the consanguinity is $p_I$, and with probability (*n*-1)/*n* the adult is not the parent then if neither of them disperses, i.e. $(1-m)^2$, their consanguinity would be that between two random juveniles born in the same patch, $p_x$, and with probability 1/2 the juvenile's gene comes from the parent of the opposite sex, in which case the consanguinity is that of mating partners, i.e. *f*. The consanguinity between an individual and itself, $p_I$, is given by

$$p_I = \frac{1}{2} + \frac{1}{2}f \quad \text{(S11)}$$

That is: with probability 1/2 we pick the individual's same gene twice, in which case the consanguinity is $p_I$, and with probability 1/2 we pick one gene at the first time and pick the other at the second time, in which case the consanguinity is that of mating partners, i.e. $f$., and $f$ is given by

$$f = (1 - m)^2 p_{\mathrm{x}} \tag{S12}$$

That is: with probability $(1 - m)^2$ neither mating partner disperses, in which case the consanguinity is that between two random juveniles born in the same patch $p_{\mathrm{x}}$, and $p_{\mathrm{x}}$ is given by

$$p_{\mathrm{x}} = \frac{1}{4}\left(\frac{1}{n}p_{\mathrm{I}} + \frac{n-1}{n}(1-m)^2 p_{\mathrm{x}}\right) + \frac{1}{4}\left(\frac{1}{n}p_{\mathrm{I}} + \frac{n-1}{n}(1-m)^2 p_{\mathrm{x}}\right) + \frac{1}{2}f \tag{S13}$$

That is: with probability 1/4 one juvenile's gene comes from her mother and the other juvenile's gene also comes from her mother, in which case the consanguinity is that between the two mothers, which is with probability $1/n$ the two individuals share one mother, and the consanguinity is that between the mother and herself, i.e. $p_{\mathrm{I}}$, and with probability $(n-1)/n$ the two individuals do not share one mother, and if neither of the mothers disperses i.e. $(1 - m)^2$, and the consanguinity is that between two random juveniles born in the same patch, i.e. $p_{\mathrm{x}}$, and with probability 1/4 one juvenile's gene comes from her father and the other juvenile's gene also comes from her father, in which case the consanguinity is the same polynomials with the situation that the genes we pick both come from the juveniles' mothers, and with probability 1/2 one juvenile's gene comes from her mother and the other juvenile's gene comes from her father, in which case the consanguinity is that of mating partners, i.e. $f$. Solving expressions (S9)-(S13) simultaneously, we obtain

$$f = \frac{(1-m)^2}{1 + (1 - (1-m)^2)(4n-1)} \tag{S14}$$

$$p_{\mathrm{x}} = \frac{1}{1 + (1 - (1-m)^2)(4n-1)} \tag{S15}$$

$$p_{\mathrm{I}} = \frac{1 + (1 - (1-m)^2)(2n-1)}{1 + (1 - (1-m)^2)(4n-1)} \tag{S16}$$

$$p_{\mathrm{J}} = \frac{1}{1 + (1 - (1 - m)^2)(4n - 1)} \tag{S17}$$

$$p_{\mathrm{O}} = \frac{1 + (1 - (1 - m)^2)(n - 1)}{1 + (1 - (1 - m)^2)(4n - 1)} \tag{S18}$$

*1.33 | Convergence stable strategy*

As $f'(z) < 0$ is true for all the values of $z$, the equilibrium value of left-handedness is globally convergence stable (Christiansen 1991, Taylor 1996). We will use the term "optimum" or "optimal value" to be synonymous with this convergence stable strategy. Substituting all the parameters of relatedness to expression (S8), we obtain the optimum of left-handedness $z^*$:

$$z^* = \frac{1}{2}\frac{(b_{\mathrm{f}} + b_{\mathrm{m}})(1 - (1 - m)^2)(n - 1)}{(2 + b_{\mathrm{f}} + b_{\mathrm{m}})(1 - (1 - m)^2)(n - 1) + 2} \tag{S19}$$

We set the relative importance of combat relative to all types of competition for the female $b_{\mathrm{f}}$ and male $b_{\mathrm{m}}$ both to be 1, and the number of individuals each sex born in the same patch $n$ to be 5 for Figure 1*a*.

### 1.4 | Sex-biased dispersal

*1.41 | Marginal fitness and evolutionary equilibrium*

Here we relax the assumption of no sex bias in dispersal i.e. $m_{\mathrm{f}} \neq m_{\mathrm{m}}$, hence $p_{\mathrm{JA}} \neq p_{\mathrm{JU}}$. In this section, the relative fitness function is the same as expression (S4), while the consanguinity and the conditions that favour the increase of left-handedness would change. Using expression (S4) to calculate the corresponding partial derivatives, we obtain the condition for an increase in left-handedness to be favoured when we consider within-group combat

$$-\frac{(b_{\mathrm{f}}(r_{\mathrm{JA}} - r_{\mathrm{O}}) + b_{\mathrm{m}}(r_{\mathrm{JU}} - r_{\mathrm{O}}))(1 - 2z)}{2(1 - z)z} - \frac{c_{\mathrm{f}} r_{\mathrm{O}}}{1 - c_{\mathrm{f}} z} - \frac{c_{\mathrm{m}} r_{\mathrm{O}}}{1 - c_{\mathrm{m}} z} > 0 \tag{S20}$$

where $r_{JA} = p_{JA}/p_I$ is the relatedness between a juvenile and a random adult female in its mother's social group, $p_{JA}$ is the consanguinity between a juvenile and a random adult female in its mother's social group, $r_{JU} = p_{JU}/p_I$ is the relatedness between a juvenile and a random adult male in its father's social group, $p_{JU}$ is the consanguinity between a juvenile and a random adult male in its father's social group. Letting $f(z)$ be the LHS of expression (S20), (S7), then at evolutionary equilibrium if there is an intermediate level of left-handedness $z^*$, this satisfies $f(z^*) = 0$. For example, letting $c_f = c_m = 1$ i.e. no sex difference in the cost of developing as left-handed, we obtain

$$z^* = \frac{b_f r_{JA} + b_m r_{JU} - (b_f + b_m) r_O}{2(b_f r_{JA} + b_m r_{JU} - (2 + b_f + b_m) r_O)} \qquad \text{(S21)}$$

This is the overall optima of left-handedness for all the loci involved, as $f'(z) < 0$ is true for all the values of $z$.

*1.42 | Relatedness*

Substituting the dispersal rate $m$ in $p_J$ (S10) with female dispersal rate $m_f$, we obtain the consanguinity between a juvenile and a random adult female in its mother's group $p_{JA}$

$$p_{JA} = \frac{1}{2}\left(\frac{1}{n}p_I + \frac{n-1}{n}(1 - m_f)^2 p_x'\right) + \frac{1}{2}f' \qquad \text{(S22)}$$

Substituting the dispersal rate $m_f$ in $p_{JA}$ (S22) with male dispersal rate $m_m$, we obtain the consanguinity between a juvenile and a random adult male in its father's group $p_{JU}$

$$p_{JU} = \frac{1}{2}\left(\frac{1}{n}p_I + \frac{n-1}{n}(1 - m_m)^2 p_x'\right) + \frac{1}{2}f' \qquad \text{(S23)}$$

Substituting the corresponding $m$ with $m_f$ and $m_m$ in $p_x$ (S13), we obtain the consanguinity between two random juveniles born in the same patch $p_x$'

$$p_x' = \frac{1}{4}\left(\frac{1}{n}p_I + \frac{n-1}{n}(1 - m_f)^2 p_x'\right) + \frac{1}{4}\left(\frac{1}{n}p_I + \frac{n-1}{n}(1 - m_m)^2 p_x'\right) + \frac{1}{2}f' \qquad \text{(S24)}$$

Substituting the dispersal rate $m$ in expression (S12) with $m_f$ and $m_m$, we obtain the consanguinity between mating partners $f'$

$$f' = (1 - m_{\mathrm{f}})(1 - m_{\mathrm{m}})p_{\mathrm{x}}' \quad \text{(S25)}$$

*1.43 | Convergence stable strategy*

Substituting all the parameters of relatedness with expression (S22) in expression (S21), we obtain the optimal value of left-handedness $z^*$:

$$z^* = ((n-1)(\Delta b \Delta m(\bar{m}-1) + 4\bar{b}(\bar{m}-2)\bar{m}n))/(-8n + 2(n-1)(\Delta b \Delta m(\bar{m} - 1) + 4(1+\bar{b})(\bar{m}-2)\bar{m}n)) \quad \text{(S26)}$$

where $\Delta m = m_{\mathrm{f}} - m_{\mathrm{m}}$, $\bar{m} = (m_{\mathrm{f}} + m_{\mathrm{m}})/2$, $\Delta b = b_{\mathrm{f}} - b_{\mathrm{m}}$, $\bar{b} = (b_{\mathrm{f}} + b_{\mathrm{m}})/2$.

**1.5 | Parent-of-origin effects**

*1.51 | Marginal fitness and evolutionary equilibrium*

Here we consider how the origin of genes mediates the role of kin selection in the optimum of different set of genes under the circumstances of within-group combat. We now relax the assumption that the gene's influence on the phenotype is independent of its parent of origin, and we consider sex-specific dispersal as well ($m_{\mathrm{f}} \neq m_{\mathrm{m}}$). In this section, the relative fitness function is the same as expression (S4), while the conditions that favour the increase of left-handedness would change. If only the maternal-origin gene at locus G affects the individual's handedness phenotype, then:

$$\frac{\mathrm{d}W}{\mathrm{d}g} = \frac{\partial W}{\partial x_{\mathrm{Mo}}}\frac{\mathrm{d}x_{\mathrm{Mo}}}{\mathrm{d}\tilde{g}_{\mathrm{M}}}\frac{\mathrm{d}\tilde{g}_{\mathrm{M}}}{\mathrm{d}g} + \frac{\partial W}{\partial y_{\mathrm{Mo}}}\frac{\mathrm{d}y_{\mathrm{Mo}}}{\mathrm{d}\tilde{g}_{\mathrm{M}}'}\frac{\mathrm{d}\tilde{g}_{\mathrm{M}}'}{\mathrm{d}g} + \frac{\partial W}{\partial x_{\mathrm{Fa}}}\frac{\mathrm{d}x_{\mathrm{Fa}}}{\mathrm{d}\tilde{g}_{\mathrm{M}}}\frac{\mathrm{d}\tilde{g}_{\mathrm{M}}}{\mathrm{d}g} + \frac{\partial W}{\partial y_{\mathrm{Fa}}}\frac{\mathrm{d}y_{\mathrm{Fa}}}{\mathrm{d}\tilde{g}_{\mathrm{M}}'}\frac{\mathrm{d}\tilde{g}_{\mathrm{M}}'}{\mathrm{d}g} \quad \text{(S27)}$$

where $\tilde{g}_{\mathrm{M}}$ is the genic value of an individual's maternal-origin genes at locus G, $\tilde{g}_{\mathrm{M}}'$ is the average genic value of the individual's female social partners' maternal-origin genes at locus G, $\frac{\mathrm{d}x_{\mathrm{Mo}}}{\mathrm{d}\tilde{g}_{\mathrm{M}}} = \frac{\mathrm{d}y_{\mathrm{Mo}}}{\mathrm{d}\tilde{g}_{\mathrm{M}'}} = \frac{\mathrm{d}x_{\mathrm{Fa}}}{\mathrm{d}\tilde{g}_{\mathrm{M}}} = \frac{\mathrm{d}y_{\mathrm{Fa}}}{\mathrm{d}\tilde{g}_{\mathrm{M}'}} = \gamma_{\mathrm{M}}$ describes the mapping between maternal-origin gene and phenotype, $\frac{\mathrm{d}\tilde{g}_{\mathrm{M}}}{\mathrm{d}g} = p_{\mathrm{OM|\text{-}M}}$ is the consanguinity between a juvenile and its mother conditional on picking the mother's maternal-origin genes, $\frac{\mathrm{d}\tilde{g}_{\mathrm{M}'}}{\mathrm{d}g} = p_{\mathrm{JA|\text{-}M}}$ is the consanguinity between a juvenile and a random female adult in its parent group conditional on picking the adult female's maternal-origin genes, $\frac{\mathrm{d}\tilde{g}_{\mathrm{M}}}{\mathrm{d}g}$ = $p_{\mathrm{OF|\text{-}M}}$ is the consanguinity between a juvenile and its father conditional on picking the father's maternal-origin genes, $\frac{\mathrm{d}\tilde{g}_{\mathrm{M}'}}{\mathrm{d}g}$ = $p_{\mathrm{JU|\text{-}M}}$ is the consanguinity between a juvenile and a random male adult in its parent group conditional on picking the adult male's maternal-origin genes. We have $p_{\mathrm{O|\text{-}M}} = p_{\mathrm{OM|\text{-}M}} = p_{\mathrm{OF|\text{-}M}}$. Thus the condition that favours the increase of the probability of being left-handed from the perspective of maternal-origin genes is:

$$\frac{\partial W}{\partial x_{\mathrm{Mo}}} r_{\mathrm{OM|-M}} + \frac{\partial W}{\partial y_{\mathrm{Mo}}} r_{\mathrm{JA|-M}} + \frac{\partial W}{\partial x_{\mathrm{Fa}}} r_{\mathrm{OF|-M}} + \frac{\partial W}{\partial y_{\mathrm{Fa}}} r_{\mathrm{JU|-M}} > 0 \qquad (S28)$$

where $r_{\mathrm{OM|\text{-}M}} = \frac{p_{\mathrm{OM|-M}}}{p_{\mathrm{I}}}$, $r_{\mathrm{JA|\text{-}M}} = \frac{p_{\mathrm{JA|-M}}}{p_{\mathrm{I}}}$, $r_{\mathrm{OF|\text{-}M}} = \frac{p_{\mathrm{OF|-M}}}{p_{\mathrm{I}}}$, $r_{\mathrm{JU|\text{-}M}} = \frac{p_{\mathrm{JU|-M}}}{p_{\mathrm{I}}}$. Similarly, if only the paternal-origin gene at locus G affects the individual's handedness phenotype, then the condition that favours the increase of the probability of being left-handed from the perspective of paternal-origin genes is:

$$\frac{\partial W}{\partial x_{\mathrm{Mo}}} r_{\mathrm{OM|-P}} + \frac{\partial W}{\partial y_{\mathrm{Mo}}} r_{\mathrm{JA|-P}} + \frac{\partial W}{\partial x_{\mathrm{Fa}}} r_{\mathrm{OF|-P}} + \frac{\partial W}{\partial y_{\mathrm{Fa}}} r_{\mathrm{JU|-P}} > 0 \qquad (S29)$$

where $r_{\mathrm{OM|\text{-}P}} = \frac{p_{\mathrm{OM|-P}}}{p_{\mathrm{I}}}$, $r_{\mathrm{JA|\text{-}P}} = \frac{p_{\mathrm{JA|-P}}}{p_{\mathrm{I}}}$, $r_{\mathrm{OF|\text{-}P}} = \frac{p_{\mathrm{OF|-P}}}{p_{\mathrm{I}}}$, $r_{\mathrm{JU|\text{-}P}} = \frac{p_{\mathrm{JU|-P}}}{p_{\mathrm{I}}}$, and $p_{\mathrm{OM|\text{-}P}}$ is the consanguinity between a juvenile and its mother conditional on picking the mother's paternal-origin genes, $p_{\mathrm{JA|\text{-}P}}$ is the consanguinity between a juvenile and a random adult female in its parent group conditional on picking the adult female's paternal-origin genes, $p_{\mathrm{OF|\text{-}P}}$ is the consanguinity between a juvenile and its father conditional on picking the father's paternal-origin genes, $p_{\mathrm{JU|\text{-}P}}$ is the consanguinity between a juvenile and a random adult male in its parent group conditional on picking the adult male's paternal-origin genes. We have $p_{\mathrm{O|\text{-}P}} = p_{\mathrm{OM|\text{-}P}} = p_{\mathrm{OF|\text{-}P}}$. Letting the LHS of the expression (S28) be $f(z_{\mathrm{M}})$ and that of condition (S29) be $f(z_{\mathrm{P}})$, then at evolutionary equilibrium if there is an intermediate level of left-handedness $z_{\mathrm{M}}^{*}$ and $z_{\mathrm{P}}^{*}$, this satisfies $f(z_{\mathrm{M}}) = 0$ and $f(z_{\mathrm{P}}) = 0$ respectively, and we obtain

$$z_{\mathrm{M}}^{*} = \frac{1}{2}\frac{b_{\mathrm{f}}r_{\mathrm{JA|-M}} + b_{\mathrm{m}}r_{\mathrm{JU|-M}} - (b_{\mathrm{f}} + b_{\mathrm{m}})r_{\mathrm{O|-M}}}{b_{\mathrm{f}}r_{\mathrm{JA|-M}} + b_{\mathrm{m}}r_{\mathrm{JU|-M}} - (2 + b_{\mathrm{f}} + b_{\mathrm{m}})r_{\mathrm{O|-M}}} \quad \text{(S30)}$$

$$z_{\mathrm{P}}^{*} = \frac{1}{2}\frac{b_{\mathrm{f}}r_{\mathrm{JA|-P}} + b_{\mathrm{m}}r_{\mathrm{JU|-P}} - (b_{\mathrm{f}} + b_{\mathrm{m}})r_{\mathrm{O|-P}}}{b_{\mathrm{f}}r_{\mathrm{JA|-P}} + b_{\mathrm{m}}r_{\mathrm{JU|-P}} - (2 + b_{\mathrm{f}} + b_{\mathrm{m}})r_{\mathrm{O|-P}}} \quad \text{(S31)}$$

where $r_{\mathrm{O|-M}} = \frac{p_{\mathrm{O|-M}}}{p_{\mathrm{I}}}$, $r_{\mathrm{O|-P}} = \frac{p_{\mathrm{O|-P}}}{p_{\mathrm{I}}}$ and, $z_{\mathrm{M}}^{*}$ and $z_{\mathrm{P}}^{*}$ are the optima of left-handedness from the perspective of maternal- and paternal-origin genes, as $f'(z_{\mathrm{M}}) < 0$ and $f'(z_{\mathrm{P}}) < 0$ are true for all the values of $z$.

*1.52 | Relatedness*

The consanguinity between mother and offspring from the perspective of the mother's own maternal-origin genes is

$$p_{\mathrm{OM|-M}} = \frac{1}{2}\left(\frac{1}{2} + \frac{1}{2}f'\right) + \frac{1}{2}(1 - m_{\mathrm{f}})(1 - m_{\mathrm{m}})\left(\frac{1}{2}\left(\frac{1}{n}p_{\mathrm{I}} + \frac{n-1}{n}(1 - m_{\mathrm{f}})^{2}p_{\mathrm{x}}'\right) + \frac{1}{2}f'\right) \quad \text{(S32)}$$

That is: with probability 1/2 of picking the juvenile's gene that is inherited from the mother, in which case the consanguinity is, with probability 1/2 this gene is the mother's maternal-origin genes, and the consanguinity is that between the mother's maternal gene to itself which is 1, and with probability 1/2 the juvenile's gene picked is not the mother's maternal-origin genes, and the consanguinity if that between mating partners i.e. *f'*, and with probability 1/2 of picking the individual's gene that is inherited from the father, in which case the consanguinity is that between the father and the mother's maternal-origin genes, which is the probability that neither the mother nor the father disperses $(1 - m_{\mathrm{f}})(1 - m_{\mathrm{m}})$, and then with probability 1/2 of picking the father's gene that comes from his mother, and with probability $1/n$ the father and the mother share the same mother, and the consanguinity is that of the mother to herself i.e. $p_{\mathrm{I}}$, and with the probability $(n-1)/n$ the father and the mother do not share mother, with probability that neither of the two mothers disperse $(1 - m_{\mathrm{f}})^{2}$, and the consanguinity is that between two random juveniles born in the same patch i.e. $p_x'$, plus the probability 1/2 of picking the father's genes that come from his father, times the consanguinity between mating partners $f'$. The consanguinity between a juvenile and its father's maternal-origin genes $p_{\mathrm{OF|-M}}$ is

$$p_{\mathrm{OF|-M}} = \frac{1}{2}(1-m_\mathrm{f})(1-m_\mathrm{m})\left(\frac{1}{2}\left(\frac{1}{n}p_\mathrm{I} + \frac{n-1}{n}(1-m_\mathrm{f})^2 p_\mathrm{x}'\right) + \frac{1}{2}f'\right) + \frac{1}{2}\left(\frac{1}{2} + \frac{1}{2}f'\right) \quad \text{(S33)}$$

That is: with probability 1/2 of picking the juvenile's gene that comes from its mother, in which case the consanguinity is that between the mother and the father's maternal-origin genes, which is with probability $(1-m_\mathrm{f})(1-m_\mathrm{m})$ that neither the mother nor the father disperses, and with probability 1/2 of picking the mother's maternal-origin genes, with probability $1/n$ that the mother and father share the same mother, and the consanguinity is that of the mother to herself i.e. $p_\mathrm{I}$, and with probability $(n-1)/n$ the mother and father do not share mother, with probability $(1-m_\mathrm{f})^2$ neither of the two mothers disperses, and the consanguinity is that between two random juveniles born in the same patch i.e. $p_x'$, with probability 1/2 of picking the mother's paternal-origin genes, and the consanguinity is that between mating partners i.e. $f'$, and with probability 1/2 of picking the juvenile's gene that comes from the father, in which case the consanguinity is, with probability 1/2 this gene is the father's maternal-origin genes, then and the consanguinity is that of the father's maternal-origin gene to itself which is 1, and with probability 1/2 the juvenile's gene is not the father's maternal-origin gene, then the consanguinity is that between mating partners $f'$. Hence we have $p_{\mathrm{OP|-M}} = p_{\mathrm{OM|-M}} = p_{\mathrm{OF|-M}}$. The consanguinity between a juvenile and the maternal-origin genes of a random female in its mother's social group $p_{\mathrm{JA|-M}}$ is

$$p_{\mathrm{JA|-M}} = \frac{1}{2}\left(\frac{1}{n}p_{\mathrm{I}} + \frac{n-1}{n}(1-m_{\mathrm{f}})^2\left(\frac{1}{2}\left(\frac{1}{n}p_{\mathrm{I}} + \frac{n-1}{n}(1-m_{\mathrm{f}})^2 p_{\mathrm{x}}'\right) + \frac{1}{2}f'\right)\right) + \frac{1}{2}(1-m_{\mathrm{f}})(1-m_{\mathrm{m}})\left(\frac{1}{2}\left(\frac{1}{n}p_{\mathrm{I}} + \frac{n-1}{n}(1-m_{\mathrm{f}})^2 p_{\mathrm{x}}'\right) + \frac{1}{2}f'\right) \quad (S34)$$

That is: with probability 1/2 of picking the juvenile's maternal-origin gene, in which case the consanguinity is that between the juvenile's mother and the maternal-origin genes of a random adult female in the mother's social group (including the mother), which is with probability $1/n$ that the adult female is the juvenile's mother, then the consanguinity is that of an individual to itself i.e. $p_{\mathrm{I}}$, plus the probability $(n$-$1)/n$ that the adult female is not the juvenile's mother, then the consanguinity is with probability $(1-m_{\mathrm{f}})^2$ that neither of these two females disperses, and with probability 1/2 of picking the maternal-origin gene of the juvenile's mother, then with probability $1/n$ that the two females share one mother, and the consanguinity is that of the mother to herself i.e. $p_{\mathrm{I}}$, and with probability $(n$-$1)/n$ that the two females do not share one mother, with probability $(1-m_{\mathrm{f}})^2$ that neither of the mothers of these two females disperses, and the consanguinity is that between two random juveniles born in the same patch i.e. $p_{\mathrm{x}}'$, and with probability 1/2 of picking the gene of the paternal-origin genes of the juvenile's mother, times the consanguinity of mating partners i.e. $f'$, and with probability 1/2 of picking the juvenile's paternal-origin gene, in which case the consanguinity is that between the juvenile's father and the maternal-origin gene of a random adult female in the mother's social group, which is the probability $(1-m_{\mathrm{f}})(1-m_{\mathrm{m}})$ that neither of the adult female nor the juvenile's father disperses, and with probability 1/2 of picking the maternal-origin gene of the father, with probability $1/n$ that the juvenile's father and the adult female share one mother, and the consanguinity is that of the mother to herself i.e. $p_{\mathrm{I}}$, and with probability $(n$-$1)/n$ that the juvenile's father and the female do not share one mother, with probability $(1-m_{\mathrm{f}})^2$ that neither of the mothers of these two individuals disperses, and the consanguinity is that between two random juveniles born in the same patch i.e. $p_{\mathrm{x}}'$, with probability 1/2 of picking the paternal-origin gene of the father, then the consanguinity is that between mating partners i.e. $f'$. The consanguinity between the focal juvenile and the maternal-origin gene of a random male in its father's social group $p_{\mathrm{JU|-M}}$ is

$$p_{\mathrm{JU|-M}} = \frac{1}{2}(1-m_{\mathrm{f}})(1-m_{\mathrm{m}})\left(\frac{1}{2}\left(\frac{1}{n}p_{\mathrm{I}} + \frac{n-1}{n}(1-m_{\mathrm{f}})^2 p_{\mathrm{x}}'\right) + \frac{1}{2}f'\right) + \frac{1}{2}\left(\frac{1}{n}p_{\mathrm{I}} + \frac{n-1}{n}(1-m_{\mathrm{m}})^2\left(\frac{1}{2}\left(\frac{1}{n}p_{\mathrm{I}} + \frac{n-1}{n}(1-m_{\mathrm{f}})^2 p_{\mathrm{x}}'\right) + \frac{1}{2}f'\right)\right) \quad (S35)$$

That is: with probability 1/2 of picking the juvenile's gene that comes from the mother, in which case the consanguinity is that between the juvenile's mother and the maternal-origin genes of a random adult male in the father's social group, which is with probability $(1-m_{\mathrm{f}})(1-m_{\mathrm{m}})$ that neither the mother nor the adult male disperses, with probability 1/2 of picking the mother's maternal-origin genes, with probability $1/n$ these two genes come from the same mother and the consanguinity is that of the mother to herself i.e. $p_{\mathrm{I}}$, and with probability $(n-1)/n$ these two genes come from different mothers, with probability $(1-m_{\mathrm{f}})^2$ that neither of the two mothers disperses, and the consanguinity is that between two random juveniles born in the same patch i.e. $p_{\mathrm{x}}'$, and with probability 1/2 of picking the mother's paternal-origin gene, and the consanguinity is that of mating partners i.e. $f'$, and with probability 1/2 of picking the juvenile's gene that comes from the father, in which case the consanguinity is that between the juvenile's father and the maternal-origin genes of a random adult male in the father's social group (including this father), which is with probability $1/n$ these two genes come from the same mother, and the consanguinity is that of the mother to herself i.e. $p_{\mathrm{I}}$, with probability $(n-1)/n$ these two genes comes from different mothers, with probability $(1-m_{\mathrm{m}})^2$ neither of the two males disperses, and with probability 1/2 of picking the father's maternal-origin gene, with probability $1/n$ the juvenile's father and the random male in the father's group share one mother, and the consanguinity is that between the mother and herself i.e. $p_I$, with probability ($n$-1)/$n$ the two males do not share one mother, with probability $(1-m_f)^2$ that neither of the two mothers of the two males disperses, and the consanguinity is that between two random juveniles born in the same patch $p_x'$, with probability 1/2 of picking the juvenile's father's paternal-origin gene, and the consanguinity is that between mating partners i.e. $f'$. The consanguinity between a juvenile and its mother from the perspective of the mother's paternal-origin gene $p_{\mathrm{OM|-P}}$ is

$$p_{\mathrm{OM|-P}} = \frac{1}{2}\left(\frac{1}{2}f' + \frac{1}{2}\right) + \frac{1}{2}(1-m_f)(1-m_m)\left(\frac{1}{2}f' + \frac{1}{2}\left(\frac{1}{n}p_I + \frac{n-1}{n}(1-m_m)^2 p_x'\right)\right) \quad \text{(S36)}$$

That is: with probability 1/2 of picking the juvenile's gene that comes from the mother, in which case the consanguinity is that between the mother and the mother's paternal-origin gene, which is with probability 1/2 the gene is the mother's maternal-origin genes, and the consanguinity is that between the mother's maternal-origin genes and its paternal-origin genes i.e. $f'$, and with probability 1/2 the juvenile's gene picked is the mother's paternal-origin genes, then the consanguinity is 1, and with probability 1/2 of picking the juvenile's gene that comes from its father, in which case the consanguinity is that between the mother's maternal-origin genes and the father, which is with probability $(1-m_f)(1-m_m)$ neither of the mother and father disperses, and with probability 1/2 of picking the father's maternal-origin gene, and the consanguinity is that between mating partners i.e. $f'$, and with probability 1/2 of picking the father's paternal-origin gene, and with probability 1/$n$ the mother and father share the same father, and the consanguinity is that of the mother to herself i.e. $p_I$, and with probability ($n$-1)/$n$ the mother and father do not share father, with probability $(1-m_m)^2$ neither of the two fathers disperses, and the consanguinity is that between two random juveniles born in the same patch i.e. $p_x'$. From expression (S32) and (S33), according to the same rule we can get $p_{\mathrm{O|-P}} = p_{\mathrm{OM|-P}} = p_{\mathrm{OF|-P}}$. The consanguinity between a juvenile and a random adult female in its mother's social group (including the mother) from the perspective of the adult female's paternal-origin genes $p_{\mathrm{JA|-P}}$ is

$$p_{\mathrm{JA|-P}} = \frac{1}{2}\left(\frac{1}{n}p_{\mathrm{I}} + \frac{n-1}{n}(1-m_{\mathrm{f}})^2\left(\frac{1}{2}f' + \frac{1}{2}\left(\frac{1}{n}p_{\mathrm{I}} + \frac{n-1}{n}(1-m_{\mathrm{m}})^2 p_{\mathrm{x}}'\right)\right)\right) + \frac{1}{2}(1-m_{\mathrm{f}})(1-m_{\mathrm{m}})\left(\frac{1}{2}f' + \frac{1}{2}\left(\frac{1}{n}p_{\mathrm{I}} + \frac{n-1}{n}(1-m_{\mathrm{m}})^2 p_{\mathrm{x}}'\right)\right) \quad \text{(S37)}$$

That is: with probability 1/2 of picking the juvenile's gene that come from the mother, in which case the consanguinity is that between the juvenile's mother and the paternal-origin genes of a random adult female in the mother's social group, which is with probability $1/n$ the adult female is the juvenile's mother, times the consanguinity of the mother to herself $p_{\mathrm{I}}$, and with probability $(n\text{-}1)/n$ that the adult female is not the juvenile's mother, and with probability $(1-m_{\mathrm{f}})^2$ that neither of the two females disperses, with probability 1/2 of picking the juvenile's mother's maternal-origin gene, and the consanguinity is that between the mother's maternal-origin genes and paternal-origin genes i.e. $f'$, and with probability 1/2 of picking the mother's paternal-origin genes, with probability $1/n$ the juvenile's mother and the random female in the mother's group share one father, and the consanguinity is that between the father and himself i.e. $p_{\mathrm{I}}$, and with probability $(n\text{-}1)/n$ the two females do not share one father, with probability $(1-m_{\mathrm{m}})^2$ neither of the two fathers of the two females disperses, and the consanguinity is that between two random juveniles born in the same patch i.e. $p_{\mathrm{x}}'$, and with probability 1/2 of picking the juvenile's gene that comes from the father, in which case the consanguinity is that between the juvenile's father and the paternal-origin genes of a random adult female in the mother's group, which is with probability $(1-m_{\mathrm{f}})(1-m_{\mathrm{m}})$ that neither the adult female nor the father disperses, and with probability 1/2 of picking the father's maternal-origin gene, and the consanguinity is that between mating partners i.e. $f'$, with probability 1/2 of picking the father's paternal-origin gene, and with probability $1/n$ that the adult female and the father share one father, and the consanguinity is that of the father to himself i.e. $p_\mathrm{I}$, and with probability ($n$-1)/$n$ the adult female and the father do not share one father, and with probability $(1-m_\mathrm{m})^2$ neither of the two fathers disperses, and the consanguinity is that between two random juveniles born in the same patch i.e. $p_\mathrm{x}'$. The consanguinity between a juvenile and the paternal-origin gene of a random adult male in its father's social group (including the father) $p_{\mathrm{JU|-P}}$ is:

$$p_{\mathrm{JU|-P}} = \frac{1}{2}(1-m_\mathrm{f})(1-m_\mathrm{m})\left(\frac{1}{2}f' + \frac{1}{2}\left(\frac{1}{n}p_\mathrm{I} + \frac{n-1}{n}(1-m_\mathrm{m})^2 p_\mathrm{x}'\right)\right) + \frac{1}{2}\left(\frac{1}{n}p_\mathrm{I} + \frac{n-1}{n}(1-m_\mathrm{m})^2\left(\frac{1}{2}f' + \frac{1}{2}\left(\frac{1}{n}p_\mathrm{I} + \frac{n-1}{n}(1-m_\mathrm{m})^2 p_\mathrm{x}'\right)\right)\right) \tag{S38}$$

That is: with probability 1/2 of picking the juvenile's maternal-origin gene, in which case the consanguinity is that between the juvenile's mother and the paternal-origin genes of a random adult male in the father's social group, which is the probability $(1-m_\mathrm{f})(1-m_\mathrm{m})$ that neither of the juvenile's mother nor the adult male disperses, and with probability 1/2 of picking the maternal-origin gene of the mother, and the consanguinity is that between mating partners i.e. $f'$, and with probability 1/2 of picking the paternal-origin gene of the mother, with probability 1/$n$ the juvenile's mother and the adult male share one father, and the consanguinity is that of the father to himself i.e. $p_\mathrm{I}$, and with probability ($n$-1)/$n$ the juvenile's mother and the adult male do not share one father, with probability $(1-m_\mathrm{m})^2$ neither of the fathers disperses, and the consanguinity is that between two random juveniles born in the same patch i.e. $p_\mathrm{x}'$, and with probability 1/2 of picking the juvenile's paternal-origin gene, in which case the consanguinity is that between the juvenile's father and the paternal-origin gene of a random adult male in the father's social group, which is with probability 1/$n$ the adult male is the juvenile's father, and the consanguinity is that of the father to himself i.e. $p_\mathrm{I}$, and with probability ($n$-1)/$n$ the adult male is not the juvenile's father, with probability $(1-m_\mathrm{m})^2$

that neither of the fathers disperses, and with probability 1/2 that picking the maternal-origin gene of the juvenile's father, and the consanguinity is that between mating partners i.e. *f'*, and with probability 1/2 of picking the paternal-origin gene of the juvenile's father, with probability 1/*n* the two males share one father, and the consanguinity of the father to himself i.e. $p_{\mathrm{I}}$, and with probability (*n*-1)/*n* the two males do not share one father, with probability $(1-m_{\mathrm{m}})^2$ that neither of the fathers disperses, and the consanguinity is that between two random juveniles born in the same patch i.e. $p_{\mathrm{x}}'$. Solving expressions (S32)-(S38) with the solutions of $p_{\mathrm{I}}$, $p_{\mathrm{x}}'$ and *f'* from previous section simultaneously, we obtain

$$p_{\mathrm{O|-M}} = ((-2\Delta m(M-2\bar{m}+1)(1-\bar{m}) + 2(1-\bar{m})(M\Delta m - 2\Delta m\bar{m} + 2m_{\mathrm{f}} + 2\bar{m} - 4)n - 8(2-\bar{m})\bar{m}n^2)) / ((8n(2\bar{m} - 1 - 4\bar{m}^2 + 3M - 4(2-\bar{m})\bar{m}n))) \quad \text{(S39)}$$

$$p_{\mathrm{JA|-M}} = -((-2\Delta m(1-m_{\mathrm{f}})^2(1-\bar{m}) + 2\Delta m(1-\bar{m})(5 - m_{\mathrm{m}} + m_{\mathrm{f}}(2m_{\mathrm{f}} - 5 + m_{\mathrm{m}}))n + (8 + m_{\mathrm{f}}{}^4 - m_{\mathrm{f}}{}^3(5 - \mathrm{m_m}) - (4 - m_{\mathrm{m}})H_{\mathrm{m}} - m_{\mathrm{f}}(8 + (4 - m_{\mathrm{m}})(1 - m_{\mathrm{m}})m_{\mathrm{m}}) - m_{\mathrm{f}}{}^2\,(m_{\mathrm{m}} - 10 + m_{\mathrm{m}}{}^2))n^2)) / ((8n^2\,(2\bar{m} - 1 - 4\bar{m}^2 + 3M - 4(2-\bar{m})\bar{m}n)))) \quad \text{(S40)}$$

$$p_{\mathrm{JU|-M}} = (2\Delta m(1-m_{\mathrm{m}})^2(1-\bar{m}) - 2\Delta m(1-\bar{m})(1 + M - 2\bar{m} + 2H_{\mathrm{m}})n + (2\Delta m(1-\bar{m})(M - 2\bar{m} + H_{\mathrm{m}}) - 8)n^2)) / ((8n^2\,(2\bar{m} - 1 - 4\bar{m}^2 + 3M - 4(2-\bar{m})\bar{m}n))) \quad \text{(S41)}$$

$$p_{\mathrm{O|-P}} = (((M - 2\bar{m} + 1) + 2\Delta m(1-\bar{m}) + 2(1-\bar{m})(2\Delta m\bar{m} - M\Delta m + 2m_{\mathrm{m}} + 2\bar{m} - 4)n - 8(2 - \bar{m})\bar{m}n^2)) / ((8n(2\bar{m} - 1 - 4\bar{m}^2 + 3M - 4(2-\bar{m})\bar{m}n))) \quad \text{(S42)}$$

$$p_{\mathrm{JA|-P}} = (-2\Delta m(1-m_{\mathrm{f}})^2(1-\bar{m}) + 2\Delta m(1-\bar{m})(1 - 2\bar{m} + M + 2H_{\mathrm{f}})n + \big(-8 - 4\Delta m(1-\bar{m})(M - \bar{m} + H_{\mathrm{f}} - m_{\mathrm{f}})\big)n^2)) / ((8n^2\,(2\bar{m} - 1 - 4\bar{m}^2 + 3M - 4(2-\bar{m})\bar{m}n))) \quad \text{(S43)}$$

$$p_{\mathrm{JU|-P}} = \big(-2\Delta m(1-m_{\mathrm{m}})^2(1-\bar{m}) + 2\Delta m(1-\bar{m})(5+M-2\bar{m}+2H_{\mathrm{m}})n + \big(-8 + m_{\mathrm{f}}{}^2(H_{\mathrm{m}} - 3m_{\mathrm{m}} + 6) - m_{\mathrm{f}}{}^3(1-m_{\mathrm{m}}) - H_{\mathrm{m}}(4 + H_{\mathrm{m}} - m_{\mathrm{m}}) + m_{\mathrm{f}}(H_{\mathrm{m}} - 8 + 6m_{\mathrm{m}} - m_{\mathrm{m}}{}^3)\big)n^2\big) \,/\, (8n^2(2\bar{m} - 1 - 4\bar{m}^2 + 3M - 4(2-\bar{m})\bar{m}n)) \tag{S44}$$

where $\Delta m = m_{\mathrm{f}} - m_{\mathrm{m}}$, $\bar{m} = (m_{\mathrm{f}} + m_{\mathrm{m}})/2$, $M = m_{\mathrm{f}}m_{\mathrm{m}}$, $\Delta b = b_{\mathrm{f}} - b_{\mathrm{m}}$, $\bar{b} = (b_{\mathrm{f}} + b_{\mathrm{m}})/2$, $H_{\mathrm{f}} = (m_{\mathrm{f}} - 2)m_{\mathrm{f}}$, $H_{\mathrm{m}} = (m_{\mathrm{m}} - 2)m_{\mathrm{m}}$.

*1.53 | Convergence stable strategy*

By solving the expression $dW/dg$ = 0, we could get the optimal value of left-handedness from the perspective of maternal-origin genes $z_{\mathrm{M}}{}^*$:

$$z_{\mathrm{M}}{}^* = ((2\bar{b}(n-1)(-H_{\mathrm{f}}(2+H_{\mathrm{f}}) + H_{\mathrm{m}}(2+H_{\mathrm{m}}) - 2\Delta m(1-\bar{m})(2+H_{\mathrm{f}}+H_{\mathrm{m}})n - 16(2-\bar{m})\bar{m}n^2))) \,/\, ((-8\bar{b}\Delta m(1-\bar{m})(2+H_{\mathrm{f}}+H_{\mathrm{m}}) + 16\Delta m(1-\bar{m})(\bar{b}(2+H_{\mathrm{f}}+H_{\mathrm{m}}) - 1 + 2\bar{m} - M)n + 2(2\bar{b}m_{\mathrm{f}}{}^4 - 32 - 4m_{\mathrm{f}}{}^3\,(2\bar{b} - 1 + m_{\mathrm{m}}) + 4m_{\mathrm{f}}{}^2\,(\bar{b} - 5 + 3m_{\mathrm{m}}) + 4m_{\mathrm{f}}(10 + 6\bar{b} - 4(\bar{b}+1)m_{\mathrm{m}} - 3m_{\mathrm{m}}{}^2 + m_{\mathrm{m}}{}^3) + 2m_{\mathrm{m}}(10b_{\mathrm{f}} - 10\bar{b}m_{\mathrm{m}} + 2(2\bar{b} - 1)m_{\mathrm{m}}{}^2 - \bar{b}m_{\mathrm{m}}{}^3 + 2(6 + 5b_{\mathrm{m}} + m_{\mathrm{m}})))n^2 - 64(\bar{b}+1)(2 - \bar{m})\bar{m}n^3)) \tag{S45}$$

where $\Delta m = m_{\mathrm{f}} - m_{\mathrm{m}}$, $\bar{m} = (m_{\mathrm{f}} + m_{\mathrm{m}})/2$, $M = m_{\mathrm{f}}m_{\mathrm{m}}$, $\Delta b = b_{\mathrm{f}} - b_{\mathrm{m}}$, $\bar{b} = (b_{\mathrm{f}} + b_{\mathrm{m}})/2$, $H_{\mathrm{f}} = (m_{\mathrm{f}} - 2)m_{\mathrm{f}}$, $H_{\mathrm{m}} = (m_{\mathrm{m}} - 2)m_{\mathrm{m}}$. Solving the expression $dW/dg$ = 0, we obtain the optimal value of left-handedness from the perspective of paternal-origin genes $z_{\mathrm{P}}{}^*$:

$$
\begin{aligned}
z_{\mathrm{P}}{}^{*} = {} & ((2\bar{b}(n-1)(-\big(H_{\mathrm{f}}(2+H_{\mathrm{f}})\big) + H_{\mathrm{m}}(2+H_{\mathrm{m}}) - 2\Delta m(1-\bar{m})(2+H_{\mathrm{f}}+H_{\mathrm{m}})n \\
& + 16(2-\bar{m})\bar{m}n^{2}))) \,/\, ((-8\Delta m\bar{b}(1-\bar{m})(2+H_{\mathrm{f}}+H_{\mathrm{m}}) \\
& + 8\Delta m(1-\bar{m})\big(b_{\mathrm{m}}H_{\mathrm{f}} - 2(b_{\mathrm{m}}+m_{\mathrm{f}})m_{\mathrm{m}} + b_{\mathrm{m}}m_{\mathrm{m}}{}^{2} \\
& + 2(b_{\mathrm{m}} - 1 + 2\bar{m}) + b_{\mathrm{f}}(2+H_{\mathrm{f}}+H_{\mathrm{m}})\big)n \\
& + 4\Big(16 + \bar{b}m_{\mathrm{f}}{}^{4} - 4(5+3\bar{b})m_{\mathrm{m}} - 2(\bar{b}-5)m_{\mathrm{m}}{}^{2} + 2(2\bar{b}-1)m_{\mathrm{m}}{}^{3} \\
& - \bar{b}m_{\mathrm{m}}{}^{4} - 2m_{\mathrm{f}}{}^{3}(2\bar{b} - 1 + m_{\mathrm{m}}) + 2m_{\mathrm{f}}{}^{2}(5\bar{b} - 1 + 3m_{\mathrm{m}}) \\
& + 2m_{\mathrm{f}}\big(4(\bar{b}+1)m_{\mathrm{m}} - 6 - 10\bar{b} - 3m_{\mathrm{m}}{}^{2} + m_{\mathrm{m}}{}^{3}\big)\Big)n^{2} + 64(\bar{b} \\
& + 1)(2-\bar{m})\bar{m}n^{3}))
\end{aligned} \qquad \text{(S46)}
$$

The optimal value of left-handedness for the perspective of the whole genes of the individual $z^*$ is:

$$
z^{*} = \frac{(n-1)(\Delta b\Delta m(1-\bar{m}) + 4\bar{b}(2-\bar{m})\bar{m})}{2(n-1)(\Delta b\Delta m(1-\bar{m}) + 8n + 4(\bar{b}+1)(2-\bar{m})\bar{m})} \qquad \text{(S47)}
$$

We set the female dispersal rate $m_{\mathrm{f}}$ to be 0.5, the relative importance of combat relative to all types of competition for the female $b_{\mathrm{f}}$ and male $b_{\mathrm{m}}$ both to be 1, and the number of individuals each sex born in the same patch $n$ to be 5 for Figure 2. For the two zoomed-in parts, the range of male dispersal rate $m_{\mathrm{m}}$ is from 0.499 to 0.501, the range for the equilibrium frequency of left-handedness is from 0.21426 to 0.21431.

### 1.6 | Sex-specific effects

#### *1.61 | Marginal fitness and evolutionary equilibrium*

Here we consider how sex effects add to the mediation of kin selection on handedness. In this section, the fitness functions of the focal juvenile are the same as previous sections. We use $g_1$ to denote the genic value for the locus G1, which affects handedness only when it is carried by a female. We use $g_2$ and to denote the genic value for the locus G2 which affects handedness only when it is carried by a male. The relative fitness functions are the same as expression (S4). Then we explore the optimal value of the level of left-handedness for locus $G_1$ which only controls the handedness trait of females. For juveniles, the relationship between the phenotype and genotype is:

$$\begin{aligned}\frac{dW}{dg_1} &= \frac{\partial W}{\partial x_{\mathrm{Mo}}}\frac{dx_{\mathrm{Mo}}}{d\tilde{g}_{1\mathrm{f}}}\frac{d\tilde{g}_{1\mathrm{f}}}{dg_1}+\frac{\partial W}{\partial y_{\mathrm{Mo}}}\frac{dy_{\mathrm{Mo}}}{d\tilde{g}_{1\mathrm{f}}'}\frac{dG_{1\mathrm{f}}'}{dg_1}+\frac{\partial W}{\partial x_{\mathrm{Fa}}}\frac{dx_{\mathrm{Fa}}}{d\tilde{g}_{1\mathrm{m}}}\frac{d\tilde{g}_{1\mathrm{m}}}{dg_1}+\frac{\partial W}{\partial y_{\mathrm{Fa}}}\frac{dy_{\mathrm{Fa}}}{d\tilde{g}_{1\mathrm{m}}'}\frac{d\tilde{g}_{1\mathrm{m}}'}{dg_1} \\ &= \left(\frac{\partial W}{\partial x_{\mathrm{Mo}}}p_{\mathrm{OM}}+\frac{\partial W}{\partial y_{\mathrm{Mo}}}p_{\mathrm{JA}}\right)\gamma_{1\mathrm{f}}+\left(\frac{\partial W}{\partial x_{\mathrm{Fa}}}p_{\mathrm{OF}}+\frac{\partial W}{\partial y_{\mathrm{Fa}}}p_{\mathrm{JU}}\right)\gamma_{1\mathrm{m}}\end{aligned} \quad \text{(S48)}$$

where $\tilde{g}_{1\mathrm{f}}$ is the additive breeding value of a juvenile for its mother's genes in locus $G_1$, $\tilde{g}_{1\mathrm{f}}'$ is the breeding value of the juvenile for a random adult female's genes in locus $G_1$, $\tilde{g}_{1\mathrm{m}}$ is the breeding value of the juvenile for its father's genes in locus $G_1$, $\tilde{g}_{1\mathrm{m}}'$ is the breeding value of the juvenile for a random adult male's genes in locus $G_1$, and $\gamma_{1\mathrm{f}}$ and $\gamma_{1\mathrm{m}}$ is the mapping between genotype and phenotype for the focal females and males respectively. According to our assumption that locus $G_1$ would only take an effect if its carrier is a female, we have $\gamma_{1\mathrm{f}} = 1$, $\gamma_{1\mathrm{m}} = 0$. Then expression (S48) can be simplified to

$$\frac{dW}{dg_1} = \frac{\partial W}{\partial x_{\mathrm{Mo}}}p_{\mathrm{OM}}+\frac{\partial W}{\partial y_{\mathrm{Mo}}}p_{\mathrm{JA}} \quad \text{(S49)}$$

Then the condition that favours the increase of left-handedness is

$$\frac{\partial W}{\partial x_{\mathrm{Mo}}}r_{\mathrm{OM}}+\frac{\partial W}{\partial y_{\mathrm{Mo}}}r_{\mathrm{JA}} > 0 \quad \text{(S50)}$$

Letting the LHS of expression (S50) be $f(z)$, as $f'(z) < 0$ is true for all the values of $z$, hence at evolutionary equilibrium if there is an intermediate level of left-handedness $z_{\mathrm{f}}^*$, this satisfies $f(z^*) = 0$, we obtain the optimum of left-handedness for all the loci that only control the handedness when they are carried by females

$$z_{\mathrm{f}}^* = \frac{1}{2}\frac{b_{\mathrm{f}}(r_{\mathrm{OM}}-r_{\mathrm{JA}})}{(1+b_{\mathrm{f}})r_{\mathrm{OM}}-b_{\mathrm{f}}r_{\mathrm{JA}}} \quad \text{(S51)}$$

Now we explore the optimum value of the probability of developing as left-handedness for locus $G_2$ which only controls the handedness trait of males. For a juvenile, the relationship between the phenotype and genotype is

$$\frac{dW}{dg_2} = \frac{\partial W}{\partial x_{\mathrm{Mo}}}\frac{dx_{\mathrm{Mo}}}{d\tilde{g}_{2\mathrm{f}}}\frac{d\tilde{g}_{2\mathrm{f}}}{dg_2} + \frac{\partial W}{\partial y_{\mathrm{Mo}}}\frac{dy_{\mathrm{Mo}}}{d\tilde{g}_{2\mathrm{f}}'}\frac{d\tilde{g}_{2\mathrm{f}}'}{dg_2} + \frac{\partial W}{\partial x_{\mathrm{Fa}}}\frac{dx_{\mathrm{Fa}}}{d\tilde{g}_{2\mathrm{m}}}\frac{d\tilde{g}_{2\mathrm{m}}}{dg_2}$$

$$+ \frac{\partial W}{\partial y_{\mathrm{Fa}}}\frac{dy_{\mathrm{Fa}}}{d\tilde{g}_{2\mathrm{m}}'}\frac{d\tilde{g}_{2\mathrm{m}}'}{dg_2} \quad \text{(S52)}$$

$$= \left(\frac{\partial W}{\partial x_{\mathrm{Mo}}}p_{\mathrm{OM}} + \frac{\partial W}{\partial y_{\mathrm{Mo}}}p_{\mathrm{JA}}\right)\gamma_{2\mathrm{f}} + \left(\frac{\partial W}{\partial x_{\mathrm{Fa}}}p_{\mathrm{OF}} + \frac{\partial W}{\partial y_{\mathrm{Fa}}}p_{\mathrm{JU}}\right)\gamma_{2\mathrm{m}}$$

where $\tilde{g}_{2\mathrm{f}}$ is the additive breeding value of a juvenile for its mother's genes in locus $G_2$, $\tilde{g}_{2\mathrm{f}}'$ is the breeding value of the juvenile for a random adult female's genes in locus $G_2$, $\tilde{g}_{2\mathrm{m}}$ is the breeding value of the juvenile for its father's genes in locus $G_2$, $\tilde{g}_{2\mathrm{m}}'$ is the breeding value of the juvenile for a random adult male's genes in locus $G_2$, $\gamma_{2\mathrm{f}}$ and $\gamma_{2\mathrm{m}}$ is the mapping between genotype and phenotype for an adult female or male respectively. According to our assumption that locus $G_2$ would only take an effect if its carrier is a male, thus $\gamma_{2\mathrm{f}} = 0$, $\gamma_{2\mathrm{m}} = 1$. Then $\mathrm{d}W_{\mathrm{f}}/\mathrm{d}g_{2\mathrm{f}}$ can be simplified to

$$\frac{\mathrm{d}W}{\mathrm{d}g_2} = \frac{\partial W}{\partial x_{\mathrm{Fa}}}p_{\mathrm{OF}} + \frac{\partial W}{\partial y_{\mathrm{Fa}}}p_{\mathrm{JU}} \quad \text{(S53)}$$

Using the same way as deriving the optimal value of locus $G_1$, $z_{\mathrm{f}}^*$, we could obtain the optimal value of left-handedness $z_{\mathrm{m}}^*$ for all the loci that only control handedness when they are carried by males:

$$z_{\mathrm{m}}^* = \frac{1}{2}\frac{b_{\mathrm{m}}(r_{\mathrm{OF}} - r_{\mathrm{JU}})}{(1 + b_{\mathrm{m}})r_{\mathrm{OF}} - b_{\mathrm{m}}r_{\mathrm{JU}}} \quad \text{(S54)}$$

*1.62 | Convergence stable strategy*

Combining with parent-of-origin effects, we can write the optimal value of left-handedness for all the loci that control female’s handedness from the perspective of maternal-origin genes, $z_{\mathrm{fM}}^*$, and that from the perspective of paternal-origin genes, $z_{\mathrm{fP}}^*$, as well as the optimal value of left-handedness for all the loci that control male’s handedness from the perspective of maternal-origin genes and paternal-origin genes respectively: $z_{\mathrm{mM}}^*$ and $z_{\mathrm{mP}}^*$:

$$z_{\mathrm{fM}}{}^{*} = \frac{1}{2}\frac{b_{\mathrm{f}}\left(r_{\mathrm{OM|-M}} - r_{\mathrm{JA|-M}}\right)}{(1+b_{\mathrm{f}})r_{\mathrm{OM|-M}} - b_{\mathrm{f}}r_{\mathrm{JA|-M}}} \quad \text{(S55)}$$

$$z_{\mathrm{fP}}{}^{*} = \frac{b_{\mathrm{f}}\left(r_{\mathrm{OM|-P}} - r_{\mathrm{JA|-P}}\right)}{(1+b_{\mathrm{f}})r_{\mathrm{OM|-P}} - b_{\mathrm{f}}r_{\mathrm{JA|-P}}} \quad \text{(S56)}$$

$$z_{\mathrm{mM}}{}^{*} = \frac{1}{2}\frac{b_{\mathrm{m}}\left(r_{\mathrm{OF|-M}} - r_{\mathrm{JU|-M}}\right)}{(1+b_{\mathrm{m}})r_{\mathrm{OF|-M}} - b_{\mathrm{m}}r_{\mathrm{JU|-M}}} \quad \text{(S57)}$$

$$z_{\mathrm{mP}}{}^{*} = \frac{1}{2}\frac{b_{\mathrm{m}}\left(r_{\mathrm{OF|-P}} - r_{\mathrm{JU|-P}}\right)}{(1+b_{\mathrm{m}})r_{\mathrm{OF|-P}} - b_{\mathrm{m}}r_{\mathrm{JU|-P}}} \quad \text{(S58)}$$

where $r_{\mathrm{OM|-P}} = p_{\mathrm{OM|-P}}/p_{\mathrm{I}}$, $r_{\mathrm{OF|-P}} = p_{\mathrm{OF|-P}}/p_{\mathrm{I}}$, $r_{\mathrm{JA|-P}} = p_{\mathrm{JA|-P}}/p_{\mathrm{I}}$, $r_{\mathrm{JU|-P}} = p_{\mathrm{JU|-P}}/p_{\mathrm{I}}$. Substituting all the relatedness in expressions (S51), (S54) and (S55-(S58), we obtain the optimal values of left-handedness when it is involved in within-group combat:

$$z_{\mathrm{f}}{}^{*} = ((b_{\mathrm{f}}(n-1)(H_{\mathrm{f}} - H_{\mathrm{m}} - 4(2-\bar{m})\bar{m}n))) \,/\, ((-8n + 2(n - 1)(-2b_{\mathrm{f}}\Delta m(1-\bar{m}) - 4(1+b_{\mathrm{f}})(2-\bar{m})\bar{m}n))) \quad \text{(S59)}$$

$$\begin{aligned} z_{\mathrm{fM}}{}^{*} = {} & ((b_{\mathrm{f}}(-2\Delta m(1-m_{\mathrm{f}})^2(1-\bar{m}) + 4\Delta m(2+H_{\mathrm{f}})(1-\bar{m})n \\ & + \left(m_{\mathrm{f}}\left(2 + m_{\mathrm{f}}(5 + H_{\mathrm{f}} - 2m_{\mathrm{f}})\right) + 2(7 + H_{\mathrm{f}} - 2m_{\mathrm{f}})m_{\mathrm{m}} \right. \\ & \left. - (5+m_{\mathrm{f}})m_{\mathrm{m}}{}^2\right)n^2 - 8(2-\bar{m})\bar{m}n^3))) \\ & /\, ((-4b_{\mathrm{f}}\Delta m(1-m_{\mathrm{f}})^2(1-\bar{m}) + 4\Delta m(1-\bar{m})(m_{\mathrm{f}} - 1 + 2b_{\mathrm{f}}(2 \\ & + H_{\mathrm{f}}) + m_{\mathrm{m}} - M)n + 2(-8 + m_{\mathrm{f}}(10 + H_{\mathrm{f}} - 3m_{\mathrm{f}} + b_{\mathrm{f}}(2 + m_{\mathrm{f}}(5 \\ & + H_{\mathrm{f}} - 2m_{\mathrm{f}}))) + 6m_{\mathrm{m}} + (2b_{\mathrm{f}}(7 + H_{\mathrm{f}} - 2m_{\mathrm{f}}) - m_{\mathrm{f}}(4 + H_{\mathrm{f}} - m_{\mathrm{f}}))m_{\mathrm{m}} \\ & - (3m_{\mathrm{f}} - 1 + b_{\mathrm{f}}(5 + H_{\mathrm{f}}))m_{\mathrm{m}}{}^2 - (1-m_{\mathrm{f}})m_{\mathrm{m}}{}^3)n^2 - 16(1+b_{\mathrm{f}})(2 \\ & - \bar{m})\bar{m}n^3)) \end{aligned} \quad \text{(S60)}$$

$$\begin{aligned} z_{\mathrm{fP}}{}^{*} = {} & ((b_{\mathrm{f}}(-2\Delta m(H_{\mathrm{f}}+1)(1-\bar{m}) + 4H_{\mathrm{f}}\Delta m(1-\bar{m})n + ((H_{\mathrm{f}} - m_{\mathrm{f}})(2 + H_{\mathrm{f}} + m_{\mathrm{f}}) \\ & + 2(m_{\mathrm{f}}{}^2 - 5)m_{\mathrm{m}} - (H_{\mathrm{f}} - 3)m_{\mathrm{m}}{}^2)n^2 - 8(\bar{m} - 2)\bar{m}n^3))) \\ & /\, ((-4b_{\mathrm{f}}\Delta m(1-m_{\mathrm{f}})^2(1-\bar{m}) + 4\Delta m(1-\bar{m})(2\bar{m} - 1 + 2b_{\mathrm{f}}H_{\mathrm{f}} \\ & - M)n + 2(8 + (H_{\mathrm{f}} - m_{\mathrm{f}})(2 + m_{\mathrm{f}} + b_{\mathrm{f}}(2 + H_{\mathrm{f}} + m_{\mathrm{f}})) - 10m_{\mathrm{m}} \\ & + (-(H_{\mathrm{f}} - 2m_{\mathrm{f}})(1 + m_{\mathrm{f}}) + 2b_{\mathrm{f}}(m_{\mathrm{f}}{}^2 - 5))m_{\mathrm{m}} + (5 - 3m_{\mathrm{f}} - \mathrm{b}_{\mathrm{f}}(H_{\mathrm{f}} \\ & - 3))m_{\mathrm{m}}{}^2 - (1-m_{\mathrm{f}})m_{\mathrm{m}}{}^3)n^2 + 16(1+b_{\mathrm{f}})(2-\bar{m})\bar{m}n^3)) \end{aligned} \quad \text{(S61)}$$

$$z_{\mathrm{m}}{}^{*} = ((b_{\mathrm{m}}(n-1)(H_{\mathrm{m}} - H_{\mathrm{f}} - 4(2-\bar{m})\bar{m}n))) \,/\, ((-8n + 2(n - 1)(2b_{\mathrm{m}}\Delta m(1-\bar{m}) - 4(1+b_{\mathrm{m}})(2-\bar{m})\bar{m}n))) \tag{S62}$$

$$z_{\mathrm{mM}}{}^{*} = ((2b_{\mathrm{m}}(-\Delta m(1-m_{\mathrm{m}})^2(1-\bar{m}) + 4H_{\mathrm{m}}\Delta m(1-\bar{m})n + \left(m_{\mathrm{f}}{}^2(H_{\mathrm{m}}-3) - (H_{\mathrm{m}} - m_{\mathrm{m}})(2 + H_{\mathrm{m}} + m_{\mathrm{m}}) - 2m_{\mathrm{f}}(m_{\mathrm{m}}{}^2 - 5)\right)n^2 - 8(2-\bar{m})\bar{m}n^3))) \,/\, ((-4b_{\mathrm{m}}\Delta m(1-m_{\mathrm{m}})^2(1-\bar{m}) + 4\Delta m(1 - \bar{m})(2\bar{m} - 1 - M + 2b_{\mathrm{m}}H_{\mathrm{m}})n + 2(-8 - m_{\mathrm{f}}{}^3(m_{\mathrm{m}} - 1) + m_{\mathrm{f}}{}^2(-5 + 3m_{\mathrm{m}} + b_{\mathrm{m}}(H_{\mathrm{m}} - 3)) - (H_{\mathrm{m}} - m_{\mathrm{m}})(2 + m_{\mathrm{m}} + b_{\mathrm{m}}(2 + H_{\mathrm{m}} + m_{\mathrm{m}})) + m_{\mathrm{f}}(10 + m_{\mathrm{m}}(H_{\mathrm{m}} - m_{\mathrm{m}} - 4) - 2b_{\mathrm{m}}(-5 + m_{\mathrm{m}}{}^2)))n^2 - 16(1 + b_{\mathrm{m}})(2-\bar{m})\bar{m}n^3)) \tag{S63}$$

$$z_{\mathrm{mP}}{}^{*} = ((-2b_{\mathrm{m}}\Delta m(n-1)(-(1-m_{\mathrm{m}})^2(1-\bar{m}) - 2\Delta m(1-\bar{m})(3+H_{\mathrm{m}})n + 8(2 - \bar{m})\bar{m}n^2))) \,/\, ((-4b_{\mathrm{m}}\Delta m(n-1)(-(H_{\mathrm{m}}+1)(1-\bar{m}) - 2\Delta m(1-\bar{m})(3+H_{\mathrm{m}})n + 8(2-\bar{m})\bar{m}n^2) + 2n(-2\Delta m(M - 2\bar{m} + 1)(1-\bar{m}) - 2(1-\bar{m})(2\bar{m} - 4 + 2m_{\mathrm{m}} + 2\Delta m\bar{m} - M\Delta m)n + 8(2 - \bar{m})\bar{m}n^2))) \tag{S64}$$

where $\Delta m = m_{\mathrm{f}} - m_{\mathrm{m}}$, $\bar{m} = (m_{\mathrm{f}} + m_{\mathrm{m}})/2$, $M = m_{\mathrm{f}}m_{\mathrm{m}}$, $\Delta b = b_{\mathrm{f}} - b_{\mathrm{m}}$, $\bar{b} = (b_{\mathrm{f}} + b_{\mathrm{m}})/2$, $H_{\mathrm{f}} = (m_{\mathrm{f}} - 2)m_{\mathrm{f}}$, $H_{\mathrm{m}} = (m_{\mathrm{m}} - 2)m_{\mathrm{m}}$. To plot $z_{\mathrm{f}}{}^{*}$ and $z_{\mathrm{m}}{}^{*}$ (Figure 1*b*) we set the female dispersal rate $m_{\mathrm{f}}$ to be 0.5, the relative importance of combat relative to all types of competition for the female $b_{\mathrm{f}}$ and male $b_{\mathrm{m}}$ both to be 1, and number of the number of individuals each sex born in the same patch $n$ to be 5.

### 1.7 | Parental genetic effects

#### *1.71 | Marginal fitness and evolutionary equilibrium*

Now we consider the parental effects, i.e. the effect on the phenotype of the parents of the focal juvenile is caused by the genes carried by the grandparents of the focal juvenile, regardless of the parents' genotype. In this section, the fitness function and relatedness remain the same as previous ones, while the conditions that favours the increase of left-handedness change according to specific situations. Depending on whether there is difference between maternal and paternal effects, and/or between the parental effects on daughters versus those on sons, there can be nine situations: 1) When both parents control the parental effect and all offspring experience the parental effect in their handedness (we denote the optima for left-handedness as $z_{PO}^*$). 2) When both parents control the parental effect and only daughters experience the parental effect in their handedness ($z_{PD}^*$). 3) When both parents control the parental effect and only sons experience the parental effect in their handedness ($z_{PS}^*$). 4) When only mother controls the parental effect and all offspring experience the parental effect in their handedness ($z_{MO}^*$). 5) When only mother controls the parental effect and only daughters experience the parental effect in their handedness ($z_{MD}^*$). 6) When only mother controls the parental effect and only sons experience the parental effect in their handedness ($z_{MS}^*$). 7) When only father controls the parental effect and all offspring experience the parental effect in their handedness ($z_{FO}^*$). 8) When only father controls the parental effect and only daughters experience the parental effect in their handedness ($z_{FD}^*$). 9) When only father controls the parental effect and only sons experience the parental effect in their handedness ($z_{FS}^*$).

*1) Parental control of offspring phenotype* ($z_{PO}^*$)

We consider there is only locus G controlling the phenotype of handedness, and there is no difference in who carries the genes influence the phenotype of offspring, and it affects the handedness phenotype of daughters and sons in the same way. We denote the genic value as $g_f$ and $g_m$ for the juvenile females and males, $G_f$ and $G_m$ for the breeding value for the maternal grandparent and paternal grandparent of the focal juvenile respectively, $G'_f$ for the breeding value of the parent of a random adult in the focal juvenile's mother's group, $G'_m$ for the breeding value of the parent of a random adult in the focal juvenile's father's group. The relationship between the phenotype and genotype can be described as:

$$\frac{dW}{dg}=\frac{\partial W}{\partial x_{\mathrm{Mo}}}\frac{dx_{\mathrm{Mo}}}{dG_{\mathrm{f}}}\frac{dG_{\mathrm{f}}}{dg}+\frac{\partial W}{\partial y_{\mathrm{Mo}}}\frac{dy_{\mathrm{Mo}}}{dG_{\mathrm{f}}'}\frac{dG_{\mathrm{f}}'}{dg}+\frac{\partial W}{\partial x_{\mathrm{Fa}}}\frac{dx_{\mathrm{Fa}}}{dG_{\mathrm{m}}}\frac{dG_{\mathrm{m}}}{dg}+\frac{\partial W}{\partial y_{\mathrm{Fa}}}\frac{dy_{\mathrm{Fa}}}{dG_{\mathrm{m}}'}\frac{dG_{\mathrm{m}}'}{dg}$$

$$=\left(\frac{\partial W}{\partial x_{\mathrm{Mo}}}p_{\mathrm{JMGP}}+\frac{\partial W}{\partial y_{\mathrm{Mo}}}p_{\mathrm{JMAP}}\right)\gamma_{\mathrm{Pf}} \tag{S65}$$

$$+\left(\frac{\partial W}{\partial x_{\mathrm{Fa}}}r_{\mathrm{JPGP}}+\frac{\partial W}{\partial y_{\mathrm{Fa}}}r_{\mathrm{JPUP}}\right)\gamma_{\mathrm{Pm}}$$

where $p_{\mathrm{JMGP}}$ is the consanguinity between the focal juvenile female and its maternal grandparent (here we treat the maternal grandparent as a "tetraploidy"), $p_{\mathrm{JMAP}}$ is the coefficient of the consanguinity between the focal juvenile female and the parent of a random adult female (here “A” denotes "Aunt") in the focal juvenile's  mother's group, $p_{\mathrm{JPGP}}$ is the coefficient of the consanguinity between the focal juvenile female and its paternal grandparent, $p_{\mathrm{JPUP}}$ is the coefficient of the consanguinity between the focal juvenile female and the parent of a random adult male (here “U” denotes "Uncle") in the focal juvenile's father's group, $\gamma_{\mathrm{Pf}}=\frac{dx_{\mathrm{Mo}}}{dG_{\mathrm{f}}}=\frac{dy_{\mathrm{Mo}}}{dG_{\mathrm{f}}'}$ is the mapping between the gene of parents and its expressed phenotype in a female offspring, $\gamma_{\mathrm{Pm}}=\frac{dx_{\mathrm{Fa}}}{dG_{\mathrm{m}}}=\frac{dy_{\mathrm{Fa}}}{dG_{\mathrm{m}}'}$ is the mapping between the gene of parents and its expressed phenotype in a male offspring, and under our assumption $\gamma_{\mathrm{Pf}}$ $=\gamma_{\mathrm{Pm}}=1$. The condition that favours the increase of left-handedness is:

$$\frac{\partial W_{\mathrm{f}}}{\partial x_{\mathrm{Mo}}}r_{\mathrm{JMGP}}+\frac{\partial W_{\mathrm{f}}}{\partial y_{\mathrm{Mo}}}r_{\mathrm{JMAP}}+\frac{\partial W_{\mathrm{f}}}{\partial x_{\mathrm{Fa}}}r_{\mathrm{JPGP}}+\frac{\partial W_{\mathrm{f}}}{\partial y_{\mathrm{Fa}}}r_{\mathrm{JPUP}}>0 \tag{S66}$$

where $r_{\mathrm{JMGP}}=p_{\mathrm{JMGP}}/p_{\mathrm{I}}$, $r_{\mathrm{JMAP}}=p_{\mathrm{JMAP}}/p_{\mathrm{I}}$, $r_{\mathrm{JPGP}}=p_{\mathrm{JPGP}}/p_{\mathrm{I}}$, $r_{\mathrm{JPUP}}=p_{\mathrm{JPUP}}/p_{\mathrm{I}}$. Letting the LHS of expression (S66) be $f(z)$, $f'(z)<0$ is true for all the values of $z$, hence at evolutionary equilibrium if there is intermediate level of left-handedness $z_{PO}{}^{*}$ that satisfies $f(z_{PO}{}^{*})=0$, we obtain the optimum of left-handedness from the perspective of parent’s genes:

$$z_{\mathrm{PO}}{}^{*}=\frac{1}{2}\left(1-\frac{r_{\mathrm{JMAP}}+r_{\mathrm{JPGP}}}{r_{\mathrm{JMGP}}+b_{\mathrm{f}}\left(-r_{\mathrm{JMAP}}+r_{\mathrm{JMGP}}\right)+r_{\mathrm{JPGP}}+b_{\mathrm{m}}r_{\mathrm{JPGP}}-b_{\mathrm{m}}r_{\mathrm{JPUP}}}\right) \tag{S67}$$

if we set $b_{\mathrm{f}} = b_{\mathrm{m}} = 1$, expression (S67) can be re-written as: $\frac{1}{2}+\frac{1}{2}\frac{1}{\frac{p_{J\mathrm{AveAUP}}}{p_{J\mathrm{AveGP}}}-2}$, where $p_{\mathrm{AveAUP}}$ is the consanguinity between an individual and the parent of the individual's parent's social partner, and $p_{\mathrm{AveAUP}} = 1/2\ (p_{\mathrm{JMAP}} + p_{\mathrm{JPUP}})$, $p_{\mathrm{AveGP}}$ is the consanguinity between an individual and its grandparent, and $p_{\mathrm{AveGP}} = 1/2\ (p_{\mathrm{JMGP}} + p_{\mathrm{JPGP}})$. If we set $b_{\mathrm{f}} = b_{\mathrm{m}} = 1$, expression (S8) can be re-written as: $\frac{1}{2}+\frac{1}{2}\frac{1}{\frac{p_J}{p_J}-2}$. We use ratio $r_1 = p_{\mathrm{AveAUP}}/p_{\mathrm{AveGP}}$ for considering the optima from the perspective of parents, and $r_2 = p_{\mathrm{J}}/p_{\mathrm{O}}$ for considering the optimum from the perspective of the offspring. As $r_1$ is always greater than $r_2$, parents always favour a lower value of left-handedness in their offspring than the offspring would, in the context of within-group combat.

*2) Parental control of daughter's phenotype* ($z_{\mathrm{PD}}^*$)

Under our assumption that only daughters experience parental effect, $\gamma_{\mathrm{Pf}} = 1$, $\gamma_{\mathrm{Pm}} = 0$. The condition that favours the increase of left-handedness is

$$\frac{\partial W}{\partial x_{\mathrm{Mo}}} r_{\mathrm{JMGP}} + \frac{\partial W}{\partial y_{\mathrm{Mo}}} r_{\mathrm{JMAP}} > 0 \qquad \text{(S68)}$$

with similar process of obtaining $z_{PO}{}^*$ we obtain the optimal value of left-handedness from the perspective of parent's genes to its daughter

$$z_{\mathrm{PD}}{}^* = \frac{1}{2}\frac{b_{\mathrm{f}}(r_{\mathrm{JMAP}} - r_{\mathrm{JMGP}})}{b_{\mathrm{f}} r_{\mathrm{JMAP}} - (1 + b_{\mathrm{f}}) r_{\mathrm{JMGP}}} \qquad \text{(S69)}$$

*3) Parental control of son's phenotype* ($z_{\mathrm{PS}}^*$)

Under our assumption that only daughters experience parental effect, $\gamma_{\mathrm{Pf}} = 0$, $\gamma_{\mathrm{Pm}} = 1$. The condition that favours the increase of left-handedness is:

$$\frac{\partial W}{\partial x_{\mathrm{Fa}}} r_{\mathrm{JPGP}} + \frac{\partial W}{\partial y_{\mathrm{Fa}}} r_{\mathrm{JPUP}} > 0 \qquad \text{(S70)}$$

with similar process, we obtain the optimal value of left-handedness from the perspective of parent's genes to its son:

$$z_{\mathrm{PS}}{}^{*} = \frac{1}{2}\frac{b_{\mathrm{m}}(r_{\mathrm{JPGP}} - r_{\mathrm{JPUP}})}{r_{\mathrm{JPGP}} + b_{\mathrm{m}}r_{\mathrm{JPGP}} - b_{\mathrm{m}}r_{\mathrm{JPUP}}} \tag{S71}$$

*4) Maternal control of offspring phenotype* ($z_{\mathrm{MO}}{}^{*}$)

In this case, the relationship between phenotype and genotype is

$$\frac{dW}{dg} = \left(\frac{\partial W}{\partial x_{\mathrm{Mo}}}p_{\mathrm{JMGM}} + \frac{\partial W}{\partial y_{\mathrm{Mo}}}p_{\mathrm{JMAM}}\right)\gamma_{\mathrm{Ff}} + \left(\frac{\partial W}{\partial x_{\mathrm{Fa}}}p_{\mathrm{JPGM}} + \frac{\partial W}{\partial y_{\mathrm{Fa}}}p_{\mathrm{JPUM}}\right)\gamma_{\mathrm{Fm}} \tag{S72}$$

where $p_{\mathrm{JMGM}}$ is the consanguinity between the focal juvenile female and its maternal grandmother, $p_{\mathrm{JMAM}}$ is the consanguinity between the focal juvenile female and the mother of a random adult female in the focal juvenile's mother's group, $p_{\mathrm{JPGM}}$ is the consanguinity between the focal juvenile female and its paternal grandmother, $p_{\mathrm{JPUM}}$ is the consanguinity between the focal juvenile female and the mother of a random adult male in the focal juvenile's father's group. $\gamma_{\mathrm{Ff}}$ is the mapping between the gene of mother and its expressed phenotype in a female offspring, $\gamma_{\mathrm{Fm}}$ is the mapping between the gene of mother and its expressed phenotype in a male offspring. Under our assumption that all offspring experience parental effect, $\gamma_{\mathrm{Ff}} = \gamma_{\mathrm{Fm}} = \gamma$. The condition that favours the increase of left-handedness is

$$\frac{\partial W}{\partial x_{\mathrm{Mo}}}r_{\mathrm{JMGM}} + \frac{\partial W}{\partial y_{\mathrm{Mo}}}r_{\mathrm{JMAM}} + \frac{\partial W}{\partial x_{\mathrm{Fa}}}p_{\mathrm{JPGM}} + \frac{\partial W}{\partial y_{\mathrm{Fa}}}p_{\mathrm{JPUM}} > 0 \tag{S73}$$

where $r_{\mathrm{JMGM}} = p_{\mathrm{JMGM}}/p_{\mathrm{I}}$, $r_{\mathrm{JMAM}} = p_{\mathrm{JMAM}}/p_{\mathrm{I}}$, $r_{\mathrm{JPGM}} = p_{\mathrm{JPGM}}/p_{\mathrm{I}}$, $r_{\mathrm{JPUM}} = p_{\mathrm{JPUM}}/p_{\mathrm{I}}$. With similar process as previous situations, we obtain the optimal value of left-handedness from the perspective of mother's genes to her offspring

$$z_{\mathrm{MO}}{}^{*} = \frac{1}{2}\left(1 - \frac{r_{\mathrm{JMGM}} + r_{\mathrm{JPGM}}}{r_{\mathrm{JMGM}} + b_{\mathrm{f}}(r_{\mathrm{JMGM}} - r_{\mathrm{JMAM}}) + r_{\mathrm{JPGM}} + b_{\mathrm{m}}r_{\mathrm{JPGM}} - b_{\mathrm{m}}r_{\mathrm{JPUM}}}\right) \tag{S74}$$

*5) Maternal control of daughter's phenotype* ($z_{\mathrm{MD}}{}^{*}$)

By changing $\gamma_{\mathrm{Ff}}$ to 1, $\gamma_{\mathrm{Fm}}$ to 0, we could obtain the condition for an increase in left-handedness to be favoured

$$\frac{\partial W}{\partial x_{\mathrm{Mo}}} r_{\mathrm{JMGM}} + \frac{\partial W}{\partial y_{\mathrm{Mo}}} r_{\mathrm{JMAM}} > 0 \qquad (S75)$$

With similar process, we obtain the optimal value of left-handedness from the perspective of mother's genes to her daughters

$$z_{\mathrm{MD}}{}^{*} = \frac{1}{2} \frac{b_{\mathrm{f}}\left(r_{\mathrm{JMAM}} - r_{\mathrm{JMGM}}\right)}{b_{\mathrm{f}} r_{\mathrm{JMAM}} - (1 + b_{\mathrm{f}}) r_{\mathrm{JMGM}}} \qquad (S76)$$

*6) Maternal control of son's phenotype* ($z_{\mathrm{MS}}{}^{*}$)

By changing $\gamma_{\mathrm{Ff}}$ to 0, $\gamma_{\mathrm{Fm}}$ to 1, we could obtain the condition for an increase in left-handedness to be favoured

$$\frac{\partial W}{\partial x_{\mathrm{Fa}}} r_{\mathrm{JPGM}} + \frac{\partial W}{\partial y_{\mathrm{Fa}}} r_{\mathrm{JPUM}} > 0 \qquad (S77)$$

With similar process, we obtain the optimal value of left-handedness from the perspective of mother's genes to her sons

$$z_{\mathrm{MS}}{}^{*} = \frac{1}{2} \frac{b_{\mathrm{m}}\left(r_{\mathrm{JPGM}} - r_{\mathrm{JPUM}}\right)}{r_{\mathrm{JPGM}} + b_{\mathrm{m}} r_{\mathrm{JPGM}} - b_{\mathrm{m}} r_{\mathrm{JPUM}}} \qquad (S78)$$

*7) Paternal control of offspring phenotype* ($z_{\mathrm{FO}}{}^{*}$)

In this case, the relationship between phenotype and genotype is

$$\frac{dW}{dg} = \left(\frac{\partial W}{\partial x_{\mathrm{Mo}}} p_{\mathrm{JMGF}} + \frac{\partial W}{\partial y_{\mathrm{Mo}}} p_{\mathrm{JMAF}}\right) \gamma_{\mathrm{Mf}} + \left(\frac{\partial W}{\partial x_{\mathrm{Fa}}} p_{\mathrm{JPGF}} + \frac{\partial W}{\partial y_{\mathrm{Fa}}} p_{\mathrm{JPUF}}\right) \gamma_{\mathrm{Mm}} \qquad (S79)$$

where $p_{\mathrm{JMGF}}$ is the consanguinity between the focal juvenile female and its maternal grandfather, $p_{\mathrm{JMAF}}$ is the consanguinity between the focal juvenile female and the father of a random adult female in its mother's group, $p_{\mathrm{JPGF}}$ is the consanguinity between the focal juvenile female and its paternal grandfather, $p_{\mathrm{JPUF}}$ is the consanguinity between the focal juvenile female and the father of a random adult male in its father's group, $\gamma_{\mathrm{Mf}}$ is the mapping between the gene of father and its expressed phenotype in a female offspring, $\gamma_{\mathrm{Mm}}$ is the mapping between the gene of parents and its expressed phenotype in a male offspring. Under our assumption that all offspring experience parental effect, $\gamma_{\mathrm{Mf}} = \gamma_{\mathrm{Mm}} = \gamma$. The condition that favours the increase of left-handedness is

$$\frac{\partial W}{\partial x_{\mathrm{Mo}}} r_{\mathrm{JMGF}} + \frac{\partial W}{\partial y_{\mathrm{Mo}}} r_{\mathrm{JMAF}} + \frac{\partial W}{\partial x_{\mathrm{Fa}}} r_{\mathrm{JPGF}} + \frac{\partial W}{\partial y_{\mathrm{Fa}}} r_{\mathrm{JPUF}} > 0 \tag{S80}$$

where $r_{\mathrm{JMGF}} = p_{\mathrm{JMGF}}/p_{\mathrm{I}}$, $r_{\mathrm{JPGF}} = p_{\mathrm{JPGF}}/p_{\mathrm{I}}$, $r_{\mathrm{JMAF}} = p_{\mathrm{JMAF}}/p_{\mathrm{I}}$, $r_{\mathrm{JPUF}} = p_{\mathrm{JPUF}}/p_{\mathrm{I}}$. With similar process as previous situations, we obtain the optimal value of left-handedness from the perspective of father's genes to his offspring

$$z_{\mathrm{FO}}{}^{*} = \frac{1}{2}\left(1 - \frac{r_{\mathrm{JMGF}} + r_{\mathrm{JPGF}}}{r_{\mathrm{JMGF}} + b_{\mathrm{f}}\left(r_{\mathrm{JMGF}} - r_{\mathrm{JMAF}}\right) + r_{\mathrm{JPGF}} + b_{\mathrm{m}} r_{\mathrm{JPGF}} - b_{\mathrm{m}} r_{\mathrm{JPUF}}}\right) \tag{S81}$$

*8) Paternal control of daughter's phenotype* ($z_{\mathrm{FD}}{}^{*}$)

By changing $\gamma_{\mathrm{Mf}}$ to 1, $\gamma_{\mathrm{Mm}}$ to 0, we could obtain the condition for an increase in left-handedness to be favoured

$$\frac{\partial W}{\partial x_{\mathrm{Mo}}} r_{\mathrm{JMGF}} + \frac{\partial W}{\partial y_{\mathrm{Mo}}} r_{\mathrm{JMAF}} > 0 \tag{S82}$$

With similar process, we obtain the optimal value of left-handedness from the perspective of father's genes to his daughters

$$z_{\mathrm{FD}}{}^{*} = \frac{1}{2} \frac{b_{\mathrm{f}}\left(r_{\mathrm{JMAF}} - r_{\mathrm{JMGF}}\right)}{b_{\mathrm{f}} r_{\mathrm{JMAF}} - (1 + b_{\mathrm{f}}) r_{\mathrm{JMGF}}} \tag{S83}$$

*9) Paternal control of son's phenotype* ($z_{\mathrm{FS}}{}^{*}$)

By changing $\gamma_{\mathrm{Mf}}$ to 0, $\gamma_{\mathrm{Mm}}$ to 1, we could obtain the condition for an increase in left-handedness to be favoured

$$\frac{\partial W}{\partial x_{\mathrm{Fa}}} r_{\mathrm{JPGF}} + \frac{\partial W}{\partial y_{\mathrm{Fa}}} r_{\mathrm{JPUF}} > 0 \tag{S84}$$

With similar process, we obtain the optimal value of left-handedness from the perspective of father's genes to his sons

$$z_{\mathrm{FS}}{}^{*} = \frac{1}{2}\frac{b_{\mathrm{m}}\left(r_{\mathrm{JPGF}} - r_{\mathrm{JPUF}}\right)}{r_{\mathrm{JPGF}} + b_{\mathrm{m}}r_{\mathrm{JPGF}} - b_{\mathrm{m}}r_{\mathrm{JPUF}}} \tag{S85}$$

*1.72 | Relatedness*

The consanguinity between the focal juvenile and its maternal grandmother $p_{\mathrm{JMGM}}$ is

$$\begin{aligned} p_{\mathrm{JMGM}} = {} & \frac{1}{2}\left(\frac{1}{2}p_{\mathrm{I}} + \frac{1}{2}f'\right) \\ & + \frac{1}{2}(1 - m_{\mathrm{f}})(1 - m_{\mathrm{m}})\left(\frac{1}{n}\left(\frac{1}{2}p_{\mathrm{I}} + \frac{1}{2}f'\right)\right. \\ & \left. + \frac{n-1}{n}\left(\frac{1}{2}(1 - m_{\mathrm{f}})^2 p_{\mathrm{x}}{}' + \frac{1}{2}f'\right)\right) \end{aligned} \tag{S86}$$

That is: with probability 1/2 the gene we pick comes from the juvenile's mother, in which case the consanguinity is that between the mother and the maternal grandmother, which is with probability 1/2 the gene comes from the maternal grandmother, and the consanguinity is that between the maternal grandmother and herself i.e. $p_{\mathrm{I}}$, and with probability 1/2 the gene comes from the maternal grandfather, and the consanguinity is that between mating partners i.e. *f'*, and with probability 1/2 that the gene we pick comes from the juvenile's father, in which case the consanguinity is that between the juvenile's father and the maternal grandmother, which is with probability $(1 - m_{\mathrm{f}})(1 - m_{\mathrm{m}})$ neither the mother nor the father disperses from their natal patch, and with probability 1/*n* the mother and the father share one mother, and with probability 1/2 the gene comes from their mother, and the consanguinity is $p_{\mathrm{I}}$, and with probability 1/2 the gene comes from their father, and the consanguinity is that between two random mating partner i.e. *f'*, and with probability (*n*-1)/*n* the mother and the father do not share one mother, and with probability 1/2 the gene comes from the paternal grandmother, with probability $(1 - m_{\mathrm{f}})^2$ neither of the two females disperses, and the consanguinity is that between two random juveniles born in the same patch i.e. $p_x'$, and with probability 1/2 the gene comes from the paternal grandfather, and the consanguinity is $f'$. The consanguinity between the focal juvenile and its maternal grandfather $p_{\mathrm{JMGF}}$ is

$$p_{\mathrm{JMGF}} = \frac{1}{2}\left(\frac{1}{2}f' + \frac{1}{2}p_{\mathrm{I}}\right) + \frac{1}{2}(1 - m_{\mathrm{f}})(1 - m_{\mathrm{m}})\left(\frac{1}{n}\left(\frac{1}{2}f' + \frac{1}{2}p_{\mathrm{I}}\right) + \frac{n-1}{n}\left(\frac{1}{2}f' + \frac{1}{2}(1 - m_{\mathrm{m}})^2 p_{\mathrm{x}}'\right)\right) \tag{S87}$$

That is: with probability 1/2 the gene we pick comes from the juvenile's mother, in which case the consanguinity is that between the mother and her father, which is with probability 1/2 the gene we pick comes from the maternal grandmother, and the consanguinity is that between mating partners i.e. $f'$, and with probability 1/2 the gene we pick comes from the maternal grandfather, and the consanguinity is that between the grandfather and himself $p_{\mathrm{I}}$, and with probability 1/2 the gene we pick comes from the juvenile's father, in which case the consanguinity is that between the juvenile's father and maternal grandfather, which is with probability $(1 - m_{\mathrm{f}})(1 - m_{\mathrm{m}})$ neither the mother nor the father disperses, and with probability 1/*n* the mother and the father share one father, with probability 1/2 the gene we pick comes from their mother, and the consanguinity is that between two random mating partner i.e. $f'$, and with probability 1/2 the gene we pick comes from their father, and the consanguinity is $p_{\mathrm{I}}$, and with probability (*n*-1)/*n* the mother and the father do not share one father, with probability 1/2 the gene we pick comes from the paternal mother, and the consanguinity is that between two random mating partners $f'$, and with probability 1/2 that the genes we pick come from the paternal father, with probability $(1 - m_{\mathrm{m}})^2$ neither of the two males disperses, and the consanguinity is that between two random juveniles born in the same patch i.e. $p_x'$. The consanguinity between the focal juvenile and the mother of a random adult female in its mother's social group $p_{\mathrm{JMAM}}$ is

$$p_{\mathrm{JMAM}} = \frac{1}{2}\left(\frac{1}{n}\left(\frac{1}{2}p_{\mathrm{I}} + \frac{1}{2}f'\right)\right.$$

$$+ \frac{n-1}{n}(1-m_{\mathrm{f}})^2\left(\frac{1}{n}\left(\frac{1}{2}p_{\mathrm{I}} + \frac{1}{2}f'\right)\right.$$

$$\left.\left.+ \frac{n-1}{n}\left(\frac{1}{2}(1-m_{\mathrm{f}})^2 p_{\mathrm{X}}' + \frac{1}{2}f'\right)\right)\right) \tag{S88}$$

$$+ \frac{1}{2}(1-m_{\mathrm{f}})(1-m_{\mathrm{m}})\left(\frac{1}{n}\left(\frac{1}{2}p_{\mathrm{I}} + \frac{1}{2}f'\right)\right.$$

$$\left.+ \frac{n-1}{n}\left(\frac{1}{2}(1-m_{\mathrm{f}})^2 p_{\mathrm{X}}' + \frac{1}{2}f'\right)\right)$$

That is: with probability 1/2 the gene we pick comes from the juvenile's mother, in which case the consanguinity is that between the juvenile's mother and the mother of a random adult female in the juvenile's mother's social group, which is, with probability 1/*n* the random adult female ("aunt" hereafter) is the juvenile's mother, and the consanguinity is that between the juvenile's mother and maternal grandmother which is $\frac{1}{2}p_{\mathrm{I}} + \frac{1}{2}f'$, and with probability (*n*-1)/*n* the aunt is not the juvenile's mother, with the probability $(1-m_{\mathrm{f}})^2$ neither of the two females disperses, and with probability 1/*n* the aunt and the juvenile's mother share one mother, with probability (*n*-1)/*n* the aunt and the juvenile's mother do not share one mother, with probability 1/2 that the mother's gene comes from her mother, with probability $(1-m_{\mathrm{f}})^2$ neither the grandmother nor the mother of the aunt disperses, and the consanguinity is that between two random juvenile born in the same patch i.e. $p_{\mathrm{X}}'$, and with probability 1/2 that the mother's gene came from her father, in which case the consanguinity is that between two random mating partners $f'$, with probability 1/2 the gene we pick comes from the juvenile's father, and with probability $(1-m_{\mathrm{f}})(1-m_{\mathrm{m}})$ neither the aunt nor the father disperses, with probability 1/*n* the aunt and the father share one mother, with probability 1/2 the gene comes from their mother, and the consanguinity is that between the grandmother and herself i.e. $p_{\mathrm{I}}$, and with probability 1/2 the gene comes from the juvenile's paternal grandfather, and the consanguinity is $f'$, and with probability $(n\text{-}1)/n$ the aunt and the father do not share one mother, with probability 1/2 the gene comes from the juvenile's paternal grandmother, with probability $(1-m_\mathrm{f})^2$ neither the mother of the juvenile's aunt nor the paternal grandmother disperses, and the consanguinity is that between two random juveniles born in the same patch $p_\mathrm{x}'$, and with probability 1/2 the gene comes from the juvenile's paternal grandfather, and the consanguinity is $f'$. The consanguinity between the focal juvenile and the father of a random adult female in its mother's group $p_\mathrm{JMAF}$ is

$$p_\mathrm{JMAF} = \frac{1}{2}\Bigg(\frac{1}{n}\left(\frac{1}{2}f' + \frac{1}{2}p_\mathrm{I}\right) + \frac{n-1}{n}(1-m_\mathrm{f})^2\left(\frac{1}{n}\left(\frac{1}{2}f' + \frac{1}{2}p_\mathrm{I}\right) + \frac{n-1}{n}\left(\frac{1}{2}f' + \frac{1}{2}(1-m_\mathrm{m})^2 p_\mathrm{x}'\right)\right)\Bigg) + \frac{1}{2}(1-m_\mathrm{f})(1-m_\mathrm{m})\left(\frac{1}{n}\left(\frac{1}{2}f' + \frac{1}{2}p_\mathrm{I}\right) + \frac{n-1}{n}\left(\frac{1}{2}f' + \frac{1}{2}(1-m_\mathrm{m})^2 p_\mathrm{x}'\right)\right) \quad \text{(S89)}$$

That is: with probability 1/2 the gene we pick comes from the juvenile's mother, in which case the consanguinity is that between the mother and the father of the aunt, which is, with probability $1/n$ the aunt is the juvenile's mother, and with probability 1/2 the gene comes from the juvenile's maternal grandmother, and the consanguinity is $f'$, with probability 1/2 the gene comes from the juvenile's maternal grandfather, and the consanguinity is that of the maternal grandfather to himself $p_\mathrm{I}$, and with probability $(n\text{-}1)/n$ the aunt is not the juvenile's mother, with probability $(1-m_\mathrm{f})^2$ neither of the two females disperses, with probability $1/n$ the aunt and the mother have a same father, with probability 1/2 the gene comes from the mother's mother, and the consanguinity is $f'$, and with probability 1/2 the gene comes from the mother's father, and the consanguinity is $p_\mathrm{I}$, and with probability $(n\text{-}1)/n$ the aunt and the mother do not have a same father, with probability 1/2 the gene comes from the juvenile's maternal grandmother, and the consanguinity is *f'*, and with probability 1/2 the gene comes from the juvenile's grandfather, with probability $(1 - m_\mathrm{m})^2$ neither of the maternal grandfather nor the aunt's father disperses, and the consanguinity is $p_\mathrm{X}$'; and with probability 1/2 that the gene we pick come from the juvenile's father, in which case the consanguinity is that between the father and the father of the aunt, which is, with probability $(1 - m_\mathrm{f})(1 - m_\mathrm{m})$ neither the aunt nor the father disperses, and with probability 1/*n* the aunt and the father share one father, with probability 1/2 the gene comes from the paternal grandmother, and the consanguinity is *f'*, with probability 1/2 the gene comes from the paternal grandfather, and the consanguinity is $p_\mathrm{I}$, and with probability (*n*-1)/*n* the aunt and the father do not share one father, with probability 1/2 the gene comes from the paternal grandmother, and the consanguinity is *f'*, with probability 1/2 the gene comes from the paternal grandfather, with probability $(1 - m_\mathrm{m})^2$ neither of the maternal grandfather nor the aunt's father disperses, and the consanguinity is $p_\mathrm{X}$'. Hence the consanguinity between the focal juvenile and the parent of the aunt $p_\mathrm{JMAP}$ can be given as

$$p_\mathrm{JMAP} = \frac{1}{2} p_\mathrm{JMAM} + \frac{1}{2} p_\mathrm{JMAF} \tag{S90}$$

Similarly, $p_\mathrm{JMGP}$ which is the consanguinity between the focal juvenile and its maternal grandparents, can be given as

$$p_\mathrm{JMGP} = \frac{1}{2} p_\mathrm{JMGM} + \frac{1}{2} p_\mathrm{JMGF} \tag{S91}$$

Now we consider the consanguinity through paternal grandparents. The consanguinity between the focal juvenile and its paternal grandmother $p_\mathrm{JPGM}$ is

$$\begin{aligned} p_\mathrm{JPGM} = &\frac{1}{2}(1 - m_\mathrm{f})(1 - m_\mathrm{m})\left(\frac{1}{n}\left(\frac{1}{2}p_\mathrm{I} + \frac{1}{2}f'\right) + \frac{n-1}{n}\left(\frac{1}{2}(1 - m_\mathrm{f})^2 p_\mathrm{X}' + \frac{1}{2}f'\right)\right) \\ &+ \frac{1}{2}\left(\frac{1}{2}p_\mathrm{I} + \frac{1}{2}f'\right) \end{aligned} \tag{S92}$$

That is: with probability 1/2 the gene we pick comes from the juvenile's mother, in which case the consanguinity is with probability $(1-m_{\mathrm{f}})(1-m_{\mathrm{m}})$ neither the mother nor the father disperses, with probability 1/*n* the mother and the father share one mother, with probability 1/2 the gene comes from the maternal grandmother, and the consanguinity is $p_{\mathrm{I}}$, with probability 1/2 the gene comes from the maternal grandfather, and the consanguinity is $f'$, and with probability (*n*-1)/*n* the mother and the father do not share one mother, with probability 1/2 the gene comes from the maternal grandmother, with probability $(1-m_{\mathrm{f}})^2$ neither of the two females disperses, and the consanguinity is $p_{\mathrm{x}}'$, with probability 1/2 the gene comes from the maternal grandfather, and the consanguinity is $f'$, with probability 1/2 the gene we pick comes from the juvenile's father, in which case the consanguinity is, with probability 1/2 the gene comes from the paternal grandmother, and the consanguinity is $p_{\mathrm{I}}$, with probability 1/2 the gene comes from the paternal grandfather, and the consanguinity is $f'$. The consanguinity between the focal juvenile and its paternal grandfather $p_{\mathrm{JPGF}}$ is

$$p_{\mathrm{JPGF}} = \frac{1}{2}(1-m_{\mathrm{f}})(1-m_{\mathrm{m}})\left(\frac{1}{n}\left(\frac{1}{2}f' + \frac{1}{2}p_{\mathrm{I}}\right) + \frac{n-1}{n}\left(\frac{1}{2}f' + \frac{1}{2}(1-m_{\mathrm{m}})^2 p_{\mathrm{x}}'\right)\right) + \frac{1}{2}\left(\frac{1}{2}f' + \frac{1}{2}p_{\mathrm{I}}\right) \tag{S93}$$

That is: with probability 1/2 the gene we pick comes from the juvenile's mother, in which case the consanguinity is, with probability $(1-m_{\mathrm{f}})(1-m_{\mathrm{m}})$ neither the mother nor the father disperses, and with probability 1/*n* the mother and the father share one mother, with probability 1/2 the gene comes from the maternal grandmother, and the consanguinity is $f'$, with probability 1/2 the gene comes from the maternal grandfather, and the consanguinity is $p_{\mathrm{I}}$, and with probability (*n*-1)/*n* the mother and the father do not share one mother, with probability 1/2 the gene comes from the maternal grandmother, and the consanguinity is $f'$, with probability 1/2 the gene comes from the maternal grandfather, with probability $(1-m_{\mathrm{m}})^2$ neither of the two males disperses, and the consanguinity is $p_{\mathrm{x}}'$, with probability

1/2 the gene we pick comes from the juvenile’s father, in which case the consanguinity is, with probability 1/2 the gene comes from the paternal grandmother, and the consanguinity is $f'$, and with probability 1/2 the gene comes from the paternal grandfather, and the consanguinity is $p_{\mathrm{I}}$. The consanguinity between the focal juvenile and the mother of a random adult male in its father's social group $p_{\mathrm{JPUM}}$ is

$$\begin{aligned} p_{\mathrm{JPUM}} = \frac{1}{2}(1-m_{\mathrm{f}})(1-m_{\mathrm{m}})\left(\frac{1}{n}\left(\frac{1}{2}p_{\mathrm{I}}+\frac{1}{2}f'\right)+\frac{n-1}{n}\left(\frac{1}{2}(1-m_{\mathrm{f}})^2 p_{\mathrm{x}}'+\frac{1}{2}f'\right)\right) \\ +\frac{1}{2}\left(\frac{1}{n}\left(\frac{1}{2}p_{\mathrm{I}}+\frac{1}{2}f'\right)\right. \\ +\frac{n-1}{n}(1-m_{\mathrm{m}})^2\left(\frac{1}{n}\left(\frac{1}{2}p_{\mathrm{I}}+\frac{1}{2}f'\right)\right. \\ \left.\left.+\frac{n-1}{n}\left(\frac{1}{2}(1-m_{\mathrm{f}})^2 p_{\mathrm{x}}'+\frac{1}{2}f'\right)\right)\right) \end{aligned} \qquad (S94)$$

That is: with probability 1/2 the gene we pick comes from the juvenile’s mother, in which case the consanguinity is, with probability $(1-m_{\mathrm{f}})(1-m_{\mathrm{m}})$ neither the mother nor the father’s social partner (“uncle” hereafter) disperses, with probability 1/*n* the mother and the uncle share one mother, with probability 1/2 the gene comes from the maternal grandmother, and the consanguinity is $p_{\mathrm{I}}$, with probability 1/2 the gene comes from the maternal grandfather, and the consanguinity is $f'$, with probability (*n*-1)/*n* the mother and the uncle do not share one mother, with probability 1/2 the gene comes from the maternal grandmother, with probability $(1-m_{\mathrm{f}})^2$ neither of the maternal grandmother nor the uncle’s mother disperses, and the consanguinity is $p_{\mathrm{x}}'$, with probability 1/2 the gene comes from the maternal grandfather, and the consanguinity is $f'$, and with probability 1/2 the gene we pick comes from the juvenile’s father, in which case the consanguinity is, with probability 1/*n* the uncle is the juvenile’s father, and with probability 1/2 the gene comes from the paternal grandmother, and the consanguinity is $p_{\mathrm{I}}$, with probability 1/2 the gene comes from the paternal grandfather, and the consanguinity is $f'$, with probability (*n*-1)/*n* the uncle is not the juvenile's father, with probability $(1-m_{\mathrm{m}})^2$ neither of the two males disperses, with probability 1/*n* the uncle and the father have a same mother, with probability 1/2 the gene comes from the paternal grandmother, and the consanguinity is $p_{\mathrm{I}}$, with probability 1/2 the gene comes from the paternal grandfather, and the consanguinity is $f'$, with probability (*n*-1)/*n* the uncle and the father do not have a same mother, with probability 1/2 the gene comes from the paternal grandmother, with probability $(1-m_{\mathrm{f}})^2$ neither of the paternal grandmother nor the uncle's mother disperses, and the consanguinity is $p_{\mathrm{x}}'$, with probability 1/2 the gene comes from the paternal grandfather, and the consanguinity is $f'$. The consanguinity between the focal juvenile and the father of an uncle $p_{\mathrm{JPUF}}$ is

$$\begin{aligned}p_{\mathrm{JPUF}} = \frac{1}{2}(1-m_{\mathrm{f}})(1-m_{\mathrm{m}})\left(\frac{1}{n}\left(\frac{1}{2}f'+\frac{1}{2}p_{\mathrm{I}}\right)+\frac{n-1}{n}\left(\frac{1}{2}f'+\frac{1}{2}(1-m_{\mathrm{m}})^2p_{\mathrm{x}}'\right)\right)\\ +\frac{1}{2}\left(\frac{1}{n}\left(\frac{1}{2}f'+\frac{1}{2}p_{\mathrm{I}}\right)\right.\\ +\frac{n-1}{n}(1-m_{\mathrm{m}})^2\left(\frac{1}{n}\left(\frac{1}{2}f'+\frac{1}{2}p_{\mathrm{I}}\right)\right.\\ \left.\left.+\frac{n-1}{n}\left(\frac{1}{2}f'+\frac{1}{2}(1-m_{\mathrm{m}})^2p_{\mathrm{x}}'\right)\right)\right)\end{aligned} \quad (S95)$$

That is: with probability 1/2 the gene we pick comes from the juvenile's mother, in which case the consanguinity is, with probability $(1-m_{\mathrm{f}})(1-m_{\mathrm{m}})$ neither the mother nor the uncle disperses, and with probability 1/*n* the mother and the uncle share one father, and with probability 1/2 the gene comes from the maternal grandmother, and the consanguinity is $f'$, and with probability 1/2 the gene comes from the maternal grandfather, and the consanguinity is $p_{\mathrm{I}}$, and with probability (*n*-1)/*n* the mother and the uncle do not share one father, with probability 1/2 the gene comes from the maternal grandmother, and the consanguinity is $f'$, with probability 1/2 the gene comes from the maternal grandfather, with probability $(1-m_{\mathrm{m}})^2$ neither the uncle's father of nor the paternal grandfather disperses, and the consanguinity is $p_{\mathrm{x}}'$, with probability 1/2 the gene we pick comes from the juvenile's father, in which case the consanguinity is, with probability 1/*n* the uncle is the juvenile's father, and the consanguinity is that between the juvenile's father and its paternal grandfather which is $\frac{1}{2}f' + \frac{1}{2}p_{\mathrm{I}}$, and with probability (*n*-1)/*n* the uncle is not the juvenile's father, with probability $(1-m_{\mathrm{m}})^2$ neither of the two males disperses, and with probability 1/*n* the uncle and the father have a same father, with probability 1/2 the gene comes from the paternal grandmother, and the consanguinity is *f'*, with probability 1/2 the gene comes from the paternal grandfather, and the consanguinity is $p_{\mathrm{I}}$, and with probability (*n*-1)/*n* the uncle and the father do not have a same father, with probability 1/2 the gene comes from the paternal grandmother, and the consanguinity is *f'*, with probability 1/2 the gene comes from the paternal grandfather, with probability $(1-m_{\mathrm{m}})^2$ neither the grandfather nor the uncle's father disperses, and the consanguinity is $p_{\mathrm{x}}'$. Hence the consanguinity between the focal juvenile and its paternal grandparents $p_{\mathrm{JPGP}}$ is

$$p_{\mathrm{JPGP}} = \frac{1}{2}p_{\mathrm{JPGM}} + \frac{1}{2}p_{\mathrm{JPGF}} \tag{S96}$$

Similarly, the consanguinity between the focal juvenile and the parent of an uncle $p_{\mathrm{JPUP}}$ is

$$p_{\mathrm{JPUP}} = \frac{1}{2}p_{\mathrm{JPUM}} + \frac{1}{2}p_{\mathrm{JPUF}} \tag{S97}$$

*1.73 | Convergence stable strategy*

Solving expression (S86), we can get all the consanguinities:

$$\begin{aligned} p_{\mathrm{JMGM}} = (&-2\Delta m(M - 2\bar{m} + 1)(1 - \bar{m}) \\ &+ (m_{\mathrm{f}}(10 + H_{\mathrm{f}} - 2m_{\mathrm{f}}) - 8 + 6m_{\mathrm{m}} - m_{\mathrm{f}}(6 + H_{\mathrm{f}} - m_{\mathrm{f}})m_{\mathrm{m}} \\ &+ (2 - 3m_{\mathrm{f}}){m_{\mathrm{m}}}^2 - (1 - m_{\mathrm{f}}){m_{\mathrm{m}}}^3)n - 4\bar{m}(2 - \bar{m})n^2)/(8n(2\bar{m} - 1 \\ &- 4\bar{m}^2 + 3M - 4\bar{m}(2 - \bar{m})n)) \end{aligned} \tag{S98}$$

$$p_{\mathrm{JMGF}} = (2\Delta m(M - 2\bar{m} + 1)(1 - \bar{m}) + \big(m_{\mathrm{f}}^2(2 - 3m_{\mathrm{m}}) - 8 - m_{\mathrm{f}}^3(1 - m_{\mathrm{m}}) + m_{\mathrm{m}}(10 + H_{\mathrm{m}} - 2m_{\mathrm{m}}) - m_{\mathrm{f}}(m_{\mathrm{m}}(6 + H_{\mathrm{m}} - m_{\mathrm{m}}) - 6)\big)n - 4\bar{m}(2 - \bar{m})n^2)/(8n(2\bar{m} - 1 - 4\bar{m}^2 + 3M - 4\bar{m}(2 - \bar{m})n)) \quad \text{(S99)}$$

$$p_{\mathrm{JMGP}} = 1/8 - (7(M - 2\bar{m} + 1))/(8(2\bar{m} - 1 - 4\bar{m}^2 + 3M - 4\bar{m}(2 - \bar{m})n)) \quad \text{(S100)}$$

$$p_{\mathrm{JMAM}} = -(((-2\Delta m(H_{\mathrm{f}} + 1)(1 - \bar{m}) - \Delta m(-10 + 2m_{\mathrm{f}}^3 + m_{\mathrm{f}}(H_{\mathrm{m}} - 6m_{\mathrm{m}} + 16) - 3m_{\mathrm{f}}^2(3 - m_{\mathrm{m}}) - H_{\mathrm{m}} + 4m_{\mathrm{m}})n + (8 + m_{\mathrm{f}}^4 - m_{\mathrm{f}}^3(5 - m_{\mathrm{m}}) + (H_{\mathrm{m}} - 3m_{\mathrm{m}} + 4)m_{\mathrm{m}} + m_{\mathrm{f}}(3 - m_{\mathrm{m}})(H_{\mathrm{m}} - 4) - m_{\mathrm{f}}^2(m_{\mathrm{m}} - 11 + m_{\mathrm{m}}^2))\, n^2))\ /\ ((8n^2\,(2\bar{m} - 1 - 4\bar{m}^2 + 3M - 4\bar{m}(2 - \bar{m})n))\,)) \quad \text{(S101)}$$

$$p_{\mathrm{JMAF}} = ((-2\Delta m(H_{\mathrm{f}} + 1)(1 - \bar{m}) - \Delta m(H_{\mathrm{f}}(2m_{\mathrm{f}} - 5) - 2 + 4m_{\mathrm{m}} + m_{\mathrm{f}}(3m_{\mathrm{f}} - 8)m_{\mathrm{m}} - (1 - m_{\mathrm{f}})m_{\mathrm{m}}^2)n + (m_{\mathrm{f}}^4 - 8 - m_{\mathrm{f}}^3(5 - m_{\mathrm{m}}) + m_{\mathrm{m}}(4 + H_{\mathrm{m}} - m_{\mathrm{m}}) - m_{\mathrm{f}}((H_{\mathrm{m}} - 3m_{\mathrm{m}} + 6)m_{\mathrm{m}} - 4) - m_{\mathrm{f}}^2(m_{\mathrm{m}} - 5 + m_{\mathrm{m}}^2))n^2))\ /\ ((8n^2(2\bar{m} - 1 - 4\bar{m}^2 + 3M - 4\bar{m}(2 - \bar{m})n))) \quad \text{(S102)}$$

$$p_{\mathrm{JMAP}} = \frac{m_{\mathrm{m}}(4 + m_{\mathrm{m}}(n - 1)) - 3m_{\mathrm{f}}^2(n - 1) - 8n - 2m_{\mathrm{f}}(2 + m_{\mathrm{m}} - (4 - m_{\mathrm{m}})n)}{8n(2\bar{m} - 1 - 4\bar{m}^2 + 3M - 4\bar{m}(2 - \bar{m})n)} \quad \text{(S103)}$$

$$p_{\mathrm{JPGM}} = (-2\Delta m(M - 2\bar{m} + 1)(1 - \bar{m}) + (-8 + m_{\mathrm{f}}(10 + H_{\mathrm{f}} - 2m_{\mathrm{f}}) + 6m_{\mathrm{m}} - M(6 + H_{\mathrm{f}} - m_{\mathrm{f}}) + (2 - 3m_{\mathrm{f}})m_{\mathrm{m}}^2 - (1 - m_{\mathrm{f}})m_{\mathrm{m}}^3)n - 4\bar{m}(2 - \bar{m})n^2)/(8n(2\bar{m} - 1 - 4\bar{m}^2 + 3M - 4\bar{m}(2 - \bar{m})n)) \quad \text{(S104)}$$

$$p_{\mathrm{JPGF}} = (2\Delta m(M - 2\bar{m} + 1)(1 - \bar{m}) + (-8 + m_{\mathrm{f}}^2\,(2 - 3m_{\mathrm{m}}) - m_{\mathrm{f}}^3\,(1 - m_{\mathrm{m}}) + m_{\mathrm{m}}(10 + H_{\mathrm{m}} - 2m_{\mathrm{m}}) - m_{\mathrm{f}}(-6 + m_{\mathrm{m}}(6 + H_{\mathrm{m}} - m_{\mathrm{m}})))n - 4\bar{m}(2 - \bar{m})n^2)/(8n(2\bar{m} - 1 - 4\bar{m}^2 + 3M - 4\bar{m}(2 - \bar{m})n)) \quad \text{(S105)}$$

$$p_{\mathrm{JPGP}} = 1/8 - (7(M - 2\bar{m} + 1))/(8(2\bar{m} - 1 - 4\bar{m}^2 + 3M - 4\bar{m}(2 - \bar{m})n)) \quad \text{(S106)}$$

$$p_{\mathrm{JPUM}} = ((2\Delta m(H_\mathrm{m}+1)(1-\bar{m}) + \Delta m(-2 - m_\mathrm{f}{}^2(1-m_\mathrm{m}) + H_\mathrm{m}(2m_\mathrm{m}-5) + m_\mathrm{f}(3H_\mathrm{m} - 2m_\mathrm{m} + 4))n + (-8 + m_\mathrm{f}{}^3(1-m_\mathrm{m}) - m_\mathrm{f}{}^2(3 + H_\mathrm{m} - 3m_\mathrm{m}) + m_\mathrm{f}(4 + (H_\mathrm{m} - m_\mathrm{m})(2 + m_\mathrm{m})) + m_\mathrm{m}(4 + m_\mathrm{m}(5 + H_\mathrm{m} - 3m_\mathrm{m})))n^2)) \,/\, ((8n^2\,(2\bar{m} - 1 - 4\bar{m}^2 + 3M - 4\bar{m}(2-\bar{m})n))) \tag{S107}$$

$$p_{\mathrm{JPUF}} = ((-2\Delta m(H_\mathrm{m}+1)(1-\bar{m}) - \Delta m(-10 + 6m_\mathrm{f} - m_\mathrm{f}{}^2 + (H_\mathrm{m} - 6m_\mathrm{m} + 16)m_\mathrm{m} - 3(3 - m_\mathrm{f})m_\mathrm{m}{}^2 + 2m_\mathrm{m}{}^3)n + (-8 - m_\mathrm{f}{}^3\,(1-m_\mathrm{m}) + m_\mathrm{f}{}^2(5 + H_\mathrm{m} - 3m_\mathrm{m}) - m_\mathrm{m}(-12 + m_\mathrm{m}(11 + H_\mathrm{m} - 3m_\mathrm{m})) + m_\mathrm{f}(-4 + m_\mathrm{m}(2 + m_\mathrm{m} - m_\mathrm{m}{}^2)))n^2)) \,/\, ((8n^2\,(2\bar{m} - 1 - 4\bar{m}^2 + 3M - 4\bar{m}(2-\bar{m})n))) \tag{S108}$$

$$p_{\mathrm{JPUP}} = \frac{m_\mathrm{f}{}^2(n-1) - 8n + m_\mathrm{m}(-4 - 3m_\mathrm{m}(n-1) + 8n) - 2m_\mathrm{f}(m_\mathrm{m} - 2 + m_\mathrm{m}n)}{8n(-1 + 2\bar{m} - 4\bar{m}^2 + 3M - 4\bar{m}(2-\bar{m})n)} \tag{S109}$$

where $\Delta m = m_\mathrm{f} - m_\mathrm{m}$, $\bar{m} = (m_\mathrm{f} + m_\mathrm{m})/2$, $M = m_\mathrm{f}m_\mathrm{m}$, $\Delta b = b_\mathrm{f} - b_\mathrm{m}$, $\bar{b} = (b_\mathrm{f} + b_\mathrm{m})/2$, $H_\mathrm{f} = (m_\mathrm{f} - 2)m_\mathrm{f}$, $H_\mathrm{m} = (m_\mathrm{m} - 2)m_\mathrm{m}$, and by substituting these values, we obtain $z_{\mathrm{PO}}{}^*$, $z_{\mathrm{PD}}{}^*$, $z_{\mathrm{PS}}{}^*$, $z_{\mathrm{MO}}{}^*$, $z_{\mathrm{MD}}{}^*$, $z_{\mathrm{MS}}{}^*$, $z_{\mathrm{FO}}{}^*$, $z_{\mathrm{FD}}{}^*$ and $z_{\mathrm{FS}}{}^*$ for the optimal values of left-handedness when considering within-group combat

$$z_{\mathrm{PO}}{}^* = (((n-1)(\Delta m\big(b_\mathrm{f}(-4 + 3m_\mathrm{f} + m_\mathrm{m}) - b_\mathrm{m}(m_\mathrm{f} - 4 + 3m_\mathrm{m})\big) - 8\bar{b}\bar{m}(2-\bar{m})n))) \,/\, ((-2\Delta m(b_\mathrm{f}(3m_\mathrm{f} - 4 + m_\mathrm{m}) - b_\mathrm{m}(m_\mathrm{f} - 4 + 3m_\mathrm{m})) - 4(8 - 4(2 + b_\mathrm{m})m_\mathrm{f} + (1 - \Delta b)m_\mathrm{f}{}^2 + 2M(3 + 2\bar{b}) + m_\mathrm{m}(-8 - b_\mathrm{f}(4 - m_\mathrm{m}) + m_\mathrm{m} - b_\mathrm{m}m_\mathrm{m}))n - 16\bar{m}(\bar{b} + 1)(2-\bar{m})n^2)) \tag{S110}$$

$$z_{\mathrm{PD}}{}^* = ((b_\mathrm{f}(n-1)(-2m_\mathrm{f}(2 + m_\mathrm{m}) + (H_\mathrm{m} - 2m_\mathrm{m})(n-1) - 2m_\mathrm{f}(2 - m_\mathrm{m})n + m_\mathrm{f}{}^2(3 + n)))) \,/\, ((-2(8 + H_\mathrm{f} - 6m_\mathrm{f} - 8m_\mathrm{m} + 6m_\mathrm{f}m_\mathrm{m} + m_\mathrm{m}{}^2)n - 8\bar{m}(2-\bar{m})n^2 + 2b_\mathrm{f}(n-1)(-2m_\mathrm{f}(2 + m_\mathrm{m}) + (H_\mathrm{m} - 2m_\mathrm{m})(n-1) - 2m_\mathrm{f}(2 - m_\mathrm{m})n + m_\mathrm{f}{}^2(3 + n)))) \tag{S111}$$

$$z_{\mathrm{PS}}{}^{*} = ((b_{\mathrm{m}}(n-1)(m_{\mathrm{f}}{}^{2}(n-1) - 2m_{\mathrm{f}}(2-m_{\mathrm{m}})(n-1) + m_{\mathrm{m}}(-4(1+n) + m_{\mathrm{m}}(3 + n))))) \,/\, ((2b_{\mathrm{m}}\Delta m(m_{\mathrm{f}} - 4 + 3m_{\mathrm{m}}) - 2\big(8 + (1+2b_{\mathrm{m}})m_{\mathrm{f}}{}^{2} + m_{\mathrm{f}}(-8 - 4b_{\mathrm{m}}(2-m_{\mathrm{m}}) + 6m_{\mathrm{m}}) + m_{\mathrm{m}}(m_{\mathrm{m}} - 8 - 2b_{\mathrm{m}}m_{\mathrm{m}})\big)n - 8\bar{m}(1+b_{\mathrm{m}})(2-\bar{m})n^{2})) \tag{S112}$$

$$z_{\mathrm{MO}}{}^{*} = (((n-1)(2\Delta m(b_{\mathrm{f}}(H_{\mathrm{f}}+1) + b_{\mathrm{m}}(H_{\mathrm{m}}+1))(1-\bar{m}) + \Delta m(2b_{\mathrm{m}} - 2b_{\mathrm{f}}(3 - m_{\mathrm{m}}) + b_{\mathrm{m}}m_{\mathrm{m}}(2 - m_{\mathrm{f}}(2-m_{\mathrm{m}}) + H_{\mathrm{m}} - 2m_{\mathrm{m}}) + b_{\mathrm{f}}m_{\mathrm{f}}(8 - 2m_{\mathrm{m}} - 2m_{\mathrm{f}}(2-\bar{m})))n - 8\bar{b}\bar{m}(2-\bar{m})n^{2}))) \,/\, ((2(2n(-2\Delta m(1 - 2\bar{m} + M)(1-\bar{m}) + (-8 + m_{\mathrm{f}}(10 + H_{\mathrm{f}} - 2m_{\mathrm{f}}) + 6m_{\mathrm{m}} - m_{\mathrm{f}}(6 + H_{\mathrm{f}} - m_{\mathrm{f}})m_{\mathrm{m}} + (2 - 3m_{\mathrm{f}})m_{\mathrm{m}}{}^{2} - (1-m_{\mathrm{f}})m_{\mathrm{m}}{}^{3})n - 4\bar{m}(2-\bar{m})n^{2}) + b_{\mathrm{m}}(n - 1)(2\Delta m(H_{\mathrm{m}}+1)(1-\bar{m}) + \Delta m(2 + m_{\mathrm{m}}(2 - m_{\mathrm{f}}(2-m_{\mathrm{m}}) + H_{\mathrm{m}} - 2m_{\mathrm{m}}))n - 4\bar{m}(2-\bar{m})n^{2}) + b_{\mathrm{f}}(n-1)(2\Delta m(H_{\mathrm{f}}+1)(1-\bar{m}) + \Delta m(-2(3-m_{\mathrm{m}}) + m_{\mathrm{f}}(8 - 2m_{\mathrm{m}} - 2m_{\mathrm{f}}(2-\bar{m})))n - 4\bar{m}(2-\bar{m})n^{2})))) \tag{S113}$$

$$z_{\mathrm{MD}}{}^{*} = ((b_{\mathrm{f}}(n-1)(-2\Delta m(H_{\mathrm{f}}+1) + \Delta m(-2(3-m_{\mathrm{m}}) + m_{\mathrm{f}}(8 - 2m_{\mathrm{m}} - 2m_{\mathrm{f}}(2-\bar{m})))n - 4\bar{m}(2-\bar{m})n^{2}))) \,/\, ((2(n(-2\Delta m(1 - 2\bar{m} + M)(1-\bar{m}) + (-8 + m_{\mathrm{f}}(10 + H_{\mathrm{f}} - 2m_{\mathrm{f}}) + 6m_{\mathrm{m}} - m_{\mathrm{f}}(6 + H_{\mathrm{f}} - m_{\mathrm{f}})m_{\mathrm{m}} + (2 - 3m_{\mathrm{f}})m_{\mathrm{m}}{}^{2} - (1-m_{\mathrm{f}})m_{\mathrm{m}}{}^{3})n - 4\bar{m}(2-\bar{m})n^{2}) + b_{\mathrm{f}}(n - 1)(2\Delta m(H_{\mathrm{f}}+1)(1-\bar{m}) + \Delta m(-2(3-m_{\mathrm{m}}) + m_{\mathrm{f}}(8 - 2m_{\mathrm{m}} - 2m_{\mathrm{f}}(2-\bar{m})))n - 4\bar{m}(2-\bar{m})n^{2})))) \tag{S114}$$

$$z_{\mathrm{MS}}{}^{*} = ((b_{\mathrm{m}}(n-1)(2\Delta m(H_{\mathrm{m}}+1)(1-\bar{m}) + \Delta m(2+m_{\mathrm{m}}(2-m_{\mathrm{f}}(2-m_{\mathrm{m}})+H_{\mathrm{m}} - 2m_{\mathrm{m}}))n - 4\bar{m}(2-\bar{m})n^2))) \,/\, ((2n(-2\Delta m(1-2\bar{m}+M)(1-\bar{m}) + (-8+m_{\mathrm{f}}(10+H_{\mathrm{f}}-2m_{\mathrm{f}}) + 6m_{\mathrm{m}} - m_{\mathrm{f}}(6+H_{\mathrm{f}}-m_{\mathrm{f}})m_{\mathrm{m}} + (2-3m_{\mathrm{f}})m_{\mathrm{m}}{}^{2} - (1-m_{\mathrm{f}})m_{\mathrm{m}}{}^{3})n - 4\bar{m}(2-\bar{m})n^2) + 2b_{\mathrm{m}}(n-1)(2\Delta m(H_{\mathrm{m}}+1)(1-\bar{m}) + \Delta m(2+m_{\mathrm{m}}(2-m_{\mathrm{f}}(2-m_{\mathrm{m}})+H_{\mathrm{m}} - 2m_{\mathrm{m}}))n - 4\bar{m}(2-\bar{m})n^2))) \quad \text{(S115)}$$

$$z_{\mathrm{FO}}{}^{*} = -((((n-1)(-2\Delta m(b_{\mathrm{f}}(H_{\mathrm{f}}+1) + b_{\mathrm{m}}(H_{\mathrm{m}}+1))(1-\bar{m}) - \Delta m(b_{\mathrm{m}}(-6 + m_{\mathrm{m}}(8+H_{\mathrm{m}}-2m_{\mathrm{m}}) + m_{\mathrm{f}}(2+H_{\mathrm{m}})) + b_{\mathrm{f}}(2+m_{\mathrm{f}}(2-2m_{\mathrm{m}} - 2m_{\mathrm{f}}(2-\bar{m})))))n - 8\bar{b}\bar{m}(2-\bar{m})n^2))) \,/\, ((4n(-2\Delta m(1-2\bar{m}+M)(1-\bar{m}) + (8+m_{\mathrm{f}}(H_{\mathrm{f}}-6) - 10m_{\mathrm{m}} + m_{\mathrm{f}}(6-H_{\mathrm{f}}+m_{\mathrm{f}})m_{\mathrm{m}} + (4-3m_{\mathrm{f}})m_{\mathrm{m}}{}^{2} - (1-m_{\mathrm{f}})m_{\mathrm{m}}{}^{3})n + 4\bar{m}(2-\bar{m})n^2) - 2b_{\mathrm{m}}(n-1)(-2\Delta m(H_{\mathrm{m}}+1)(1-\bar{m}) - \Delta m(-6+m_{\mathrm{m}}(8+H_{\mathrm{m}}-2m_{\mathrm{m}}) + m_{\mathrm{f}}(2+H_{\mathrm{m}}))n - 4\bar{m}(2-\bar{m})n^2) - 2b_{\mathrm{f}}(n-1)(-2\Delta m(H_{\mathrm{f}}+1)(1-\bar{m}) - \Delta m(2+m_{\mathrm{f}}(2-2m_{\mathrm{m}}-2m_{\mathrm{f}}(2-\bar{m})))n - 4\bar{m}(2-\bar{m})n^2)))) \quad \text{(S116)}$$

$$z_{\mathrm{FD}}{}^{*} = ((b_{\mathrm{f}}(\mathrm{n}-1)(-2\Delta m(H_{\mathrm{f}}+1)(1-\bar{m}) + \Delta m(2+m_{\mathrm{f}}(2-2m_{\mathrm{m}} - 2m_{\mathrm{f}}(2-\bar{m})))n + 4\bar{m}(2-\bar{m})n^2))) \,/\, ((2n(-2\Delta m(1-2\bar{m}+M)(1-\bar{m}) + (8+m_{\mathrm{f}}(H_{\mathrm{f}}-6) - 10m_{\mathrm{m}} + m_{\mathrm{f}}(6-H_{\mathrm{f}}+m_{\mathrm{f}})m_{\mathrm{m}} + (4-3m_{\mathrm{f}})m_{\mathrm{m}}{}^{2} - (1-m_{\mathrm{f}})m_{\mathrm{m}}{}^{3})n + 4\bar{m}(2-\bar{m})n^2) + 4b_{\mathrm{f}}\Delta m(\mathrm{n}-1)(H_{\mathrm{f}}+1)(1-\bar{m}) - \Delta m(2+m_{\mathrm{f}}(2-2m_{\mathrm{m}} - 2m_{\mathrm{f}}(2-\bar{m})))n - 4\bar{m}(2-\bar{m})n^2))) \quad \text{(S117)}$$

$$z_{\mathrm{FS}}{}^{*} = -(((b_{\mathrm{m}}(n-1)(-m_{\mathrm{f}}{}^{2}(H_{\mathrm{m}}+1)-n)(n-1)+2m_{\mathrm{f}}(n-1)(H_{\mathrm{m}}+1)-(2-m_{\mathrm{m}})n)+m_{\mathrm{m}}((2-m_{\mathrm{m}})(H_{\mathrm{m}}+1)+(-6+m_{\mathrm{m}}(8+H_{\mathrm{m}}-2m_{\mathrm{m}}))n-(4-m_{\mathrm{m}})n^{2})))) \\ /\ ((2n(-2\Delta m(1-m_{\mathrm{f}})(1-m_{\mathrm{m}})(1-\bar{m}) + (8+m_{\mathrm{f}}(-6+H_{\mathrm{f}})-10m_{\mathrm{m}}+m_{\mathrm{f}}(6-H_{\mathrm{f}}+m_{\mathrm{f}})m_{\mathrm{m}} + (4-3m_{\mathrm{f}})m_{\mathrm{m}}{}^{2}-(1-m_{\mathrm{f}})m_{\mathrm{m}}{}^{3})n+4\bar{m}(2-\bar{m})n^{2})-2b_{\mathrm{m}}(\mathrm{n} - 1)(2\Delta m(H_{\mathrm{m}}+1)(1-\bar{m})-\Delta m(-6+m_{\mathrm{m}}(8+H_{\mathrm{m}}-2m_{\mathrm{m}}) + m_{\mathrm{f}}(2+H_{\mathrm{m}}))n-4\bar{m}(2-\bar{m})n^{2})))) \quad (S118)$$

where $\Delta m = m_{\mathrm{f}} - m_{\mathrm{m}}$, $\bar{m} = (m_{\mathrm{f}} + m_{\mathrm{m}})/2$, $M = m_{\mathrm{f}}m_{\mathrm{m}}$, $\Delta b = b_{\mathrm{f}} - b_{\mathrm{m}}$, $\bar{b} = (b_{\mathrm{f}} + b_{\mathrm{m}})/2$, $H_{\mathrm{f}} = (m_{\mathrm{f}} - 2)m_{\mathrm{f}}$, $H_{\mathrm{m}} = (m_{\mathrm{m}} - 2)m_{\mathrm{m}}$. We set the female dispersal rate $m_{\mathrm{f}}$ to be 0.5, the relative importance of combat relative to all types of competition for the female $b_{\mathrm{f}}$ and male $b_{\mathrm{m}}$ both to be 1, and number of the number of individuals each sex born in the same patch $n$ to be 5 for Figure 1*c* and Figure S1&S2.

Here we show what if there are differences between the parental genetic effects on daughters and those on sons in the context of within-group combats, hence left-handedness is marginally selfish. Under female-biased dispersal, the relatedness between the parent and the social partner through daughters' side would be lower than that through sons' side, hence genes carried by parents would favour a higher level of left-handedness for daughters than for sons; while under male-biased dispersal, the relatedness between social partners through daughters' side would be higher than that through sons' side, genes carried by parent would favour a lower expression level of left-handedness for daughters than for sons (Figure S2).

**2 | Between-group combat**

Here we make an illustration of the scenario where left-handedness is marginally altruistic, when between-group combat is the most frequent form of combat, as left-handed individuals are more likely to win the fights for their group, and this incurs a cost to themselves. The models here are based on the same life cycle, but with different fitness function. We investigate with the same process as that in “Within-group combat”, starting from “Kin selection”, through “Sex-biased dispersal”, “Parent-of-origin effect”, “Sex-specific effects” to “Parental genetic effects”. All the consanguinities are the same as those in the context of “Within-group combat”.

**2.1 | Kin selection**

We assume that an individual's payoff from between-group combat is proportional to the ratio of the competitive ability of the local group and the average competitive ability in the whole population. We assume that each group's competitive ability is proportional to the average disposition to the opposite handedness within their social arena. That is, with proportion *y* the members of the focal group are left-handed and have competitive ability 1-*z*, where *z* is the average proportion of left-handers in the whole population. And with proportion 1-*y* the members of the focal group are right handed and have competitive ability *z*. And the average competitive ability in the whole population is made up of the proportion *z* of left-handed individuals in an average group with competitive ability 1-*z* and the proportion 1-*z* of right-handed individuals in an average group with competitive ability *z*. Which gives

$$y\frac{(1-z)}{z(1-z)+(1-z)z}+(1-y)\frac{z}{z(1-z)+(1-z)z} \qquad \text{(S119)}$$

Which simplifies to

$$\frac{y}{2z}+\frac{1-y}{2(1-z)} \qquad \text{(S120)}$$

Accordingly, the fitness of a juvenile *w’* is

$$w' = \left(1 - b_{\mathrm{f}} + b_{\mathrm{f}}\left(\frac{y_{\mathrm{Mo}}}{2z} + \frac{1 - y_{\mathrm{Mo}}}{2(1-z)}\right)\right)(1 - c_{\mathrm{f}}x_{\mathrm{Mo}})\left(1 - b_{\mathrm{m}} + b_{\mathrm{m}}\left(\frac{y_{\mathrm{Fa}}}{2z} + \frac{1 - y_{\mathrm{Fa}}}{2(1-z)}\right)\right)(1 - c_{\mathrm{m}}x_{\mathrm{Fa}}) \quad \text{(S121)}$$

Similarly, the average fitness of a random juvenile $\overline{w'}$ can be described by evaluating expression (S121) at $x_{\mathrm{Mo}} = y_{\mathrm{Mo}} = z_{\mathrm{f}}$, $x_{\mathrm{Fa}} = y_{\mathrm{Fa}} = z_{\mathrm{m}}$, and the relative fitness of the focal juvenile $W'$ is $w'/\overline{w'}$

$$W' = \left(1 - b_{\mathrm{f}} + b_{\mathrm{f}}\left(\frac{y_{\mathrm{Mo}}}{2z} + \frac{1 - y_{\mathrm{Mo}}}{2(1-z)}\right)\right)\left(\frac{1 - c_{\mathrm{f}}x_{\mathrm{Mo}}}{1 - c_{\mathrm{f}}z_{\mathrm{f}}}\right)\left(1 - b_{\mathrm{m}} + b_{\mathrm{m}}\left(\frac{y_{\mathrm{Fa}}}{2z} + \frac{1 - y_{\mathrm{Fa}}}{2(1-z)}\right)\right)\left(\frac{1 - c_{\mathrm{m}}x_{\mathrm{Fa}}}{1 - c_{\mathrm{m}}z_{\mathrm{m}}}\right) \quad \text{(S122)}$$

Similarly using expression (S122), we obtain the condition for an increase in left-handedness to be favoured when we consider between-group combat

$$\frac{(b_{\mathrm{f}} + b_{\mathrm{m}})(1 - 2z)r_{\mathrm{J}}}{2(1-z)z} - \frac{c_{\mathrm{f}}r_{\mathrm{O}}}{1 - c_{\mathrm{f}}z} - \frac{c_{\mathrm{m}}r_{\mathrm{O}}}{1 - c_{\mathrm{m}}z} > 0 \quad \text{(S123)}$$

Letting the LHS of expression (S7) be $f(z)$, then at evolutionary equilibrium, if there is an intermediate level of left-handedness $z'^{*}$, this satisfies $f(z'^{*}) = 0$, we get the optimal value of developing as left-handed for a random individual when we consider between-group combat

$$z'^{*} = \frac{1}{2}\frac{(b_{\mathrm{f}} + b_{\mathrm{m}})r_{\mathrm{JR}}}{r_{\mathrm{JR}}(b_{\mathrm{f}} + b_{\mathrm{m}}) - 2r_{\mathrm{O}}} \quad \text{(S124)}$$

Substituting all the parameters of relatedness to expression (S124), we can get the optimal value of left-handedness for the genes at locus G when left-handedness is altruistic, $z'^{*}$

$$\mathrm{z}'^{*} = \frac{1}{2}\frac{b_{\mathrm{f}} + b_{\mathrm{m}}}{2 + b_{\mathrm{f}} + b_{\mathrm{m}} + 2(1 - (1 - m)^2)(n - 1)} \quad \text{(S125)}$$

#### 2.2 | Sex-biased dispersal

Here we relax the assumption of no sex bias in dispersal i.e. $m_f \neq m_m$, hence $p_{JA} \neq p_{JU}$. In this section, the relative fitness function is the same as expression (S122). Using expressions (S122) to calculate the corresponding partial derivatives, we obtain the condition for an increase in left-handedness to be favoured when we consider between-group combat

$$-\frac{(b_f r_{JA} + b_m r_{JU})(1-2z)}{2(1-z)z} - \frac{c_f r_O}{1-c_f z} - \frac{c_m r_O}{1-c_m z} > 0 \qquad (S126)$$

Letting $f(z)$ be the LHS of expression (S126), than at evolutionary equilibrium, if there is an intermediate level of left-handedness, this satisfies $f(z'^*) = 0$, we obtain the optimum of left-handedness in the context of between-group combat. For example, letting $c_f = c_m = 1$, i.e. there is no sex difference in the cost of developing as left-handed, we have

$$z'^* = \frac{1}{2}\frac{b_f r_{JA} + b_m r_{JU}}{b_f r_{JA} + b_m r_{JU} + 2r_O} \qquad (S127)$$

This is the convergence stable strategy, i.e. the overall optima level of left-handedness for all the loci involved, as $f'(z) < 0$ is true for all the values of $z$. Here all the consanguinity are the same as the previous section under the situation of "within-group combat", substituting all the parameters of relatedness to expression (S21), we obtain the optimal value of left-handedness $z'^*$

$$z'^* = (2\Delta b \Delta m(1-\bar{m}) + b_f(4 + H_f - H_m)n + b_m(4 - H_f + H_m)n)/(4\Delta b \Delta m(1 - \bar{m}) + 2(8(1-\bar{m})^2 + b_f(4 + H_f - H_m + b_m(4 - H_f + H_m))n + 16(2-\bar{m})\bar{m}n^2) \qquad (S128)$$

where $\Delta m = m_f - m_m$, $\bar{m} = (m_f + m_m)/2$, $\Delta b = b_f - b_m$, $\bar{b} = (b_f + b_m)/2$, $H_f = (m_f - 2)m_f$, $H_m = (m_m - 2)m_m$.

### 2.3 | Parent-of-origin effects

Here we consider how the origin of genes mediates the role of kin selection in the optima of different set of genes, under the circumstances of between-group combat. In this section the conditions that favour the increase of left-handedness in the population and the relatedness are the same as previous section "§S1.5 Parental-of-origin effects" when considering within-group combat, while the relative fitness function change to expression (S122). Letting the LHS of the expression (S28) be $f(z)$, then at evolutionary equilibrium, if there is an intermediate level of left-handedness $z_{\mathrm{M}}'^{*}$ and $z_{\mathrm{M}}'^{*}$, which satisfies $f(z_M'^{*}) = 0$ and $f(z_P'^{*}) = 0$, respectively, we obtain the optima

$$z_{\mathrm{M}}'^{*} = \frac{1}{2}\frac{b_{\mathrm{f}} r_{\mathrm{JA|-M}} + b_{\mathrm{m}} r_{\mathrm{JU|-M}}}{2r_{\mathrm{O|-M}} + b_{\mathrm{f}} r_{\mathrm{JA|-M}} + b_{\mathrm{m}} r_{\mathrm{JU|-M}}} \tag{S129}$$

$$z_{\mathrm{P}}'^{*} = \frac{1}{2}\frac{b_{\mathrm{f}} r_{\mathrm{JA|-P}} + b_{\mathrm{m}} r_{\mathrm{JU|-P}}}{2r_{\mathrm{O|-P}} + b_{\mathrm{f}} r_{\mathrm{JA|-P}} + b_{\mathrm{m}} r_{\mathrm{JU|-P}}} \tag{S130}$$

$f'(z) < 0$ is true for all the values of $z$, thus $z_{\mathrm{M}}'^{*}$ and $z_{\mathrm{P}}'^{*}$ are the optimal values of left-handedness from the perspective of maternal- and paternal-origin genes, respectively. Substituting all the parameters of relatedness, we obtain optimal value of maternal-origin genes, $z_{\mathrm{M}}'^{*}$

$$\begin{aligned} z_{\mathrm{M}}'^{*} = {} & ((b_{\mathrm{m}}(-2\Delta m(H_{\mathrm{m}}+1)(1-\bar{m}) + 2\Delta m(1-\bar{m})(1-M-2\bar{m}+2H_{\mathrm{m}})n \\ & + (8-2\Delta m(1-\bar{m})(M-2\bar{m}+H_{\mathrm{m}}))n^2) + b_{\mathrm{f}}(H_{\mathrm{f}} \\ & + 1)(-2\Delta m(1-\bar{m}) + 2\Delta m(1-\bar{m})(5-2\bar{m}+2H_{\mathrm{f}}+M)n \\ & + (8+m_{\mathrm{f}}{}^4 - m_{\mathrm{f}}{}^3(5-m_{\mathrm{m}}) - (4-m_{\mathrm{m}})H_{\mathrm{m}} - m_{\mathrm{f}}(8+(H_{\mathrm{m}} \\ & - 3m_{\mathrm{m}}+4)m_{\mathrm{m}}) - m_{\mathrm{f}}{}^2\,(-10+3m_{\mathrm{m}}+H_{\mathrm{m}}))n^2))) \\ & /\ ((2(-2b_{\mathrm{m}}\Delta m(H_{\mathrm{m}}+1)(1-\bar{m}) - 2\Delta m(1-\bar{m})(b_{\mathrm{m}}+2(M \\ & - 2\bar{m}+1) + b_{\mathrm{m}}(M-m_{\mathrm{f}}) + b_{\mathrm{m}}(2H_{\mathrm{m}}-m_{\mathrm{m}}))n + (b_{\mathrm{m}}(8 \\ & - 2\Delta m(1-\bar{m})(M-2\bar{m}+H_{\mathrm{m}})) - 4(1-\bar{m})(-4-m_{\mathrm{f}}{}^2(1 \\ & - m_{\mathrm{m}}) + m_{\mathrm{m}} + m_{\mathrm{m}}{}^2 - m_{\mathrm{f}}(m_{\mathrm{m}}{}^2-3)))n^2 + 16(2-\bar{m})\bar{m}n^3 \\ & + b_{\mathrm{f}}(-2\Delta m(H_{\mathrm{f}}+1)(1-\bar{m}) + 2\Delta m(1-\bar{m})(5-2\bar{m}+2H_{\mathrm{f}} \\ & + M)n + (8+m_{\mathrm{f}}{}^4 - m_{\mathrm{f}}{}^3(5-m_{\mathrm{m}}) - (4-m_{\mathrm{m}})H_{\mathrm{m}} - m_{\mathrm{f}}(8 \\ & + (H_{\mathrm{m}}-3m_{\mathrm{m}}+4)m_{\mathrm{m}}) - m_{\mathrm{f}}{}^2(-10+H_{\mathrm{m}}+3m_{\mathrm{m}}))n^2)))) \end{aligned} \tag{S131}$$

With similar process, we obtain the optimal value left-handedness $z_{\mathrm{P}}'^{*}$:

$$
\begin{aligned}
z_{\mathrm{P}}'^{*} = {} & ((-2b_{\mathrm{m}}\Delta m(1-\bar{m})(H_{\mathrm{m}}+1) + 2b_{\mathrm{m}}\Delta m(1-\bar{m})(5+M-2\bar{m} \\
& + 2H_{\mathrm{m}})n - 8b_{\mathrm{f}}n^2 + b_{\mathrm{m}}(-8+(4-m_{\mathrm{f}})H_{\mathrm{f}} - H_{\mathrm{m}}(4+H_{\mathrm{m}} \\
& - m_{\mathrm{m}}) + M(4+2\bar{m}\Delta m + M - 4m_{\mathrm{f}} - \Delta m))n^2 - 2b_{\mathrm{f}}\Delta m(1 \\
& - \bar{m})(H_{\mathrm{f}}+1+(2\bar{m}-1-2H_{\mathrm{f}}-M)n + ((2\bar{m}-3)m_{\mathrm{f}} \\
& - m_{\mathrm{m}})n^2))) \,/\, ((2(-2b_{\mathrm{m}}\Delta m(1-\bar{m})(H_{\mathrm{m}}+1) + 2\Delta m(1 \\
& - \bar{m})(2(M-2\bar{m}+1) + b_{m}(5+M-2\bar{m}+2H_{\mathrm{m}}))n \qquad (S132) \\
& + (b_{\mathrm{m}}(-8+(4-m_{\mathrm{f}})H_{\mathrm{f}} - H_{\mathrm{m}}(4+H_{\mathrm{m}}-m_{\mathrm{m}}) + M(4 \\
& + 2\bar{m}\Delta m + M - 4m_{\mathrm{f}} - \Delta m)) - 4(1-\bar{m})(4-m_{\mathrm{f}}^2(1-m_{\mathrm{m}}) \\
& + H_{\mathrm{m}} - m_{\mathrm{m}} - m_{\mathrm{f}}(1+m_{\mathrm{m}}^2)))n^2 - 16(2-\bar{m})\bar{m}n^3 \\
& + b_{\mathrm{f}}(-8n^2 - 2\Delta m(1-\bar{m})(H_{\mathrm{f}}+1+(2\bar{m}-1-2H_{\mathrm{f}}-M)n \\
& + ((2\bar{m}-3)m_{\mathrm{f}} - m_{\mathrm{m}})n^2)))))
\end{aligned}
$$

The optimal value of left-handedness for the perspective of the whole genes of the individual $z'^{*}$ is

$$
\begin{aligned}
z'^{*} = {} & (2\Delta b\Delta m(1-\bar{m}) + (b_{\mathrm{f}}(4+H_{\mathrm{f}}-H_{\mathrm{m}}) + b_{\mathrm{m}}(4-H_{\mathrm{f}} \\
& + H_{\mathrm{m}}))n)/(4\Delta b\Delta m(1-\bar{m}) - 2(b_{\mathrm{m}}(H_{\mathrm{f}}-H_{\mathrm{m}}-4) - 8 \qquad (S133) \\
& - b_{\mathrm{f}}(4+H_{\mathrm{f}}-H_{\mathrm{m}}) - 8\bar{m}(2-\bar{m})(n-1))n)
\end{aligned}
$$

where $\Delta m = m_{\mathrm{f}} - m_{\mathrm{m}}$, $\bar{m} = (m_{\mathrm{f}} + m_{\mathrm{m}})/2$, $\Delta b = b_{\mathrm{f}} - b_{\mathrm{m}}$, $\bar{b} = (b_{\mathrm{f}} + b_{\mathrm{m}})/2$, $H_{\mathrm{f}} = (m_{\mathrm{f}} - 2)m_{\mathrm{f}}$, $H_{\mathrm{m}} = (m_{\mathrm{m}} - 2)m_{\mathrm{m}}$. We set the female dispersal rate $m_{\mathrm{f}}$ to be 0.5, the relative importance of combat relative to all types of competition for the female $b_{\mathrm{f}}$ and male $b_{\mathrm{m}}$ both to be 1, and the number of individuals each sex born in the same patch $n$ to be 5 for Figure 2. For the zoomed-in parts, the range of male dispersal rate $m_{\mathrm{m}}$ is from 0.499 to 0.501, the range of the equilibrium frequency of left-handedness is from 0.09995 to 0.10005.

### 2.4 | Sex-specific effects

Here we consider how sex effects add to the mediation of kin selection on handedness under the circumstances of between-group combat. In this section, the conditions that favour the increase of left-handedness, the relatedness are the same as the previous section "§S1.6 Sex-specific effects" when considering within-group combat, while the relative fitness function changes to expression (S122). For locus $G_1$ which only controls the handedness trait of females, using similar methods as previous sections, letting the LHS of expression (S50) be $f(z)$, $f'(z) < 0$ is true for all the values of z and all of the four coefficients of relatedness above, at evolutionary equilibrium, if there is an intermediate level of left-handedness $z_f'^*$, this satisfies $f(z_f'^*) = 0$, we obtain the optimal value of left-handedness $z_f'^*$ for all the loci that control handedness only when they are carried by females

$$z_f'^* = \frac{1}{2}\frac{b_f r_{JA}}{r_{OM} + b_f r_{JA}} \tag{S134}$$

Similarly, we obtain the optimal value of locus $G_2$ when left-handedness is altruistic, $z_m'^*$

$$z_m'^* = \frac{1}{2}\frac{b_m r_{JU}}{r_{OF} + b_m r_{JU}} \tag{S135}$$

Similarly, we can obtain the optimal value for the locus $G_1$ from the perspective of maternal-origin genes, $z_{fM}'^*$, and that from the perspective of paternal-origin genes, $z_{fP}'^*$, and the optimal value for the locus $G_2$ from the perspective of maternal-origin genes and paternal-origin genes respectively: $z_{mM}'^*$ and $z_{mP}'^*$

$$z_{fM}'^* = \frac{1}{2}\frac{b_f r_{JA|-M}}{r_{OM|-M} + b_f r_{JA|-M}} \tag{S136}$$

$$z_{fP}'^* = \frac{1}{2}\frac{b_f r_{JA|-P}}{r_{OM|-P} + b_f r_{JA|-P}} \tag{S137}$$

$$z_{mM}'^* = \frac{1}{2}\frac{b_m r_{JU|-M}}{r_{OF|-M} + b_m r_{JU|-M}} \tag{S138}$$

$$z_{mP}'^* = \frac{1}{2}\frac{b_m r_{JU|-P}}{r_{OF|-P} + b_m r_{JU|-P}} \tag{S139}$$

Substituting all the relatedness in expressions (S134)-(S139) we obtain the optimal values of left-handedness when considering between-group combat:

$$z_{\mathrm{f}}{}'^{*} = \frac{b_{\mathrm{f}}\big(H_{\mathrm{m}} - H_{\mathrm{f}} + 2\big(2 - \Delta m(1-\bar{m})\big)n\big)}{8n + 8\bar{m}(2-\bar{m})(n-1)n + 2b_{\mathrm{f}}\big(H_{\mathrm{m}} - H_{\mathrm{f}} + 2\big(2 - \Delta m(1-\bar{m})\big)n\big)} \tag{S140}$$

$$\begin{aligned} z_{\mathrm{fM}}{}'^{*} = {} & ((b_{\mathrm{f}}((8 + H_{\mathrm{f}}(4 + H_{\mathrm{f}} - m_{\mathrm{f}}) - H_{\mathrm{m}}(4 - m_{\mathrm{m}}) \\ & + M(H_{\mathrm{f}} - H_{\mathrm{m}} + 2\bar{m} + 2m_{\mathrm{m}} - M))n^2 - 2\Delta m(1-\bar{m})(H_{\mathrm{f}} + 1 \\ & + (2\bar{m} - 5 - 2H_{\mathrm{f}} - M)n)))) \,/\, ((2(2\Delta m(1-\bar{m})(M - 2\bar{m} + 1)n \\ & + b_{\mathrm{f}}(8 + H_{\mathrm{f}}(4 + H_{\mathrm{f}} - m_{\mathrm{f}}) - H_{\mathrm{m}}(4 - m_{\mathrm{m}}) + M(H_{\mathrm{f}} - H_{\mathrm{m}} + 2\bar{m} \\ & + 2m_{\mathrm{m}} - 4 - M))n^2 + 2n^2(-(1-\bar{m})(-4 + M\Delta m - 2\bar{m}\Delta m + 2\bar{m} \\ & + 2m_{\mathrm{f}}) + 4(2-\bar{m})\bar{m}n) - 2b_{\mathrm{f}}\Delta m(1-\bar{m})(H_{\mathrm{f}} + 1 + (2\bar{m} - 5 - 2H_{\mathrm{f}} \\ & - M)n)))) \end{aligned} \tag{S141}$$

$$\begin{aligned} z_{\mathrm{fP}}{}'^{*} = {} & -(((b_{\mathrm{f}}(-8n^2 - 2\Delta m(1-\bar{m})(H_{\mathrm{f}} + 1 + (2\bar{m} - 1 - 2H_{\mathrm{f}} - M)n + ((2\bar{m} \\ & - 3)m_{\mathrm{f}} - m_{\mathrm{m}})n^2)))) \,/\, ((2(2\Delta m(1-\bar{m})(M - 2\bar{m} + 1)(1 - m_{\mathrm{m}})n \\ & + 8b_{\mathrm{f}}n^2 - 2(1-\bar{m})(2\bar{m} + 2m_{\mathrm{m}} - 4 + 2\bar{m}\Delta m - M\Delta m)n^2 \\ & + 8(2-\bar{m})\bar{m}n^3 + 2b_{\mathrm{f}}\Delta m(1-\bar{m})(H_{\mathrm{f}} + 1 + (2\bar{m} - 1 - 2H_{\mathrm{f}} - M)n \\ & + ((2\bar{m} - 3)m_{\mathrm{f}} - m_{\mathrm{m}})n^2))))) \end{aligned} \tag{S142}$$

$$z_{\mathrm{m}}{}'^{*} = \frac{b_{\mathrm{m}}(H_{\mathrm{f}} - H_{\mathrm{m}} + 2(2 + \Delta m - \Delta m\bar{m})n)}{8n + 8\bar{m}(2-\bar{m})(n-1)n + 2b_{\mathrm{m}}(H_{\mathrm{f}} - H_{\mathrm{m}} + 2(2 + \Delta m - \Delta m\bar{m})n)} \tag{S143}$$

$$\begin{aligned} z_{\mathrm{mM}}{}'^{*} = {} & ((-b_{\mathrm{m}}(-2(1-\bar{m})(H_{\mathrm{m}} + 1)\Delta m - 2\Delta m(1-\bar{m})(1 + M - 2\bar{m} + 2H_{\mathrm{m}})n \\ & + (-8 + 2\Delta m(1-\bar{m})(M - 2\bar{m} + H_{\mathrm{m}}))n^2))) \\ & /\, ((2(n(-2\Delta m(1-\bar{m})(M - 2\bar{m} + 1) \\ & + 2(1-\bar{m})(M\Delta m - 4 - 2\bar{m}\Delta m + 2\bar{m} + 2m_{\mathrm{f}})n - 8(2-\bar{m})\bar{m}n^2) \\ & + b_{\mathrm{m}}(2(1-\bar{m})(H_{\mathrm{m}} + 1)\Delta m - 2\Delta m(1-\bar{m})(1 + M - 2\bar{m} + 2H_{\mathrm{m}})n \\ & + (-8 + 2\Delta m(1-\bar{m})(M - 2\bar{m} + H_{\mathrm{m}}))n^2)))) \end{aligned} \tag{S144}$$

$$
\begin{aligned}
z_{\mathrm{mP}}{}'^{*} = -(((&b_{\mathrm{m}}\Delta m(-2(1-\bar{m})(H_{\mathrm{m}}+1) + 2\Delta m(1-\bar{m})(5+M-2\bar{m}+2H_{\mathrm{m}})n \\
&+ (-8+(4-m_{\mathrm{f}})H_{\mathrm{f}} - H_{\mathrm{m}}(4+H_{\mathrm{m}}-m_{\mathrm{m}}) + M(4+2\bar{m}\Delta m + M \\
&- 4m_{\mathrm{f}} - \Delta m))n^2))) \;/\; ((2(n(-2\Delta m(1-\bar{m})(M-2\bar{m}+1) \\
&- 2(1-\bar{m})(2\bar{m}-4+2\bar{m}\Delta m+2m_{\mathrm{m}}-M\Delta m)n + 8(2-\bar{m})\bar{m}n^2) \\
&+ b_{\mathrm{m}}(2(1-\bar{m})(H_{\mathrm{m}}+1)\Delta m - 2\Delta m(1-\bar{m})(5+M-2\bar{m}+2H_{\mathrm{m}})n \\
&+ (8-H_{\mathrm{f}}(4-m_{\mathrm{f}}) + m_{\mathrm{m}}(-8-(H_{\mathrm{f}}-3m_{\mathrm{f}}+4)m_{\mathrm{f}}+10m_{\mathrm{m}}-M \\
&- M\Delta m - 5{m_{\mathrm{m}}}^2 + {m_{\mathrm{m}}}^3))n^2)))))
\end{aligned}
\tag{S145}
$$

where $\Delta m = m_{\mathrm{f}} - m_{\mathrm{m}}$, $\bar{m} = (m_{\mathrm{f}} + m_{\mathrm{m}})/2$, $M = m_{\mathrm{f}}m_{\mathrm{m}}$, $\Delta b = b_{\mathrm{f}} - b_{\mathrm{m}}$, $\bar{b} = (b_{\mathrm{f}} + b_{\mathrm{m}})/2$, $H_{\mathrm{f}} = (m_{\mathrm{f}} - 2)m_{\mathrm{f}}$, $H_{\mathrm{m}} = (m_{\mathrm{m}} - 2)m_{\mathrm{m}}$. We set the female dispersal rate $m_{\mathrm{f}}$ to be 0.5, the relative importance of combat relative to all types of competition for the female $b_{\mathrm{f}}$ and male $b_{\mathrm{m}}$ both to be 1, and number of the number of individuals each sex born in the same patch $n$ to be 5 for Figure 1$b$.

**2.5 | Parental genetic effects**

Here we consider how parental effects mediate handedness considering handedness under the circumstances of between-group combat. In this section the coefficients of relatedness and all the nine situations are the same as previous section "§S1.7 Parental genetic effects" when considering within-group combat, but the relative fitness function changes to expression (S122). Using similar methods as previous sections, letting the LHS of expression (S66) be $f(z)$, $f'(z) < 0$ is true for all the values of $z$ and all of the four relatedness, then at evolutionary equilibrium, if there is an intermediate level of left-handedness $z_{\mathrm{PO}}{}'^{*}$, this satisfies $f(z_{PO}{}'^{*}) = 0$, we obtain the optimum of left-handedness from the perspective of parent's genes

$$
z_{\mathrm{PO}}{}'^{*} = \frac{1}{2}\frac{b_{\mathrm{f}}r_{\mathrm{JMAP}} + b_{\mathrm{m}}r_{\mathrm{JPUP}}}{b_{\mathrm{f}}r_{\mathrm{JMAP}} + r_{\mathrm{JMGP}} + r_{\mathrm{JPGP}} + b_{\mathrm{m}}r_{\mathrm{JPUP}}} \tag{S146}
$$

Similarly, we can obtain the optimal value of left-handedness from the perspective of parent's genes to its daughter

$$z_{\mathrm{PD}}{'}^{*} = \frac{1}{2}\frac{b_{\mathrm{f}} r_{\mathrm{JMAP}}}{b_{\mathrm{f}} r_{\mathrm{JMAP}} + r_{\mathrm{JMGP}}} \tag{S147}$$

the optimal value of left-handedness from the perspective of parent's genes to its son

$$z_{\mathrm{PS}}{'}^{*} = \frac{1}{2}\frac{b_{\mathrm{m}} r_{\mathrm{JPUP}}}{r_{\mathrm{JPGP}} + b_{\mathrm{m}} r_{\mathrm{JPUP}}} \tag{S148}$$

the optimal value of left-handedness from the perspective of mother's genes to her offspring

$$z_{\mathrm{MO}}{'}^{*} = \frac{1}{2}\frac{b_{\mathrm{f}} r_{\mathrm{JMAM}} + b_{\mathrm{m}} r_{\mathrm{JPUM}}}{b_{\mathrm{f}} r_{\mathrm{JMAM}} + r_{\mathrm{JMGM}} + r_{\mathrm{JPGM}} + b_{\mathrm{m}} r_{\mathrm{JPUM}}} \tag{S149}$$

the optimal value of left-handedness from the perspective of mother's genes to her daughters

$$z_{\mathrm{MD}}{'}^{*} = \frac{1}{2}\frac{b_{\mathrm{f}} r_{\mathrm{JMAM}}}{b_{\mathrm{f}} r_{\mathrm{JMAM}} + r_{\mathrm{JMGM}}} \tag{S150}$$

the optimal value of left-handedness from the perspective of mother's genes to her sons

$$z_{\mathrm{MS}}{'}^{*} = \frac{1}{2}\frac{b_{\mathrm{m}} r_{\mathrm{JPUM}}}{r_{\mathrm{JPGM}} + b_{\mathrm{m}} r_{\mathrm{JPUM}}} \tag{S151}$$

the optimal value of left-handedness from the perspective of father's genes to his offspring

$$z_{\mathrm{FO}}{'}^{*} = \frac{1}{2}\frac{b_{\mathrm{f}} r_{\mathrm{JMAF}} + b_{\mathrm{m}} r_{\mathrm{JPUF}}}{b_{\mathrm{f}} r_{\mathrm{JMAF}} + r_{\mathrm{JMGF}} + r_{\mathrm{JPGF}} + b_{\mathrm{m}} r_{\mathrm{JPUF}}} \tag{S152}$$

the optimal value of left-handedness from the perspective of father's genes to his daughters

$$z_{\mathrm{FD}}{'}^{*} = \frac{1}{2}\frac{b_{\mathrm{f}} r_{\mathrm{JMAF}}}{b_{\mathrm{f}} r_{\mathrm{JMAF}} + r_{\mathrm{JMGF}}} \tag{S153}$$

and the optimal value of left-handedness from the perspective of father's genes to his sons

$$z_{\mathrm{FS}}{'}^{*} = \frac{1}{2}\frac{b_{\mathrm{m}} r_{\mathrm{JPUF}}}{r_{\mathrm{JPGF}} + b_{\mathrm{m}} r_{\mathrm{JPUF}}} \tag{S154}$$

Substituting all of the relatedness, we obtain the optimal values of left-handedness when considering between-group combat

$$z_{\mathrm{PO}}{}'^{*} = ((-(2\Delta m(-2\Delta b + b_{\mathrm{f}}m_{\mathrm{f}} - b_{\mathrm{m}}m_{\mathrm{m}} + \bar{m}\Delta b)) + (2b_{\mathrm{f}}(4 + \bar{m}\Delta m + H_{\mathrm{f}} + M - 2m_{\mathrm{f}}) + 2b_{\mathrm{m}}(4 - 4m_{\mathrm{m}} - \bar{m}(m_{\mathrm{f}} - 3m_{\mathrm{m}})))n)) \,/\, ((-2\Delta m(b_{\mathrm{f}}(-4 + 3m_{\mathrm{f}} + m_{\mathrm{m}}) - b_{\mathrm{m}}(-4 + m_{\mathrm{f}} + 3m_{\mathrm{m}})) + 2(2b_{\mathrm{f}}(4 + \bar{m}\Delta m + H_{\mathrm{f}} + M - 2m_{\mathrm{f}}) + 2(8 + H_{\mathrm{f}} - 12\bar{m} + 6M + H_{\mathrm{m}}) + b_{\mathrm{m}}(8 - 8m_{\mathrm{m}} - 2\bar{m}(m_{\mathrm{f}} - 3m_{\mathrm{m}})))n + 16\bar{m}(2 - \bar{m})n^2)) \quad \text{(S155)}$$

$$z_{\mathrm{PD}}{}'^{*} = ((b_{\mathrm{f}}(3m_{\mathrm{f}}{}^2(n-1) + 8n + 2m_{\mathrm{f}}(2 + m_{\mathrm{m}} + (m_{\mathrm{m}} - 4)n) + m_{\mathrm{m}}(-4 + m_{\mathrm{m}} - m_{\mathrm{m}}n)))) \,/\, ((8n(2 - 4\bar{m} + \bar{m}^2 + M + \bar{m}(2 - \bar{m})n) + 2b_{\mathrm{f}}(3m_{\mathrm{f}}{}^2(\mathrm{n} - 1) + 8n + 2m_{\mathrm{f}}(2 + m_{\mathrm{m}} + (m_{\mathrm{m}} - 4)n) + m_{\mathrm{m}}(m_{\mathrm{m}} - 4 - m_{\mathrm{m}}n)))) \quad \text{(S156)}$$

$$z_{\mathrm{PS}}{}'^{*} = (b_{\mathrm{m}}(m_{\mathrm{f}}{}^2(n-1) - 8n + m_{\mathrm{m}}(-4 - 3m_{\mathrm{m}}(n-1) + 8n) - 2M + 4m_{\mathrm{f}} - 2Mn)) \,/\, ((-2b_{\mathrm{m}}\Delta m(-4 + m_{\mathrm{f}} + 3m_{\mathrm{m}}) + 4(8\bar{m} - 4 - \bar{m}\Delta m - 4b_{\mathrm{m}} + 4b_{\mathrm{m}}m_{\mathrm{m}} + b_{\mathrm{m}}\bar{m}(m_{\mathrm{f}} - 3m_{\mathrm{m}}) - 3M)n - 8\bar{m}(2 - \bar{m})n^2)) \quad \text{(S157)}$$

$$
\begin{aligned}
z_{\mathrm{MO}}{}'^{*} = {} & ((-2\Delta m(b_{\mathrm{f}}(H_{\mathrm{f}}+1)+b_{\mathrm{m}}(H_{\mathrm{m}}+1))(1-\overline{m})-\Delta m(b_{\mathrm{f}}(-10+2m_{\mathrm{f}}{}^{3} \\
& +m_{\mathrm{f}}(H_{\mathrm{m}}-6m_{\mathrm{m}}+16)-3m_{\mathrm{f}}{}^{2}(3-m_{\mathrm{m}})-H_{\mathrm{m}}+4m_{\mathrm{m}})+b_{\mathrm{m}}(-2 \\
& -m_{\mathrm{f}}{}^{2}(1-m_{\mathrm{m}})+H_{\mathrm{m}}(-5+2m_{\mathrm{m}})+m_{\mathrm{f}}(3H_{\mathrm{m}}-2m_{\mathrm{m}}+4)))n \\
& +(b_{\mathrm{f}}(8+m_{\mathrm{f}}{}^{4}+m_{\mathrm{f}}{}^{3}(m_{\mathrm{m}}-5)+(H_{\mathrm{m}}-3m_{\mathrm{m}}+4)m_{\mathrm{m}}-m_{\mathrm{f}}(m_{\mathrm{m}} \\
& -3)(-4+H_{\mathrm{m}})-m_{\mathrm{f}}{}^{2}\,(-11+m_{\mathrm{m}}+m_{\mathrm{m}}{}^{2}))+b_{\mathrm{m}}(8+m_{\mathrm{f}}{}^{3}(m_{\mathrm{m}} \\
& -1)+m_{\mathrm{f}}{}^{2}\,(3+H_{\mathrm{m}}-3m_{\mathrm{m}})-m_{\mathrm{m}}(4+m_{\mathrm{m}}(5+H_{\mathrm{m}}-3m_{\mathrm{m}})) \\
& +m_{\mathrm{f}}(-4+m_{\mathrm{m}}(6+m_{\mathrm{m}}-m_{\mathrm{m}}{}^{2}))))n^{2}))\;/\;((-4\Delta m(b_{\mathrm{f}}(H_{\mathrm{f}}+1) \\
& +b_{\mathrm{m}}(H_{\mathrm{m}}+1))(1-\overline{m})-2\Delta m(-4(M-2\overline{m}+1)(1-\overline{m}) \\
& +b_{\mathrm{f}}(-10+2m_{\mathrm{f}}{}^{3}+m_{\mathrm{f}}(H_{\mathrm{m}}-6m_{\mathrm{m}}+16)+3m_{\mathrm{f}}{}^{2}\,(m_{\mathrm{m}}-3)-H_{\mathrm{m}} \\
& +4m_{\mathrm{m}})+b_{\mathrm{m}}(-2-m_{\mathrm{f}}{}^{2}(1-m_{\mathrm{m}})+H_{\mathrm{m}}(-5+2m_{\mathrm{m}})+m_{\mathrm{f}}(3H_{\mathrm{m}} \\
& -2m_{\mathrm{m}}+4)))n+2(b_{\mathrm{f}}(8+m_{\mathrm{f}}{}^{4}+m_{\mathrm{f}}{}^{3}\,(m_{\mathrm{m}}-5)+(H_{\mathrm{m}}-3m_{\mathrm{m}} \\
& +4)m_{\mathrm{m}}-m_{\mathrm{f}}(m_{\mathrm{m}}-3)(-4+H_{\mathrm{m}})-m_{\mathrm{f}}{}^{2}\,(-11+m_{\mathrm{m}}+m_{\mathrm{m}}{}^{2})) \\
& +2(8+m_{\mathrm{f}}{}^{2}\,(4-3m_{\mathrm{m}})-m_{\mathrm{f}}{}^{3}(1-m_{\mathrm{m}})+m_{\mathrm{m}}(-6+H_{\mathrm{m}})-m_{\mathrm{f}}(10 \\
& +m_{\mathrm{m}}(-6+H_{\mathrm{m}}-m_{\mathrm{m}})))+b_{\mathrm{m}}(8-m_{\mathrm{f}}{}^{3}\,(1-m_{\mathrm{m}})+m_{\mathrm{f}}{}^{2}\,(3+H_{\mathrm{m}} \\
& -3m_{\mathrm{m}})-m_{\mathrm{m}}(4+m_{\mathrm{m}}(5+H_{\mathrm{m}}-3m_{\mathrm{m}}))+m_{\mathrm{f}}(-4+m_{\mathrm{m}}(6+m_{\mathrm{m}} \\
& -m_{\mathrm{m}}{}^{2}))))n^{2}+16\overline{m}(2-\overline{m})n^{3}))
\end{aligned}
\tag{S158}
$$

$$
\begin{aligned}
z_{\mathrm{MD}}{}'^{*} = {} & ((-2b_{\mathrm{f}}\Delta m(H_{\mathrm{f}}+1)(1-\overline{m})-\Delta m(-10+2m_{\mathrm{f}}{}^{3}+m_{\mathrm{f}}(H_{\mathrm{m}}-6m_{\mathrm{m}}+16) \\
& -3m_{\mathrm{f}}{}^{2}(3-m_{\mathrm{m}})-H_{\mathrm{m}}+4m_{\mathrm{m}})n+(8+m_{\mathrm{f}}{}^{4}+m_{\mathrm{f}}{}^{3}(m_{\mathrm{m}}-5) \\
& +(H_{\mathrm{m}}-3m_{\mathrm{m}}+4)m_{\mathrm{m}}+m_{\mathrm{f}}(3-m_{\mathrm{m}})(-4+H_{\mathrm{m}})-m_{\mathrm{f}}{}^{2}\,(m_{\mathrm{m}}-11 \\
& +m_{\mathrm{m}}{}^{2}))n^{2}))\;/\;((-4b_{\mathrm{f}}\Delta m(H_{\mathrm{f}}+1)(1-\overline{m})-2\Delta m(-2(M-2\overline{m} \\
& +1)(1-\overline{m})+b_{\mathrm{f}}(-10+2m_{\mathrm{f}}{}^{3}+m_{\mathrm{f}}(H_{\mathrm{m}}-6m_{\mathrm{m}}+16)+3m_{\mathrm{f}}{}^{2}(m_{\mathrm{m}} \\
& -3)-H_{\mathrm{m}}+4m_{\mathrm{m}}))n+2(8+m_{\mathrm{f}}{}^{2}(4-3m_{\mathrm{m}})-m_{\mathrm{f}}{}^{3}(1-m_{\mathrm{m}}) \\
& +m_{\mathrm{m}}(-6+H_{\mathrm{m}})-m_{\mathrm{f}}(10+m_{\mathrm{m}}(-6+H_{\mathrm{m}}-m_{\mathrm{m}}))+b_{\mathrm{f}}(8+m_{\mathrm{f}}{}^{4} \\
& +m_{\mathrm{f}}{}^{3}(m_{\mathrm{m}}-5)+(H_{\mathrm{m}}-3m_{\mathrm{m}}+4)m_{\mathrm{m}}-(M-3m_{\mathrm{f}})(-4+H_{\mathrm{m}}) \\
& -m_{\mathrm{f}}{}^{2}(-11+m_{\mathrm{m}}+m_{\mathrm{m}}{}^{2})))n^{2}+8\overline{m}(2-\overline{m})n^{3}))
\end{aligned}
\tag{S159}
$$

$$
\begin{aligned}
z_{\mathrm{MS}}{}'^{*} = ((&b_{\mathrm{m}}(2\Delta m(H_{\mathrm{m}}+1)(1-\overline{m}) - \Delta m(-2 - m_{\mathrm{f}}{}^{2}(1-m_{\mathrm{m}}) + H_{\mathrm{m}}(2m_{\mathrm{m}}-5) \\
&+ m_{\mathrm{f}}(3H_{\mathrm{m}} - 2m_{\mathrm{m}} + 4))n + (8 + m_{\mathrm{f}}{}^{3}(m_{\mathrm{m}}-1) + m_{\mathrm{f}}{}^{2}\,(3 + H_{\mathrm{m}} \\
&- 3m_{\mathrm{m}}) - m_{\mathrm{m}}(4 + m_{\mathrm{m}}(5 + H_{\mathrm{m}} - 3m_{\mathrm{m}})) + m_{\mathrm{f}}(-4 + m_{\mathrm{m}}(6 + m_{\mathrm{m}} \\
&- m_{\mathrm{m}}{}^{2})))n^{2}))) \;/\; ((-4b_{\mathrm{m}}\Delta m(H_{\mathrm{m}}+1)(1-\overline{m}) - 4\Delta m(-(M - 2\overline{m} \\
&+ 1)(1-\overline{m}) + b_{\mathrm{m}}(-2 - m_{\mathrm{f}}{}^{2}(1-m_{\mathrm{m}}) + H_{\mathrm{m}}(-5 + 2m_{\mathrm{m}}) + m_{\mathrm{f}}(H_{\mathrm{m}} \\
&- 6m_{\mathrm{m}} + 4)))n + 2(8 + m_{\mathrm{f}}{}^{2}\,(4 - 3m_{\mathrm{m}}) - m_{\mathrm{f}}{}^{3}(1-m_{\mathrm{m}}) + m_{\mathrm{m}}(-6 \\
&+ H_{\mathrm{m}}) - m_{\mathrm{f}}(10 + m_{\mathrm{m}}(-6 + H_{\mathrm{m}} - m_{\mathrm{m}})) + b_{\mathrm{m}}(8 - m_{\mathrm{f}}{}^{3}(1-m_{\mathrm{m}}) \\
&+ m_{\mathrm{f}}{}^{2}\,(3 + H_{\mathrm{m}} - 3m_{\mathrm{m}}) - m_{\mathrm{m}}(4 + m_{\mathrm{m}}(5 + H_{\mathrm{m}} - 3m_{\mathrm{m}})) + m_{\mathrm{f}}(-4 \\
&+ m_{\mathrm{m}}(6 + m_{\mathrm{m}} - m_{\mathrm{m}}{}^{2}))))n^{2} + 8\overline{m}(2-\overline{m})n^{3}))
\end{aligned} \qquad (S160)
$$

$$
\begin{aligned}
z_{\mathrm{FO}}{}'^{*} = ((&-(b_{\mathrm{f}}(H_{\mathrm{f}}+1) + 2b_{\mathrm{m}}\Delta m(H_{\mathrm{m}}+1))(1-\overline{m}) - \Delta m(b_{\mathrm{f}}(-2 + H_{\mathrm{f}}(-5 + 2m_{\mathrm{f}}) \\
&+ 4m_{\mathrm{m}} + m_{\mathrm{f}}(3m_{\mathrm{f}} - 8)m_{\mathrm{m}} - (1 - m_{\mathrm{f}})m_{\mathrm{m}}{}^{2}) + b_{\mathrm{m}}(-10 + 6m_{\mathrm{f}} \\
&- m_{\mathrm{f}}{}^{2}(H_{\mathrm{f}} - 6m_{\mathrm{f}} + 16)m_{\mathrm{m}} + 3(m_{\mathrm{f}} - 3)m_{\mathrm{m}}{}^{2} + 2m_{\mathrm{m}}{}^{3}))n + (b_{\mathrm{f}}(-8 \\
&+ m_{\mathrm{f}}{}^{4} + m_{\mathrm{f}}{}^{3}\,(m_{\mathrm{m}} - 5) + m_{\mathrm{m}}(4 + H_{\mathrm{m}} - m_{\mathrm{m}}) - m_{\mathrm{f}}(-4 + (m_{\mathrm{m}} \\
&- 3)H_{\mathrm{m}}) - m_{\mathrm{f}}{}^{2}(m_{\mathrm{m}} - 5 + m_{\mathrm{m}}{}^{2})) + b_{\mathrm{m}}(-8 - m_{\mathrm{f}}{}^{3}(1 - m_{\mathrm{m}}) \\
&+ m_{\mathrm{f}}{}^{2}\,(5 + H_{\mathrm{m}} - 3m_{\mathrm{m}}) - m_{\mathrm{m}}(-12 + m_{\mathrm{m}}(11 + H_{\mathrm{m}} - 3m_{\mathrm{m}})) \\
&+ m_{\mathrm{f}}(-4 + m_{\mathrm{m}}(2 + m_{\mathrm{m}} - m_{\mathrm{m}}{}^{2}))))n^{2})) \;/\; ((-4\Delta m(b_{\mathrm{f}}(H_{\mathrm{f}}+1) \\
&+ b_{\mathrm{m}}(H_{\mathrm{m}}+1))(1-\overline{m}) - 2\Delta m(-4(M - 2\overline{m} + 1)(1-\overline{m}) + b_{\mathrm{f}}(-2 \\
&+ H_{\mathrm{f}}(2m_{\mathrm{f}} - 5) + 4m_{\mathrm{m}} + M(3m_{\mathrm{f}} - 8) - (1 - m_{\mathrm{f}})m_{\mathrm{m}}{}^{2}) + b_{\mathrm{m}}(-10 \\
&+ 6m_{\mathrm{f}} - m_{\mathrm{f}}{}^{2}(H_{\mathrm{f}} - 6m_{\mathrm{f}} + 16)m_{\mathrm{m}} + 3(m_{\mathrm{f}} - 3)m_{\mathrm{m}}{}^{2} + 2m_{\mathrm{m}}{}^{3}))n \\
&+ 2(-16 - 16\Delta b + 12m_{\mathrm{f}} + 4b_{\mathrm{f}}m_{\mathrm{f}} - 4b_{\mathrm{m}}m_{\mathrm{f}} + 4m_{\mathrm{f}}{}^{2} + 5b_{\mathrm{f}}m_{\mathrm{f}}{}^{2} \\
&+ 5b_{\mathrm{m}}m_{\mathrm{f}}{}^{2} - 2m_{\mathrm{f}}{}^{3} - 5b_{\mathrm{f}}m_{\mathrm{f}}{}^{3} - b_{\mathrm{m}}m_{\mathrm{f}}{}^{3} + b_{\mathrm{f}}m_{\mathrm{f}}{}^{4} + (4(5 + b_{\mathrm{f}} + 3b_{\mathrm{m}}) \\
&+ 2(-6 - 2b_{\mathrm{f}} - \Delta b)m_{\mathrm{f}} - (6 + 2\overline{b} + 4b_{\mathrm{m}})m_{\mathrm{f}}{}^{2} + (2 + b_{\mathrm{f}} \\
&+ b_{\mathrm{m}})m_{\mathrm{f}}{}^{3})m_{\mathrm{m}} + (-8 + 6m_{\mathrm{f}} - b_{\mathrm{f}}(3 + H_{\mathrm{f}} - 3m_{\mathrm{f}}) + b_{\mathrm{m}}(-11 + m_{\mathrm{f}} \\
&+ m_{\mathrm{f}}{}^{2}))m_{\mathrm{m}}{}^{2} + (2 + b_{\mathrm{f}} + 5b_{\mathrm{m}} - 2(1 + \overline{b})m_{\mathrm{f}})m_{\mathrm{m}}{}^{3} - b_{\mathrm{m}}m_{\mathrm{m}}{}^{4})n^{2} \\
&- 16\overline{m}(2-\overline{m})n^{3}))
\end{aligned} \qquad (S161)
$$

$$
\begin{aligned}
z_{\mathrm{FD}}'^{*} = ((&-2b_{\mathrm{f}}\Delta m(H_{\mathrm{f}}+1)(1-\bar{m}) - \Delta m(-2 + H_{\mathrm{f}}(2m_{\mathrm{f}}-5) + 4m_{\mathrm{m}} \\
&+ m_{\mathrm{f}}(3m_{\mathrm{f}}-8)m_{\mathrm{m}} - (1-m_{\mathrm{f}})m_{\mathrm{m}}^{2})n + (-8 + m_{\mathrm{f}}^{4} + m_{\mathrm{f}}^{3}\,(m_{\mathrm{m}} \\
&- 5) + m_{\mathrm{m}}(4 + H_{\mathrm{m}} - m_{\mathrm{m}}) - m_{\mathrm{f}}(-4 - H_{\mathrm{m}}(3-m_{\mathrm{m}})) - m_{\mathrm{f}}^{2}\,(m_{\mathrm{m}} \\
&- 5 + m_{\mathrm{m}}^{2}))n^{2}))) \;/\; ((2(-2b_{\mathrm{f}}\Delta m(H_{\mathrm{f}}+1)(1-\bar{m}) - \Delta m(-2(M \\
&- 2\bar{m} + 1)(1-\bar{m}) + b_{\mathrm{f}}(-2 + H_{\mathrm{f}}(2m_{\mathrm{f}}-5) + 4m_{\mathrm{m}} \\
&+ m_{\mathrm{f}}(3m_{\mathrm{f}}-8)m_{\mathrm{m}} - (1-m_{\mathrm{f}})m_{\mathrm{m}}^{2}))n + (-8 + m_{\mathrm{f}}^{2}(2-3m_{\mathrm{m}}) \\
&- m_{\mathrm{f}}^{3}(1-m_{\mathrm{m}}) + m_{\mathrm{m}}(10 + H_{\mathrm{m}} - 2m_{\mathrm{m}}\,) - m_{\mathrm{f}}(-6 + m_{\mathrm{m}}(6 + H_{\mathrm{m}} \\
&- m_{\mathrm{m}})) + b_{\mathrm{f}}(-8 + m_{\mathrm{f}}^{4} + m_{\mathrm{f}}^{3}(m_{\mathrm{m}} - 5) + m_{\mathrm{m}}(4 + H_{\mathrm{m}} - m_{\mathrm{m}}) \\
&- m_{\mathrm{f}}(-4 + H_{\mathrm{m}}(3-m_{\mathrm{m}})) - m_{\mathrm{f}}^{2}\,(m_{\mathrm{m}} - 5 + m_{\mathrm{m}}^{2})))n^{2} \\
&- 4\bar{m}(2-\bar{m})n^{3})))
\end{aligned}
\tag{S162}
$$

$$
\begin{aligned}
z_{\mathrm{FS}}'^{*} = ((&b_{\mathrm{m}}(2\Delta m(H_{\mathrm{f}}+1)\,(1-\bar{m}) - \Delta m(-10 + 6m_{\mathrm{f}} - m_{\mathrm{f}}M(H_{\mathrm{f}} - 6m_{\mathrm{f}} + 16) \\
&+ 3(-3+m_{\mathrm{f}})\,m_{\mathrm{m}}^{2} + 2m_{\mathrm{m}}^{3})n \;+ (-8 - m_{\mathrm{f}}^{3}(1-m_{\mathrm{m}}) + m_{\mathrm{f}}^{2}(5 \\
&+ H_{\mathrm{m}} - 3m_{\mathrm{m}}) - m_{\mathrm{m}}(-12 + m_{\mathrm{m}}(11 + H_{\mathrm{m}} - 3m_{\mathrm{m}})) + m_{\mathrm{f}}(-4 \\
&+ m_{\mathrm{m}}(2 + m_{\mathrm{m}} - m_{\mathrm{m}}^{2})))n^{2}))) \;/\; ((2(-2b_{\mathrm{m}}\Delta m(H_{\mathrm{m}}+1)(1-\bar{m}) \\
&- \Delta m(-2(M - 2\bar{m} + 1)(1-\bar{m}) + b_{\mathrm{m}}(-10 + 6m_{\mathrm{f}} - m_{\mathrm{f}}M(H_{\mathrm{f}} \\
&- 6m_{\mathrm{f}} + 16) + 3(-3+m_{\mathrm{f}})\,m_{\mathrm{m}}^{2} + 2m_{\mathrm{m}}^{3}))n + (-8 \\
&+ m_{\mathrm{f}}^{2}(2-3m_{\mathrm{m}}) - m_{\mathrm{f}}^{3}\,(1-m_{\mathrm{m}}) + m_{\mathrm{m}}(10 + H_{\mathrm{m}} - 2m_{\mathrm{m}}) \\
&- m_{\mathrm{f}}(-6 + m_{\mathrm{m}}(6 + H_{\mathrm{m}} - m_{\mathrm{m}})) + b_{\mathrm{m}}(-8 - m_{\mathrm{f}}^{3}(1-m_{\mathrm{m}}) + m_{\mathrm{f}}^{2}(5 \\
&+ H_{\mathrm{m}} - 3m_{\mathrm{m}}) - m_{\mathrm{m}}(-12 + m_{\mathrm{m}}(11 + H_{\mathrm{m}} - 3m_{\mathrm{m}})) + m_{\mathrm{f}}(-4 \\
&+ m_{\mathrm{m}}(2 + m_{\mathrm{m}} - m_{\mathrm{m}}^{2}))))n^{2} - 4\bar{m}(2-\bar{m})n^{3})))
\end{aligned}
\tag{S163}
$$

where $\Delta m = m_{\mathrm{f}} - m_{\mathrm{m}}$, $\bar{m} = (m_{\mathrm{f}} + m_{\mathrm{m}})/2$, $M = m_{\mathrm{f}}m_{\mathrm{m}}$, $\Delta b = b_{\mathrm{f}} - b_{\mathrm{m}}$, $\bar{b} = (b_{\mathrm{f}} + b_{\mathrm{m}})/2$, $H_{\mathrm{f}} = (m_{\mathrm{f}} - 2)m_{\mathrm{f}}$, $H_{\mathrm{m}} = (m_{\mathrm{m}} - 2)m_{\mathrm{m}}$.

Here we show what if there are differences between the parental genetic effects on daughters and those on sons in the context of between-group combats, hence left-handedness is marginally altruistic. Under female-biased dispersal, genes carried by parents would favour a lower level of left-handedness for daughters than for sons; while under male-biased dispersal, genes carried by parent would favour a higher level of left-handedness for daughters than for sons (Figure S2).
